# Molecular programs of regional specification and neural stem cell fate progression in developing macaque telencephalon

**DOI:** 10.1101/2022.10.18.512724

**Authors:** Nicola Micali, Shaojie Ma, Mingfeng Li, Suel-Kee Kim, Xoel Mato-Blanco, Suvimal Sindhu, Jon I. Arellano, Tianliuyun Gao, Alvaro Duque, Gabriel Santpere, Nenad Sestan, Pasko Rakic

## Abstract

Early telencephalic development involves patterning of the distinct regions and fate specification of the neural stem cells (NSCs). These processes, mainly characterized in rodents, remain elusive in primates and thus our understanding of conserved and species-specific features. Here, we profiled 761,529 single-cell transcriptomes from multiple regions of the prenatal macaque telencephalon. We defined the molecular programs of the early organizing centers and their cross-talk with NSCs, finding primate-biased signaling active in the antero-ventral telencephalon. Regional transcriptomic divergences were evident at early states of neocortical NSC progression and in differentiated neurons and astrocytes, more than in intermediate transitions. Finally, we show that neuropsychiatric disease- and brain cancer-risk genes have putative early roles in the telencephalic organizers’ activity and across cortical NSC progression.

**One-Sentence Summary:** Single-cell transcriptomics reveals molecular logics of arealization and neural stem cell fate specification in developing macaque brain

## INTRODUCTION

The prenatal development of the telencephalon, including the cerebral cortex, involves genetically and epigenetically controlled cellular events that determine the patterning of the prospective domains by early organizing centers and the fate specification of the radial glia (RG) cells, which serve as neural stem cells (NSCs) in the germinal zones (*1-4*). The fate potential of the RG cells progresses throughout corticogenesis. They first generate waves of radially migrating glutamatergic excitatory neurons becoming later gliogenic, and producing oligodendrocytes and astrocytes (*5-7*).

Defining the genetic mechanisms that orchestrate the spatiotemporal progression of telecephalic and cortical NSCs in fetal pimates is crucial, for understanding both the formation of the distinct cell types and the emergence of higher-order cognitive abilities (*8*). Unprecedented advances in understanding the genetic architecture governing the lifespan development of human and non-human primate brain have been achieved in the last decade by bulk tissue transcriptomic and epigenomic profiles (*9-13*). More recently, single-cell multi-omics from midfetal and adult brains (*14-17*), as well as *in vitro* neural systems (*18, 19*), have shed light on the finest mechanisms orchestrating the formation of different regions of the telencephalon. However, we are still missing a clear view of the early fetal genetic events in primates that instruct the spatial identity of NSCs and drive the diversification of the telecephalic regions and cortical areas, as cells traverse neuronal and glial trajectories.

These mechanisms governing the regional patterning of the telencephalon and cortex have been better characterized in rodents (*20*). However, our understanding of the complex primates’ brain might be limited in these animal models and it is challenging to explore these events in post-mortem early fetal human samples (*21*). Non-human primates (NHP), and especially rhesus macaque, represent a valid model to systematically dissect the molecular programs underlying the formation of the primates’ brain and translate the findings in humans. It is also essential to compare NHP prenatal transcriptomic datasets with the other genomic resources which are being generated across the development of multiple species (*14, 22*), to identify the fundamental mechanisms underlying brain evolution, the emergence of specialized cell subtypes and their link with neuropsychiatric disorders in humans (*11, 23, 24*).

Here, we profiled over 761,000 cells across multiple regions of the developing rhesus monkey telencephalon by single-cell RNA-sequencing (scRNA-seq), from the early phases when the telencephalic organizers pattern different regional anlage before the onset of neurogenesis, till mid-gliogenesis. We defined the molecular programs driving the spatial identity of the early NSCs and the specification of cortical neurons and glia, which both further diversify across the regions at more mature states. Finally, we show that brain disease-associated genes, including neuropsychiatric disorders and cancers, might have early roles in the brain organizers and across the progression of region-specific NSCs. This work sheds light on the genetic mechanisms of primate neocortical arealization and underlines early NSC events as a risk for human brain disorders.

## RESULTS

### Spatiotemporal transcriptomic characterization of prenatal macaque telencephalic cells

Employing timed-pregnant rhesus macaques, we dissected multiple prospective regions of the prenatal telencephalon, from embryonic day (E)37, prior to neurogenesis, up to mid-gliogenesis at E110 (*12*) (Fig. 1A). Anterior (A)/frontal (FR), dorso-lateral (DL)/putative motor-somatosensory (MS), putative temporal, posterior (P)/occipital (OC) areas, and ganglionic eminence (GE) were recognized at the earliest stages (E37-E78). Then, at the latest stages (E93-E110), we collected the cortical wall of up to 13 regions of the neocortex, including multiple sub-areas from the prefrontal (dorsolateral: DFC; orbital: OFC; medial: MFC; ventrolateral: VFC), primary motor (M1C), parietal (primary somatosensory: S1C; inferior parietal: IPC; posterior cingulate: PCC), insula (Ins), temporal (inferior: ITC; superior: STC; primary auditory: A1C), occipital (primary visual: V1), and three regions from GE (medial: MGE; lateral: LGE; caudal: CGE) (*24*). Eighty-three samples categorized were processed for scRNA-seq (fig. S1A). After stringent quality control, a total of 761,529 high-quality individual cells were obtained (fig. S1B and table S1).

**Fig. 1.**
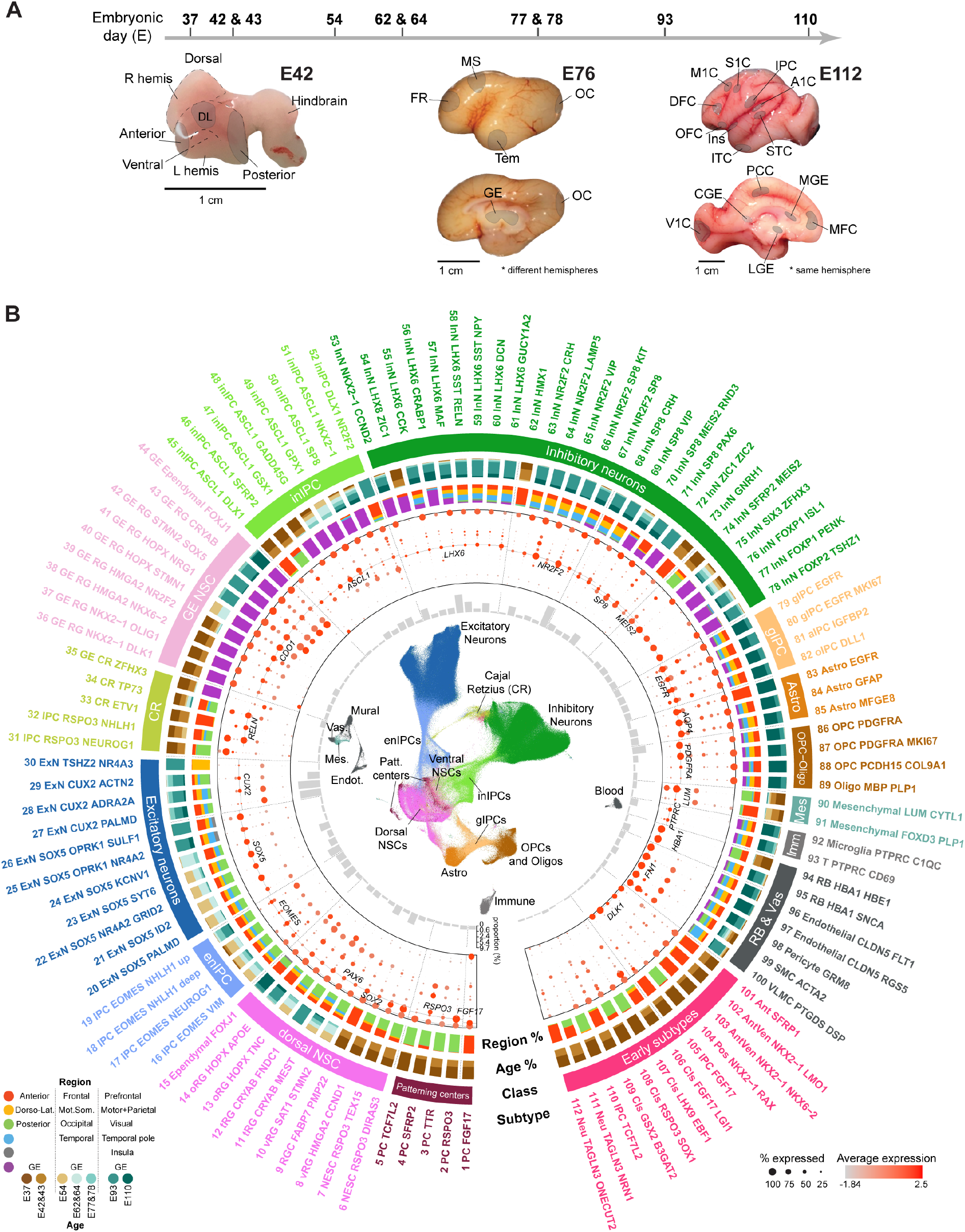
Cell atlas of macaque fetal telencephalon. (**A**) Scheme of the gestational time points (E) across macaque fetal development relevant for this study. Three representative images of E42, E76 and E112 macaque brains showing the areas dissected. Scale bar: 1 cm. R/L hemis: right and left hemispheres.(**B**) Transcriptomic taxonomy of monkey developing telencephalic cells. From innermost to outmost: uniform manifold approximation and projection (UMAP) plot visualizing cells colored by classes; subtype proportions; cell class marker expression; region and age proportion; cell class; cell subtypes. The atlas includes 15 cell classes and 112 cell subtypes. The nomenclature of the regions is based on the maturation of the telencephalon across the embryonic age (the bottom left). NSC: neural stem cells; RGC: radial glial cells; IPC: intermediate precursors cells; enIPC: excitatory neuron IPC; ExN: excitatory neurons; CR: Cajal-Retzius neurons; GE: gangionic eminence; inIPC: inhibitory neuron IPC; InN: inhibitory neurons; gIPC: glial IPC; aIPC: astrocyte IPC; Astro: astrocytes; Oligo: oligodendrocyte; OPC: oligodendrocyte precursor cells; Mes: mesenchymal cells; Imm: immune cells; RB and Vas: red blood lineage and vascular cells.

Based on unsupervised clustering and marker gene expression, 112 transcriptomically-defined telencephalic cell subtypes (hereafter referred to as “subtypes”) were identified, representing major cell classes and lineages that overlaid with age and region of origin (Fig. 1B, fig. S2, A and B). Subtypes included early progenitors of putative patterning centers; dorsal and ventral neural stem cells (NSCs) traversing glutamatergic excitatory or GABAergic inhibitory neuronal trajectories, respectively; astrocyte and oligodendrocyte lineages; Cajal-Retzius neuron lineage; non-neural cells (immune, mesenchymal, endothelial, mural and blood cells). These cell types showed a temporal and topographic continuum consistent with the spatiotemporal information of the samples (fig. S2, C-E). These data present a comprehensive single-cell spatiotemporal transcriptomic survey of the macaque developing telencephalon, enabling a more detailed investigation of the underlying molecular and cellular features. This resource is also interactively accessible at: http://resources.sestanlab.org/devmacaquebrain.

### Transcriptomic signatures of macaque telencephalic patterning centers

The events governing the regional patterning of the telencephalon occur early in development and involve progenitors of the patterning centers (PC, also called telencephalic organizers) (*3, 4, 25*). Through marker gene expression, transcriptomic integration with published datasets and RNAscope validation (Fig. 2, A-C, and fig. S3, A-D), we identified several domain-specific progenitor subtypes representing putative telencephalic organizers (hereafter referred to “organizer domain subtypes”). Three co-clustering subtypes, detected in the early anterior region of the monkey developing telencephalon, showed expression of well-established antero-ventral NSC genes (*FGF17, FGF18, FGF8, SP8, FOXG1, NKX2-1* and *SHH*) representing rostral patterning center (RPC PC *FGF17*) (*26-28*) and putative antero-ventral (AV) organizer progenitors (AV *NKX2-1/NKX6-2* and AV *NKX2-1/LMO1*) (*28*) (Fig. 2, A and B; table S2). While some of these genes have been previously identified as expressed in bulk-tissue samples of developing frontal human and primate cortical walls (*9-11, 29*), cell type-specificity of their expression pattern has not been resolved. We found ZIC genes (*ZIC1, ZIC3* and *ZIC4*) expressed in the early anterior subtypes. Then by RNAscope, we further confirmed their expression in the ventricular zone (VZ) of the antero-ventral domain of E40 monkey telencephalon, where they form a ventro-dorsal and antero-posterior gradient along with *SP8, NKX2-1, FOXG1* and *FGF18* in the same region (Figs. 2, B and C, and 3Bii, fig. S3B). Two subtypes in the early medio-posterior region were characterized by features of the cortical hem (PC *RSPO3* and PC *TTR*), the dorso-caudal organizer of the telencephalon, expressing WNT and BMP signaling members (*RSPO3, WNT8B* and *BAMBI*) and *LMX1A* (Fig. 2B and fig. S3B; table S2) (*30, 31*). These cells also expressed *ARX, FGFR3* and *LHX9*, displaying a posterior-anterior gradient (Fig. 2, B and C, figs. S3B and S4D). One subtype unique to the medio-posterior region, expressing *IRX3, WNT3* and *WNT4*, was reminiscent of the zona limitans intrathalamica (ZLI) (*32*). In addition, one putative antihem subtype was also identified, characterized by *SFRP2, PAX6, NR2F1* and *NR2F2* expression (*33*) (Fig. 2B). Finally, two early cell subtypes were found in the GE (*NKX2-1/DLK1* and *NKX2-1/OLIG1*). They co-clustered with the anterior-ventral *NKX2-1*^+^ organizer domain subtypes and expressed the SHH regulator *NKX2-1, FOXG1* and *GSX2*, likely representing the organizer of the ventral forebrain (*4, 25*) (Fig. 2, A-C, fig. S3, A and B). Other markers of these GE subtypes, such as *MEIS2*, involved in retinoic acid (RA) response in the late developing prefrontal cortex (*34*), and its co-transcription factors *PBX1 and PBX3*, known to pattern the mouse telencephalon (*35*), were detected in this early antero-ventral domain by RNAscope, forming a ventro-dorsal gradient in the NSCs (Fig. 2B and fig. S3B). Of note, all these organizer domain subtypes were transient and barely detectable in the samples collected after E43 (Fig. 2A) (*22*). These putative telencephalic organizers were further validated by transcriptomic comparisons with a mouse scRNA-seq dataset and a macaque bulk exon-array dataset of developing brain (*12, 22*) (fig. S3, C and D). Other early subtypes co-clustering with these organizer domain subtypes, including Cls *FG17-LGI1*, Cls *LHX9-EBF1*, Cls *RSPO3-SOX1* and Cls *GSX2-B3GAT2*, were small in size and their identities were left as unknown (fig. S3A).

**Fig. 2.**
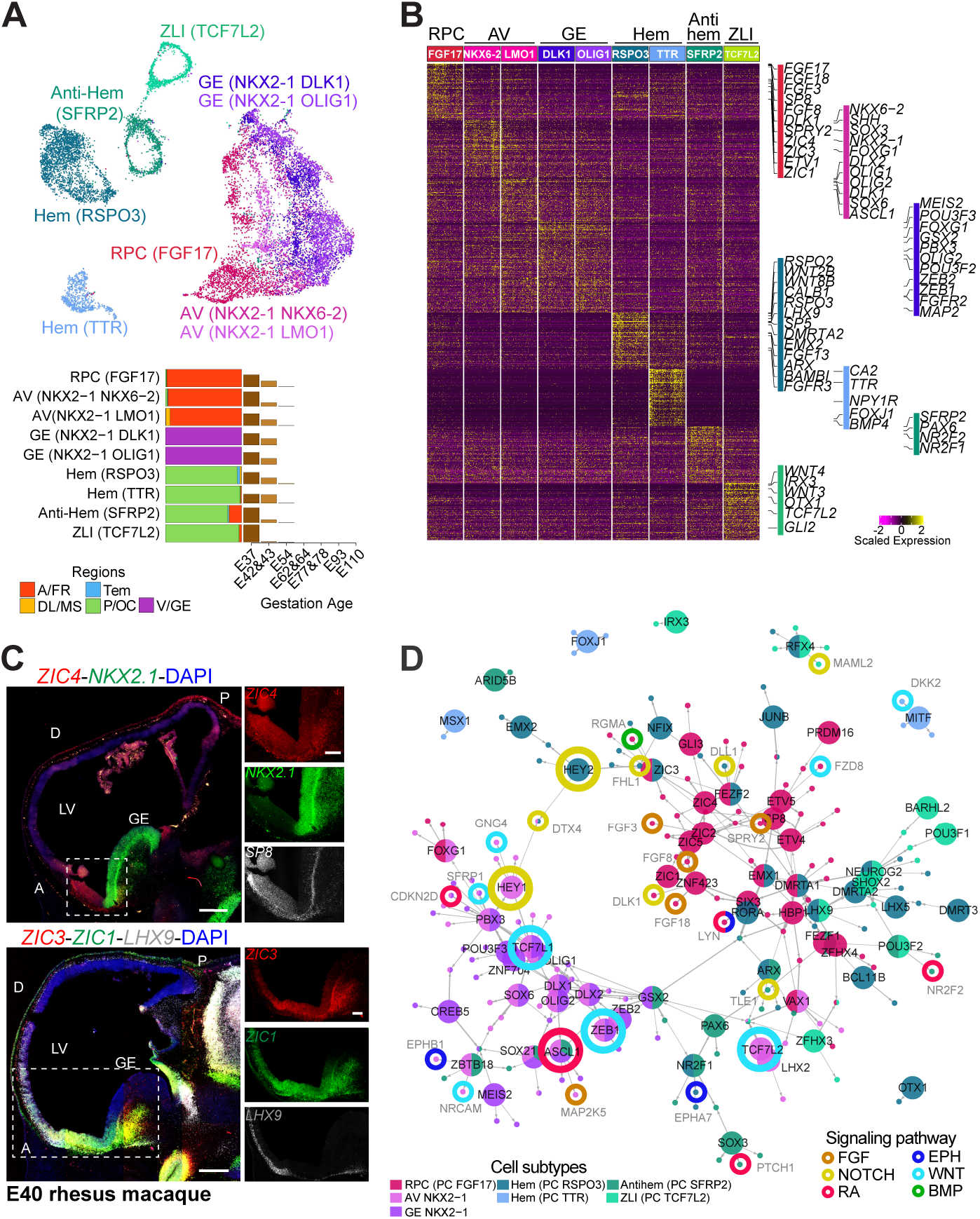
Molecular signatures of putative telencephalic organizer domains. (**A**) Visualization of putative organizer centers cell subtypes (top) on the UMAP layout, and region and age compositions (bottom). (**B**) Expression of subtype markers for the indicated subtypes. (**C**) RNAscope analysis of *ZIC4, SP8* and *NKX2-1* (top), *ZIC3, ZIC1* and *LHX9* (bottom) expression in E40 macaque sagittal brain sections. Scale bars: 500 µm (panoramic view) and 200 µm (zoom-in view), (**D**) Predicted regulatory network of transcription factors and targets. Each node represents a gene. Big and small dots denote transcriptor factors and their targets, respectively. Nodes are colored by the cell subtypes where they are predicted to have a regulatory role. Colored rings indicate annotated signaling pathways a gene belong to. A: Anterior; P: posterior; AV: antero-ventral domain; D: dorsal; V: ventral; LV: lateral ventricle.

Brain organizers secrete signaling morphogens that instruct gene and transcription factor (TF) expression gradients across the developing telencephalon (*1, 4, 27, 31, 36, 37*). By integrating motif enrichment and gene-gene coexpression information, we built a network connecting TFs with their putative target genes across different telencephalic organizers, highlighting shared and domain-specific transcriptional regulations (Fig. 2D). The rostral pattern center subtype (*FGF17*^+^) largely employed unique TF-target regulations involving *ZIC1-5*, upstream of FGF pathway-related genes such as *FGF3, FGF8, FGF18* and *SPRY2* and the fate regulator *SP8* (Fig. 2D). The *NKX2-1*^*+*^ subtypes of the antero-ventral domain and the ganglionic eminence, exhibited substantial overlap of regulations, including *OLIG2* upstream of the key interneuron gene *ASCL1* (Fig. 2D). Similarly, the organizer domain subtypes in the posterior regions of the telencephalon showed many overlapping TFs. *ARX* and *NR2F1* were shared by cortical hem and antihem; *NEUROG2* and *LHX9* were shared by cortical hem and zona limitans. These data provide a view of the putative TFs which integrate the patterning action of the organizer centers. Notably, different elements of the same signaling pathways were divergently activated in distinct domains (Fig. 2D). For example, the NOTCH signaling genes *DLK1* and *HEY1* were recruited by the antero-ventral domain, while *HEY2* and *DLL1* of the same signaling pathway were activated in posterior domains. Similarly, the WNT signaling members *TCF7L1* and *SFRP1* were found in the antero-ventral domain, in contrast to *DKK2* active in posterior domain. Together, these data denote the combinatorial interaction of TFs and signaling molecules orchestrating telencephalic organizer function to instruct regional cell identities.

To identify potential progeny neurons from these early progenitors, we employed RNA velocity and traced their differentiation trajectory towards diverse neuron subtypes (materials and methods). The anterior-ventral progenitors *NKX2-1*^+^ and *FGF17*^+^ generated a heterogeneous pool of interneurons, including an *LHX8*^+^ and an *ONECUT1*^*+*^*/ONECUT2*^+^ population, respectively. Cortical hem progenitors transitioned into *TP73*^+^ IPCs, eventually generating *CALB2*^+^/*RELN*^+^ Cajal-Retzius neurons (fig. S3E). Of note, the majority of these cells were barely datectable after E43, suggesting they might be transient (fig. S3A). These results define the origins of the earliest neurons generated by the organizer centers of the telencephalon and their transcriptional dynamics across the differentiation trajectories. Taken together, these data highlight the molecular events that control macaque telencephalic organizer activities.

### Signaling interaction between brain organizers and telencephalic radial glial cells

To define how the regional identity of the radial glia (RG) cells is instructed by the telencephalic organizers, we leveraged annotated ligand-receptor pairs databases and inferred cell-cell interactions between organizer domain subtypes (AV [*NKX2-1*^+^], RPC [*FGF17*^+^] and cortical hem [*RSPO3*^+^ and *TTR*^+^]) expressing the ligands and frontal, occipital and ventral RG cells expressing the receptors (materials and methods). The ligand-receptor pairs of diverse signaling pathways were then organized into 10 modules, reflecting shared or region-specific interaction types (Fig. 3A, fig. S4, A and B). For example, modules (M) M5 and M6, largely consisting of ligand-receptor pairs implicated in Ephrin and FGF signaling pathways, respectively, inferred signaling interactions between the anterior-ventral organizers (*NKX2-1*^+^ and *FGF17*^+^) and the RG cells (fig. S4C). M6 was characterized by *FGF18*-*FGFR1* and *FGF18*-*FGFR3* pairs, mediating the signal from the rostral patterning center progenitors expressing *FGF18* ligand towards frontal, occipital and ventral RG cells expressing the receptor *FGFR1*, or selectively towards occipital and ventral RG cells expressing *FGFR3* (Fig. 3Bi, fig. S4, C and D). This prediction was supported by RNAscope of E40 monkey brain tissue, which detected expression of *FGF18* in the antero-ventral domain of the telencephalon; *FGFR1* in the VZ of the antero-ventral region and ganglionic eminence, and with less extent in the posterior telencephalon; *FGFR3* in the caudo-ventral domain (Fig. 3B and fig. S4D). Modules M4 and M7, enriched in WNT and Notch signaling ligand-receptor pairs, respectively, inferred interactions between the cortical hem and RG cells (fig. S4C). Moreover, *WNT5A-FZD5* and *WNT5A-PTPRK* pairs, belonging to M2 and M3, respectively, predicted cross-talks from the cortical hem expressing *WNT5A* ligand towards ventral RG cells expressing Frizzled receptor *FZD5*, or towards frontal RG cells expressing the WNT signaling inhibitor Protein Tyrosine Phosphatase Receptor-type Kappa (*PTPRK*) (Fig. 3B and fig. S4C). RNAscope supported this spatial expression pattern, detecting expression of *WNT5A* by the medial-posterior cells, *FZD5* by ventral RG cells and *PTPRK* by dorsal RG cells (Fig. 3B). The domain of expression of other ligand-receptor pairs involved in Ephrin (*EFNA2*-*EPHA3*; M2) and WNT signaling (*RSPO2*-*LGR4*; M2) were further validated by RNAscope (fig. S4, C and D). These data strongly suggest that brain organizers selectively signal to competent RG cells. Moreover, the results show the regional expression of diffusible morphogens and paired receptors in macaques, denoting a signaling code integrating the cross-talks between telencephalic organizers and region-specific RG cells.

**Fig. 3.**
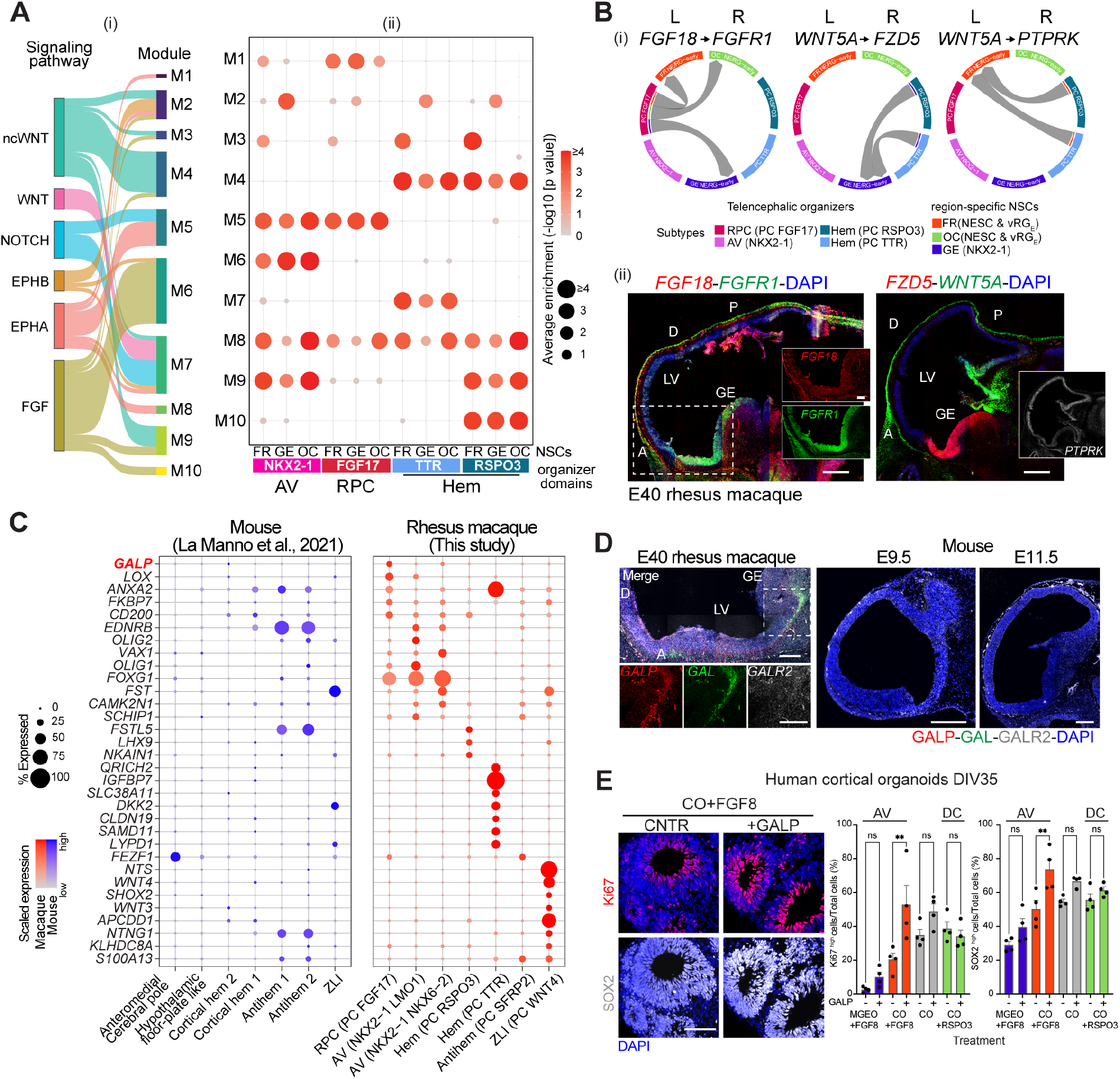
Signaling cross-talks between telencephalic organizers and radial glia cells. (**A**) (i) Sankey plot showing modules (M1-M10) of ligand (L)-receptor (R) pairs between organizer cell subtypes and RG cells for selected signaling pathways. The thickness of the links is proportional to the numbers of L-R pairs. (ii) Dot plot summarizing predicted interaction patterns between organizer cell subtypes and E37-43 NSC subtypes (NESCs and vRG_E_) of frontal (FR), ganglionic eminence (GE) and occipital (OC) regions. (**B**) Top (i): interaction patterns of selected L-R pairs. The arrows show the direction of the interaction. Bottom (ii): RNAscope of E40 macaque sagittal brain sections for L or R genes. (**C**) Genes enriched in macaque (this dataset) versus mouse subtypes identified as telencephalic organizers (*22*). (**D**) RNAscope of E40 macaque and E9.5 and E11.5 mouse sagittal brain sections for *GALP, GAL* and *GALR2*. (**E**) (Left) Immunohistochemistry for SOX2 and Ki67 in human cortical organoids exposed to FGF8 plus/minus GALP (complete panel in fig. S6E). Scale bar 100 µm. (Right) Percentage of cells expressing the indicated marker in the different conditions ± SD, One-way ANOVA, Dunnett’s multiple comparison (*: p < 0.1; **: p < 0.01; ***: p < 0.001; ****: p < 0.0001; ns: not significant). Scale bars: 500 µm (panoramic view) and 200 µm (zoom-in view).

### Putative primate-specific proliferation signaling from the antero-ventral domain

Transcriptomic comparisons between this macaque and a mouse developing brain single-cell RNAseq dataset (*22*) revealed that the telencephalic organizers of the two species share transcriptomic similarities (fig. S5A). However, notable species differences were also identified (Fig. 3C and fig. S5B). Genes enriched in macaque organizers included *ANXA2* and the insulin growth factor binding protein (*IGFBP7*) in the cortical hem, the neuropeptide Neurotensin (*NTS*) in the ZLI, several TFs and, interestingly, the neuropeptide *GALP* in the rostral patterning center (Fig. 3C). *GALP* gene encodes galanin-like peptide involved in hypothalamic functions in adult rodents (*38*). We found it expressed by the rostral patterning center cells and their progeny lineages (fig. S5C). However, *GALP* expression was dramatically reduced by E54, indicating its transient function at early telencephalic phases. RNAscope analysis validated this expression pattern. *GALP* and the family-related Galanin (*GAL*) mRNA were detected in the antero-ventral domain of the E40 monkey, but not in mouse telencephalon at the equivalent developmental ages E9.5 and E11.5. The GALP/GAL receptor 2 (*GALR2*) mRNA was detected in the monkey telencephalon and with less extent in the mouse (Fig. 3D).

To recapitulate early events of brain development *in vitro* (*18, 39*), we generated dorsal (hCO) and MGE (hMGEO) telencephalic organoids, from human induced pluripotent stem cells (hiPSCs). Organoids were directed towards a dorso-caudal or antero-ventral telencephalic identity by modulating RSPO3- and FGF8-induced patterning signaling, respectively. Treatment was done during their early differentiation (day in vitro, DIV8-21) when neuroepithelial cells form telencephalic organizer states (*40*). Then, organoids were cultured with the same basal differentiation medium and analyzed at DIV35. Standard telencephalic markers confirmed their region identity (fig. S6A), showing that organoids also recapitulated molecular features of the early domains of the fetal monkey telencephalon. These included a higher GAL and GALP protein expression in the antero-ventral MGEO versus the dorsal CO, where also we detected higher ZIC4 and SP8 protein levels (fig. S6B). GALR2 was abundant in the hMGEO, however also detectable in the other conditions. These data suggest that *ZIC4, GALP* and *GAL* expression are intrinsic features of neural cells with antero-ventral telencephalic identity. Moreover, as the expression of the regional markers was checked at DIV35, long after the ligand stimulation was finished (DIV21), this experiment suggests that the regional identity of the cells was maintained after the patterning action was terminated.

As Galanin has a role in the proliferation and differentiation of adult mouse hippocampal NSCs (*41*), we wondered whether exogenous *GALP* stimulation could change the proliferation and differentiation rate of the RG cells. We generated rhesus macaque iPSC-derived CO (rmCO) and exposed them to 30 ng/ml GALP, GAL or both ligands from DIV46 to 60 (fig. S6, C and D). GALP and GAL treated rmCO showed enhanced expression of Ki67, SOX2 and PAX6, and reduction of the neuron markers HuCD, CTIP2 and GABA, indicating increased proliferation of the NSCs and compromised neuronal differentiation. Finally, we checked whether GALP has an instructing role in a preferential telencephalic region. Both hCO and hMGEO exposed to FGF8, and hCO exposed to RSPO3, were treated with or without 30 ng/ml GALP during the patterning phase (DIV8 – 21) (Fig. 3E and fig. S6E). Exogenous GALP showed a significant effect on Ki67 and SOX2 expression in the hCO with an anterior identity (hCO+FGF8), more than the hMGEO (hMGEO+FGF8) and dorso-caudal hCO (hCO+RSPO3). Together, these data indicate that *GALP* increases the proliferation of RG cells, with a preference for those with an anterior identity.

### Transcriptomic variation of the neural stem cell progression across the telencephalic regions

Based on established marker gene expression, cortical NSCs were distinguished into subtypes showing different regional and age proportions (Fig. 4A, fig. S7, A and B). These included two NESC subtypes, two early (vRG_E_) and one late (vRG_L_) ventricular RG subtypes, two truncated (t)RG subtypes, one ependymal cell subtype, and two outer (o)RG cells. The appearance of these subtypes was correlated with the developmental ages (Fig. 4B). Moreover, pseudotime analysis further defined the continuous progression of the ventricular NSCs up to ependymal cells and distinguished the oRG cell lineage (Fig. 4A and fig. S7C). Transcriptomic comparisons between these data and human developing brain scRNA-seq datasets (*14, 42*) confirmed the identities of the RG subtypes we annotated. Moreover, extending the previous studies (*14*), this analysis defined earlier NSC states across the telencephalic regions (fig. S7D). While most of these subtypes were found in all the four main regions collected, i.e., frontal, motor-somatosensory, temporal and occipital cortical wall progressing across the time (E37-110), an early ventricular RG cell subtype (vRG_E_ *PMP22*^*+*^), marked by high expression of *CYP26A1, ZIC1, ZIC3, ZIC4* and *EFNA5*, was selectively enriched in the anterior region, representing a unique NSC population (Fig. 4B).

**Fig. 4.**
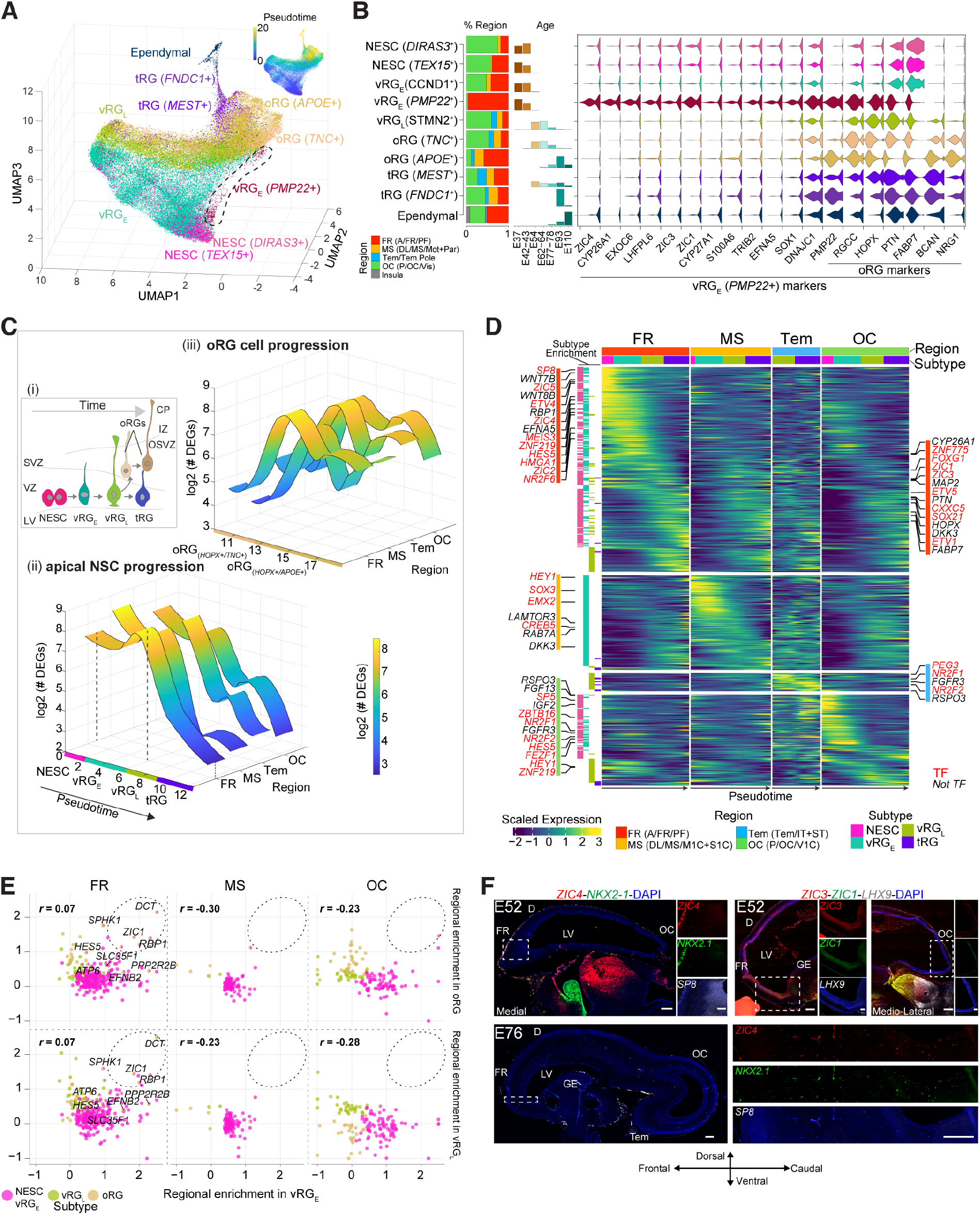
Transcriptomic variation of neural stem cell progression across telencephalic areas. (**A**) UMAP showing cortical NSC subtypes and pseudotime throughout age (top-right). The regions considered from each region follow the nomenclature across the time shown in Fig. 1. (**B**) Left: Region and age proportion of the NSCs. Right: Expression of subtype markers. (**C**) (i) Schema of the progression of the ventricular and subventricular (oRG) NSCs. (ii-iii) Number (in log2 scale) of differential expressed genes (DEGs) between regions along the pseudotime of VZ NSCs (ii) and (iii) oRG cells. (**D**) Region-specific gene expression cascades along the NSC progression. The legend indicates the regions considered across the time, as indicated in Fig. 1. (**E**) Correlation of region specificity of gene expression between early (NESCs and vRG_E_) and late (vRG_L_) NSCs or oRG cells. Coefficients are shown on the top-left. Genes are colored by the subtypes where they display regional enrichment. **(F)** RNAscope of macaque brain sagittal sections for the indicated genes at E52 (top) and E76 (bottom). Scale bars: 500 µm (panoramic view) and 200 µm (zoom-in view). NESC: neuroepithelial stem cell; RG: radial glia cells; vRG_E_: early ventricular RG cells; vRG_L_: late ventricular RG cells; oRG: outer RG cells; tRG: truncated RG cells. A:anterior; FR: frontal; PF: Prefrontal; MS: motor-somatosensory; Mot+Par: motor + parietal; Tem: temporal; P: posterior; OC: occipital; Vis: visual; VZ: ventricular zone; SVZ/OSVZ: subventricular/outer subventricular zone; IZ: intermediate zone; CP: cortical plate.

Within the region-shared subtypes, we defined the molecular programs employed by all the regions along the progression of NSCs in the VZ and the oRG cells in outer SVZ (OSVZ) (fig. S7, E and F). Interestingly, chromatin remodeling factors, including *HMGA2, JARID2* and *SMARCC1*, and TFs such as *TSHZ2* and *SP8* dominated in the transitions of the NESCs into vRG_E_ cells. The transition from tRG to ependymal cells was characterized by the expression of cilia genes such as *FOXJ1*. Cell adhesion and cell-cell signaling genes, including synaptic components such as Neurexins (*NRXN1* and *NRXN2*) and angiogenesis molecules such as the vascular endothelial growth factor (*VEGFA*), were enriched across the oRG cell development (fig. S7, E and F). These data determined the key NSC states emerging across corticogenesis and the distinct gene cascades underlying their lineage transitions.

Differential gene expression analysis performed between the regions and along the developmental lineage of the VZ and OSVZ NSCs showed accentuated area diversification at the beginning of the NSC progression, i.e., in NESCs and vRG_E_ cells, and in the late oRG cells (*HOPX*^+^/*APOE*^+^) (Fig. 4C). This analysis defined regionally-enriched gene expression cascades both for the VZ (apical) and the SVZ (basal) NSC lineages. Within the VZ NSC progression, prominent region-specific gene expression was centered at early phases in the frontal region (Fig. 4D). Temporally-regulated genes included multiple TFs (*ZNF219, ZIC1-5* and *SOX21*), WNT (*WNT7B, WNT8B*) and RA signaling members (*CYP26A1* and *RBP1*) (Fig. 4D and fig. S8A; table S3). Surprisingly, several classic oRG cell marker genes, including *HOPX, PTN, FABP7* and *PMP22*, were enriched selectively in the early frontal VZ NSCs, representing unique features of this population. However, this enrichment in the frontal region was reduced in more mature VZ NSCs (fig. S8, B and C). Genes enriched in the motor-somatosensory NSCs included *SOX3* and *EMX2*. Both temporal and occipital regions displayed higher expression of *NR2F1, NR2F2, FGFR3* and WNT (*RSPO3*) and Notch (*HES5*) signaling members during the early phases (Fig. 4D; figs. S3Bii, S4D and S8A).

We also identified genes displaying comparable expression dynamics in multiple regions (fig. S8, D and E). The identified 199 genes were dominated by expression gradient showing enrichment in the early frontal and occipital ventricular RG cells, compared to the motor-somatosensory region, and included *TET1, HES5, JUND* and *JUN* (fig. S8E). In contrast, after correlating gene expression dynamics with NSC progression pseudotime in frontal versus occipital region, 16 genes were found in both regions but exhibiting opposite temporal expression dynamics (fig. S8, D and F). For example, high expression of the neuropeptide *PENK* was observed first in early frontal NSCs, switching later to occipital ventricular NSCs (fig. S8G).

Along the oRG lineage, regional divergences were prominent at the final phase of their maturation (fig. S8H). Genes differentially expressed across regions included the RA signaling member *RBP1* in the frontal; *NR2F2* and the BDNF receptor gene *NTRK2* in the temporal; *MEF2C* and the neuropeptide *NPY* in the occipital region (fig. S8, H and I, table S4). Together, these results indicate temporally regulated region-specific gene expression patterns along VZ and OSVZ NSC progression.

Finally, we checked whether the molecular program of the vRG cells is relayed to the oRGs. Low or negative correlation was found between regionally-enriched genes in early ventricular RG cells (NESC and vRG_E_) versus oRGs, except for a few genes, including *RBP1, ZIC1* and *DCT* constantly expressed in both apical and basal RG cells of the frontal cortical wall (Fig. 4E). This result indicates that apical and basal RG cells employ divergent region-specific genetic programs across their maturation, however, in the frontal cortical wall, they might share certain molecular mechanisms including RA signaling genes.

The expression pattern of several region-specific TFs including ZICs, *SP8* and *LHX9* was validated by RNAscope analysis in fetal monkey tissue (Fig. 4F). *ZIC4* and SP8 in the anterior, and *LHX9, FEZF1* and NR2F1 in the posterior region, all highly expressed at E40, decreased at later phases in these domains. In contrast, expression of *ZIC3, ZIC1, MEIS2* and *PBX1*, mainly detected in the antero-ventral domain at E40, spread to the ventricular RG cells across the antero-caudal axis at E52 (compare Figs. 2C and 4F, fig. S3B). Together, these data indicate that the TF gradients generated at early phases in the telencephalic organizer domains are transient, and their spatial expression changes across development (*4, 43*). Thus, the developing monkey telencephalic regions involve a code of sequentially regulated genes instructing the initial identity of the NSCs and mediating their progression through defined region-specific state transitions.

### Transcriptomic diversification of the excitatory neurons across prospective cortical areas

Cortical areas are populated by a heterogeneous composition of excitatory neurons organized in connections, reflecting the specialized function of each domain (*44, 45*). Functional connectivity contributes to refining arealization during corticogenesis (*46*). We investigated the molecular events underlying the emergence of excitatory neuron diversification across the cortical regions, from proliferating IPCs transitioning into post-mitotic neurons.

Unsupervised clustering and marker gene expression identified distinct subtypes across the differentiation lineages, from *EOMES*^+^ IPCs to deep (*SOX5*^+^) and upper (*CUX2*+) layer excitatory neurons, with their trajectories further depicted by pseudotime analysis (Fig. 5A, fig. S9, A and B). Multiple waves of deep and upper layer neurogenesis were detected throughout the timeline. Deep layer neurons emerged at E37-43, although at a low rate, consistent with them being generated early (*47*) (fig. S9C). Then, they peaked at E54-64, except for the *OPRK1*^+^ subtypes (putative L6 intratelencephalic neurons) enriched at E93. The upper layer lineage was more evident at E77-78 and then enriched at E93 (fig. S9A). Such dynamics also exhibit variations across cortical regions. For instance, upper layer neurons were generated in the occipital region later than the other regions; however, in this region, they showed faster maturation at E77-78, likely due to greater myelination at this time (*11, 24*) (fig. S10, A and B).

**Fig. 5.**
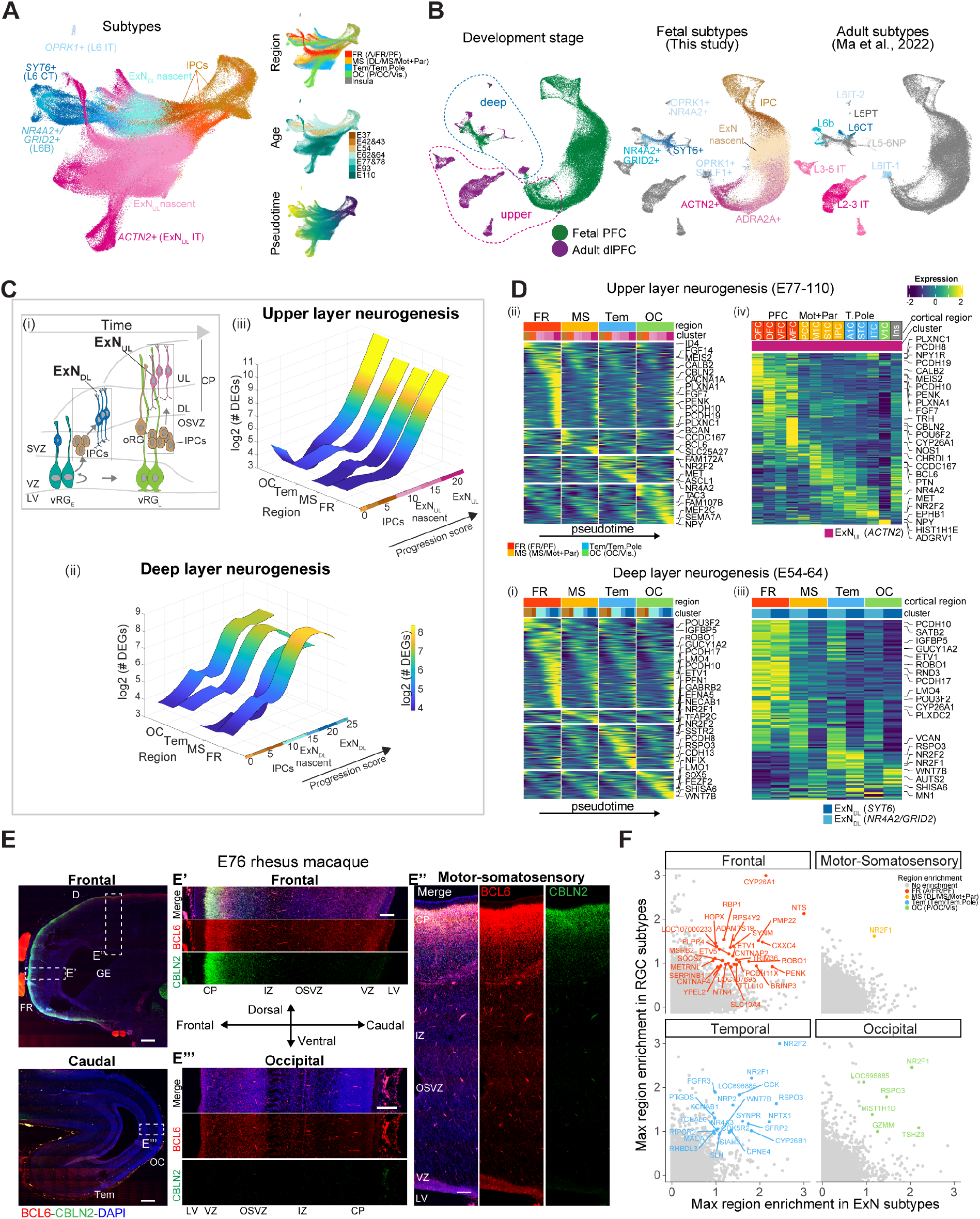
Spatial transcriptomic divergences across cortical neuronal trajectory. (**A**) UMAP of IPCs generating deep and upper layer cortical neurons showing cell subtypes, region, age and pseudotime. The legend indicates the regions considered across the time, as indicated in Fig. 1. (**B**) Transcriptomic integration of macaque fetal (this study) and adult dorsolateral prefrontal cortex (dlPFC) excitatory neurons (*48*). FR/PFC: frontal/PF regions. (**C**) (i) Schema of neurogenesis. Early and late radial glial cells (vRG_E/L_) generate IPCs and deep (DL) or upper layer (UL) excitatory neurons (ExN_DL_ or ExN_UL_), respectively. (ii and iii) Number (in log2 scale) of differential expressed genes (DEGs) between regions along the differentiation of DL (ii) and UL (iii) neurons. (**D**) Region-specific gene expression cascades along the pseudotime from IPCs to DL (i) or UL (ii) neurons. Expression level of genes in DL (iii) or UL (iv) layer differentiated excitatory neurons across more refined cortical areas. Region legend as in A (**E**) Expression of *BCL6* and *CBLN2* detected by RNAscope in E76 macaque brain sagittal sections. Scale bars: 500 µm (panoramic view) and 200 µm (zoom-in view, E’, E’’, E’’’). **(F)** Shared region-specific genes between RG cells and excitatory neurons.

To better delineate the identities of these excitatory neuron subtypes, we performed transcriptomic integration with a single nucleus RNA-seq dataset of adult macaque dorsolateral prefrontal cortex (*48*) (Fig. 5B and fig. S9D). The analysis showed co-clustering of fetal and adult deep layer neurons on the UMAP layout, predicting the mature identities of multiple fetal deep layer neurons, including an L6B (or subplate *NR4A2*^+^/*GRID2*^+^), an L6 corticothalamic (*SYT6*^+^) and two L6 intratelencephalic (*OPRK1*^+^) subtypes. In contrast, fetal upper layer neurons marked by *CUX2* expression did not diversify into different clusters of cells as shown by the adult upper layer intratelencephalic neurons. These data suggest that deep layer excitatory neurons might acquire identities comparable to their adult states relatively soon after they are generated. In contrast, upper layer neurons probably require additional maturation, as their final diversification was not evident in the fetal time analyzed. These analyses precisely define the molecular features of the developing neurons across the laminar organization of the primate neocortex.

The region and subtype information overlaying on the same UMAP layout suggested that neurons across regions become more transcriptomically distinguishable along their maturation pathway (Fig. 5A). This pattern was further validated via independent differential expression analysis and AUC scores confirming that prominent regional divergence occured during the late phases of both deep and upper layer neuronal differentiation (Fig. 5C and fig. S10C). Moreover, hierarchical clustering of the excitatory neurons from the distinct regions at E93 showed that they clustered into three groups at their terminal differentiation, belonging to the prefrontal, the motor-somatosensory and temporal, and the visual region (fig. S10D). Regional differences arised in IPCs and expanded as neuron differentiated, however, those detected before the maturation phase were not sufficient to reflect the anatomical proximity of the regions.

To understand the molecular programs underlying regional diversifications of excitatory neurons, we calculated genes enriched in each brain area for the deep and upper layer neuron lineages (fig. S10E). While the region-specific signatures largely overlapped between the two mature deep layer excitatory neuron subtypes (*SOX5*^+^/*NR4A2*^+^/*GRID2*^+^ and *SOX5*^+^/*SYT6*+), they were scarcely shared between mature deep and upper layer neurons. This indicates distinct molecular programs governing the regional identities of deep versus upper layer neurons. Furthermore, by ordering the region-specific genes from all the related subtypes along the pseudotime from IPCs to neurons, based on their expression peaks, we confirmed regional gene expression differences enriched in the late neuronal maturation phase. However, we detected non-negligible regional differences in IPCs, representing earlier cell-autonomous events contributing to neuron diversifications (Fig. 5, Di and Dii).

At the terminally differentiated phases, genes distinguishing frontal deep layer neurons included *PCDH10* and *PCDH17*, involved in cell adhesion, and *ROBO1* which modulates neurogenesis (*49*), consistent with previous data (*29*). The TFs *NR2F1, NR2F2* and *RSPO3* were among the genes distinguishing the temporal cortex. Likewise, *SOX5, WNT7B* and *AUTS2*, this implicated in neuropsychiatric disorders, defined occipital deep layer neuron identities (Fig. 5Diii). Upper layer excitatory neurons were abundantly revealed at E93 and E110 when more refined cortical regions were sampled (Fig. 5Div). Genes denoting regional divergence at late differentiation included the RA signaling members *CYP26A1* and *CBLN2* (*34, 50*), enriched in MFC and the RA target *MEIS2*, enriched in OFC. The neuropeptide *PENK* was enriched in DFC while the other neuropeptides *PTN* and *NPY* were specifically expressed in the prospective M1C and V1C, respectively. The divergent expression pattern across regions was further validated for some genes by RNAscope on E76 macaque brains, as exemplified by the antero-caudal gradient of *BCL6* and *CBLN2* (Fig. 5E). These analyses delineate the developmental dynamics of the diversification of the cortical excitatory neurons, highlighting a refinement of the regional identity in their terminally differentiating phase.

Finally, we asked whether the early protomap of the RG cells was related to the area-specificity of the post-mitotic excitatory neurons. Analysis of gene expression enrichment of the RG and neuron subtypes identified those genes constantly defining regional identity across the differentiation trajectory. These included *HOPX* and *ROBO1* and RA signaling-related genes, such as *CYP26A1* and *RBP1* in the frontal region, consistent with the role of RA signaling in patterning the prefrontal cortex of primates (*34*); *NR2F2* and *CYP26B1* in temporal; *NR2F1* and *RSPO3* in occipital (Fig. 5F, and fig. S10F). Thus, we found the genes expressed from RGs till mature neurons, likely representing the fundamental elements instructing and maintaining the regional identity of the neural cells across development. Altogether these data denote the contribution of early cell-autonomous mechanisms in shaping later neuronal diversity across neocortical areas.

### Limited diversification of the GABAergic inhibitory neurons across developing cortical areas

The neocortex is also populated by the GABAergic inhibitory neurons generated by the GE, migrating afterwards to the developing dorsal areas (*51*). Employing clustering, marker gene expression and transcriptomic integration with an independent developing macaque inhibitory neuron scRNA-seq dataset (*52*), we identified different subtypes. These were represented by MGE- (*LHX6*^+^) and CGE-(*NR2F1*^*+*^ and/or *SP8*^+^) derived cortical inhibitory neurons, striatum spiny projection neurons (*FOXP2*^+^ and/or *FOXP1*^*+*^), primate-specific MGE-derived (*LHX6*^+^/*CRABP1*^+^) inhibitory neurons (*16, 52*), and olfactory bulb neurons (*PAX6*^+^/*SP8*^+^), selectively abundant in the prefrontal cortex (fig. S11, A and B). Transcriptomic integration with adult macaque brain scRNAseq data (*48*) showed that mature inhibitory neurons reside at the tip of major inhibitory neuron lineages, denoting potential developing trajectories from fetal to adult (fig. S11C). These data confirmed well-known lineages giving rise to adult VIP, *LAMP5*^+^/*LHX6*^+^ and long range projecting (*SST*^*+*^*/NPY*^*+*^) interneurons (*16, 48, 53, 54*). In addition, the data highlighted fetal *NR2F2*^*+*^*/LAMP5*^*+*^ subtype enroute to adult *LAMP5*^*+*^*/RELN*^*+*^ interneurons, largely representing neurogliaform cells; fetal *NR2F2*^*+*^*/SP8*^*+*^*/KIT*^*+*^ subtype contributing to adult *ADARB2*^*+*^*/KCNG1*^*+*^ interneurons; fetal *SST*^*+*^*/GUCY1A2*^*+*^ contributing to a subset of *SST*^+^ and *PVALB*^+^ interneurons, likely also including the adult cortical *TH*-expressing interneurons (*48*) (fig. S11C).

Inhibitory neurons are specified in the subpallium, long before they reach the neocortex (*16, 55*). However, if and how they are further diversified across the cortical regions is still debated (*53*). Co-clustering of inhibitory neurons from different regions showed little separation on the UMAP layout (fig. S11A). Moreover, pairwise transcriptomic comparison of the *LHX6*^+^ and *NR2F1*^*+*^/*SP8*^+^ subtypes between cortical regions found high similarity for most of the neuron pairs (AUC 0.682 ± 0.004) (fig. S11D). However, these inhibitory neuron subtypes exhibited more transcriptomic uniqueness in DFC and MFC, IPC and insula than in other regions (fig. S11D). Notably, hierarchical clustering of regionally-divergent genes recapitulated the transcriptomic uniqueness and identified distinct expression patterns across these same cortical regions (fig. S11E). Such divergent expression patterns were not specific to these *LHX6*^+^ inhibitory neurons as they were also recapitulated by the *NR2F1*^*+*^/*SP8*^+^ ones, and vice versa. Moreover, they were further confirmed by the high correlation coefficients (0.78±0.03) of the regional gene expression enrichment between *LHX6*^+^ and *NR2F1*^*+*^/*SP8*^+^ interneurons (fig. S11, E and F). These results indicate transcriptional overlapping of regional identities between interneurons, suggesting that later cues might contribute to their further regional diversification in the cortex (*56*).

### Spatial transcriptomic divergences across gliogenesis

We next focused on the developmental transition of RG cells across gliogenesis. Unsupervised transcriptomic clustering analysis defined discrete phases of differentiation across the gliogenic trajectory. The analysis distinguished late RG cells transitioning into glial intermediate precursors (gIPCs) and then diverging towards astrocytes or oligodendrocytes, whose identity was confirmed by canonical markers (Fig. 6A and fig. S12A). Comparative analyses with fetal human and mouse datasets (*22, 57, 58*) denoted matching between glial subtypes and confirmed the annotation (fig. S12B). Leveraging the abundant astrocytes profiled, we identified three astrocyte subtypes, one (Astro *EGFR*) enriched in prefrontal regions and the others (Astro *GFAP* and Astro *MFGE8*) shared across regions (fig. S12C). Moreover, aligning astrocyte subtypes to adult stages (*48, 59*) showed that the *GFAP*^+^ subtype, expressing *AQP4* and the TFs *SOX2* and *ID3*, was transcriptomically more similar to interlaminar astrocytes, while *EGFR*^+^ and *MFGE8*^+^ subtypes expressing *SOX2* and the NOTCH targets *HES1 or HES5*, respectively, resembled protoplasmic astrocytes (fig. S12C). These results suggest that adult astrocyte identities emerge at a mid-fetal stage in macaques.

**Fig. 6.**
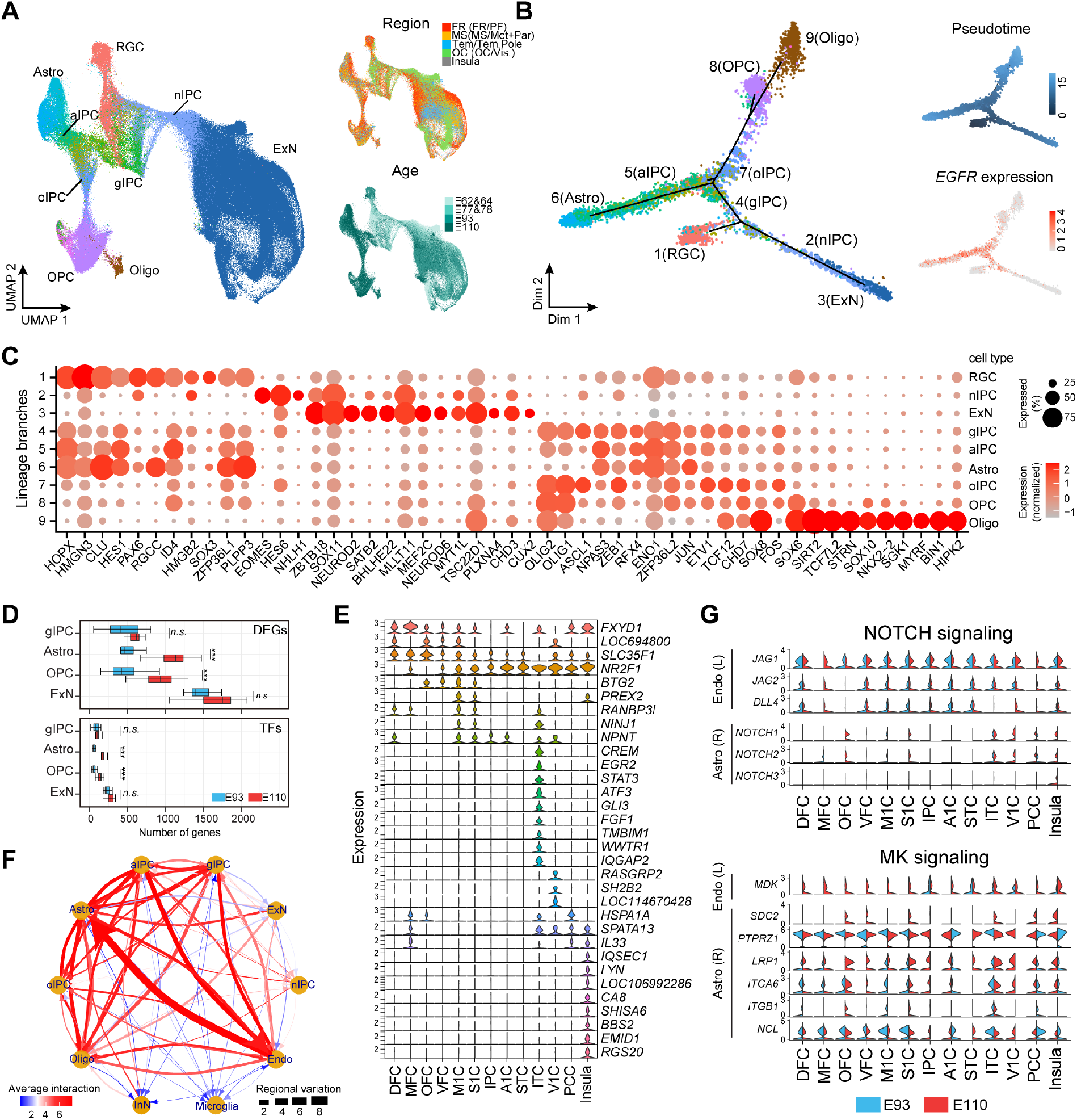
Transcriptomic versatility of gliogenesis across the cortical areas. **(A)** UMAP showing the neurogenic and gliogenic trajectories of the late RG cells (RGCs), regions and age. The regions considered across the time are as in Fig. 1. **(B)** Cell trajectory built by Monocle 2 defining the transcriptomic transition of the RGCs toward neurogenesis or gliogenesis. Main branches are numbered. The insets illustrate the pseudotime and *EGFR* expression. **(C)** Expression of the top 10 transcription factor genes specific to each branch. **(D)** Boxplots showing the numbers of genes (upper) and TFs (bottom) with regional differential expression in gIPCs, Astro, OPC and ExN at E93 (blue) and E110 (red). One-side Student t-test between E93 and E110 conducted for each cell type, P<0.001 ***, P<0.01 **, P<0.05*. **(E)** Expression of curated genes with region-specific expression in astrocytes. **(F)** Overview of the predicted L-R mediated cell-cell interactions. The arrows show the direction from the L-to the R-expressing cells. **(G)** Expression of selected L and R genes expressed by endothelial cells (Endo) or Astro, respectively, involved in the signaling pathways across all the regions at E93 and 110.

We next sought to define the transcriptomic programs governing the switch of the RG cells from neurogenic to gliogenic potential, identifying the top-ranked genes expressed at each lineage branch point. Cell trajectory reconstruction and pseudotime analysis determined the developmental transition of RG cells into excitatory neurons or glial lineage, this marked by high *EGFR* expression (*57, 60*) (Fig. 6B). Analysis of the genes expressed at the bifurcation of each trajectory highlighted well-known regulator of astrocytic fate, such as the chromatin remodeling factor *HMGN3*, and the TFs *PAX6, HES1, SOX2, SOX3* and *SOX6* in the late RGs. *SOX11*, and NEUROD genes were expressed in the neurogenic lineage. Gliogenic IPCs expressed *OLIG1, OLIG2* and *ASCL1*, sharing transcriptional features with both oligodendrocyte (oIPCs) and astrocyte (aIPC) progenitors. Distinct molecular signatures were found in the oligodendrocytes, expressing *SOX10* and *NKX2-2*, unlike astrocytes which shared more signature features with the NSCs (Fig. 6C and fig. S12D, table S5). These data reveal the temporally orchestrated combination of genes at the fate switch from neurogenic to gliogenic RG cells during monkey corticogenesis (*61, 62*).

Astrocytes are characterized by a remarkable diversity, however, how this heterogeneity emerges remains elusive (*63*). The spatial transcriptomic divergence of the oRGs, which generate neurons and glia (*64, 65*), showed above, suggested events of diversification of the glia encoded by the progenitors. Hence, we asked whether glial cells trascriptomically diversify across the areas of the developing neocortex. Differential gene expression analysis indicated that glial cell identities diverge across the regions less than excitatory neurons. However, astrocytes displayed higher differential expressed genes (DEGs), including TFs, than gliogenic precursors and oligodendrocytes at E110 (Fig. 6D and fig. S12E). Moreover, astrocytes were distinguished by the expression of region-specific genes including FXYD Domain Containing Ion Transport Regulator 1 (*FXYD1*) in the DFC, MFC and insula, *STAT3* in the temporal (ITC) and the AMPA Receptor component *SHISA6* in insula astrocytes (Fig. 6E). The data denote specific molecular features of the astrocytes of the different cortical regions. Together, these results highlight that transcriptomic variation of the astrocytes across the cortical regions is more accentuated at the late phase of their differentiation trajectory.

### Region-specific competence of the astrocytes to environmental cues

We investigated whether the interaction of neural cells with non-neural cells might be associated with region specific transcriptomic features of the neocortex. We predicted, as described above, the ligand-receptor pairs mediating putative signaling interactions among neuronal, glia and non-neural cells. The highest number of interactions that also showed the most prominent regional variation was predicted to be between endothelial cells, which line the blood vessels, and the astrocytes (Fig. 6F). Astrocyte development is influenced by signaling from endothelial cells (*66*). Hence, we wondered whether the interaction between endothelial cells and astrocytes, expressing ligands and corresponding receptors, respectively, might relate to the regional diversity of the astrocytes. Of note, we found a low transcriptional variation of the endothelial cells among the regions at E110, when they are more abundant (Fig. 1 and fig. S13A), implying that astrocytes, from distinct areas and time points, respond differently to endothelial signals. We identified ligand-receptor pairs of diverse signaling pathways displaying high regulation across time and regions (fig. S13B). However, pairs of several signaling pathways including Midkine (MK) and Notch displayed similar expression of the ligand genes (*MDK*, and *JAG1, JAG2, DLL4*, respectively) in the endothelial cells, in contrast to the variation of the paired receptors in the astrocytes (*SDC2, ITGB1* and *NOTCH1, NOTCH2, NOTCH3*, respectively) (Fig. 6G and fig. S13C). The data indicate that astrocytes have intrinsic region-specific competence to respond to the endothelial cells’ signals which might lead to their transcriptomic variation across regions. Together, these data indicate that the regional diversity of the astrocytes is in part cell-autonomous, that is, not shaped only by environmental cues.

### Spatiotemporal expression of disease-risk genes in early telencephalic development

Alterations of cortical development have been implicated in neurological and psychiatric diseases (*67*). While disease genes show often enrichment in neuronal cells (*24, 68-70*), little is known about their function in NSCs and across the early fetal development of the telencephalon. We curated gene lists associated with major neuropsychiatric diseases, including autism spectrum disorder (ASD), schizophrenia (SCZ) and attention-deficit/hyperactivity disorder (ADHD), neurodegenerative disorders such as Alzheimer’s disease (AD), as well as brain cancers including glioblastoma. Then, we mapped their expression dynamics into the spatiotemporal progression of the developing macaque telencephalon (table S6). Expression enrichment analysis was assessed for each disease gene list across all the cell subtypes (Fig. 7A and fig. S14A). Confirming previous studies, the analysis showed that risk genes for multiple neuropsychiatric disorders, including ASD, SCZ and developmental delay (DD), were largely enriched in excitatory and inhibitory neurons (*68, 71, 72*). Glioblastoma genes showed dominant expression in glial precursors and oRG, confirming common transcriptomic features between cancer cells and fetal brain progenitors (*73, 74*).

**Fig. 7.**
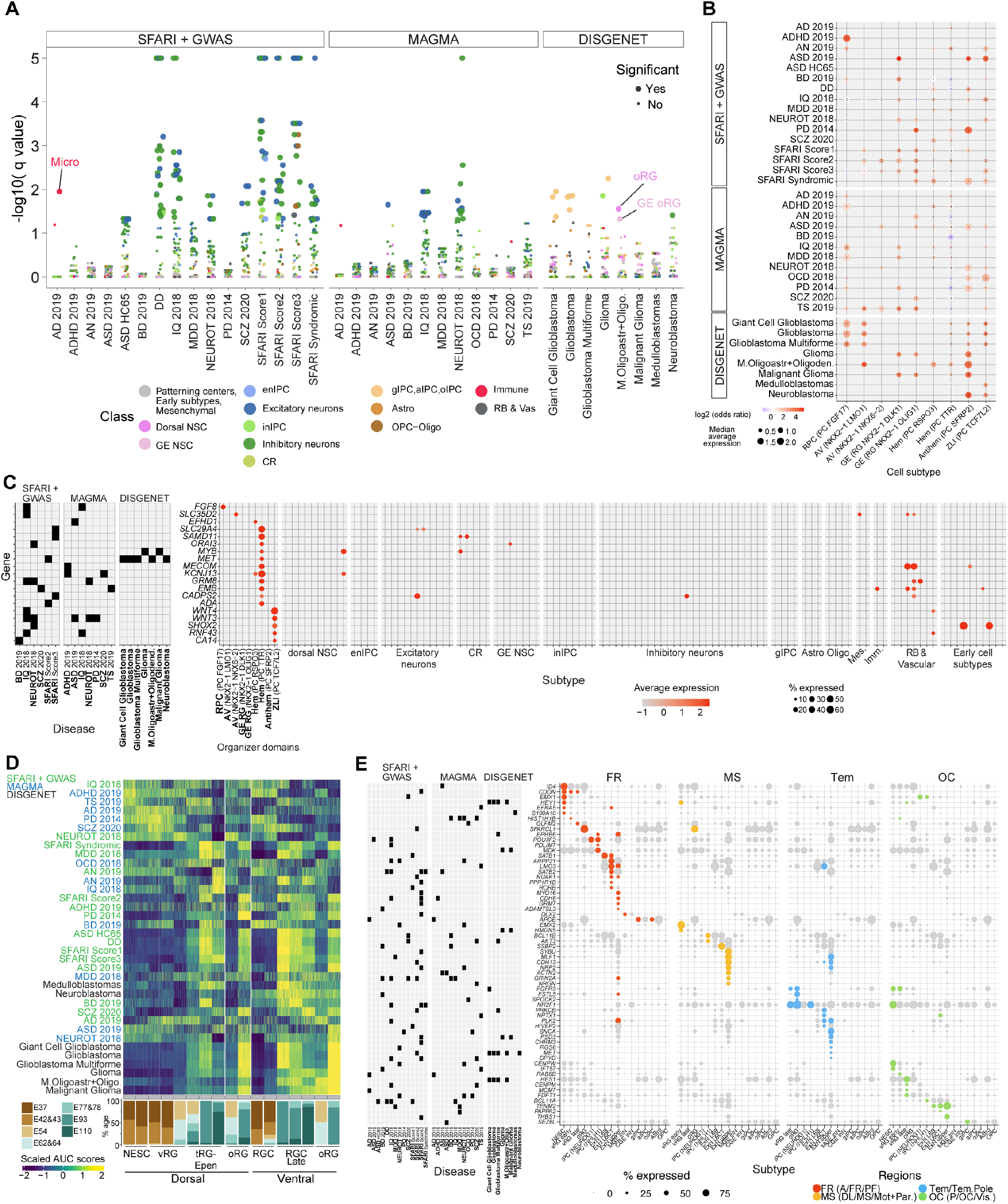
Expression of brain disease-risk genes in early telencephalic development. (**A**) Expression enrichment of disease gene sets across cell subtypes (dots), with significant ones (q value < 0.05) plotted as big dots. (**B**) Dot-plot summarizing the disease gene expression across organizer domain subtypes. The number inside each dot represents subtype markers overlapping with disease risk genes. The size of each dot represents the median average expression of the overlapping markers. Dot color indicates the odds ratio of a disease risk gene being a subtype marker. (**C**) Scaled gene expression for disease genes selectively enriched in a subset of organizer domains versus other neurogenic and gliogenic lineage subtypes. (**D**) Scaled AUC scores assessing the expression enrichment of disease genesets across NSCs. Genesets are ordered based on the peak enrichment along the horizontal axis (dorsal and ventral NSCs). (**E**) Expression of the disease genes overlapped with the top regionally-enriched cell subtype markers. For a given gene, lightgrey colors refer to no regional enrichment while region colors denote enrichment.

Although the analysis across all cell subtypes could unveil the most salient signals, it might mask the gene expression patterns in others cell types, such as brain organizers and NSCs. These cells might have implications in the origin of the brain diseases, as regional specification and fate transitions of NSCs were suggested as early fetal risk events for neurodevelopmental disorders (*75*). While we did not observe any disease gene set enriched in organizer domain subtypes (Fig. 7A and fig. S14A), an average ranging from 26.1% to 36.2% of the genes from each list was expressed in these cells (expressing ≥5% cells) (fig. S14B). Among these risk genes, 132 overlapped with patterning center subtype markers, of which 19 were unique to organizer domain subtypes and other 113 also appeared later in development in other cells (Fig. 7, B and C and fig. S14C). The 19 genes included well-known genes such as *FGF8* expressed in the rostral patterning center, and novel risk genes with restricted expression in a particular organizer domain. For example, the ASD- and glioma-associated gene *MET*, previously detected in the postnatal excitatory neurons of humans, but not in other non-human primates (*11, 23, 48*), was found to be expressed early during telencephalic development selectively in the cortical hem (Fig. 7C). The 113 genes included Dual Specificity Phosphatase 6 (*DUSP6*), a member of the mitogen-activated protein kinases (MAPKs), associated with attention-deficit/hyperactivity disorder (ADHD) and glioblastomas which was expressed early in the antero-ventral telencephalic organizers and later in glial cells (fig. S14C). These analyses indicate that certain disease risk genes commonly associated with mid- or late-gestation neural dysfunctions are expressed earlier in the telencephalic organizers, with a putative role during the phase of regional patterning of the telencephalon and NSC specification.

Similar enrichment of the disease risk genes was conducted along the progression of dorsal and ventral NSCs (Fig. 7D). Several gene sets, including the multiple ASD-associated, displayed a predominant enrichment in tRGs and/or oRGs. Similar pattern was observed for glioblastoma-associated genes, consistent with previous findings (*76*). However, ADHD, Tourette syndrome (TS), SCZ genes and two ASD gene lists exhibited early expression emerging in E37-43 dorsal NSCs. Interestingly, ADHD and bipolar disorder (BD) risk genes were highly represented in the early ventral NSCs. These spatiotemporal enrichment patterns were not affected by sequencing depth bias, as demonstrated by the consistent dynamics shown in the downsampled data (fig. S15A). Correlating gene expression with the gene set enrichment patterns further revealed the key genes driving the spatiotemporal expression preference of the disease genes (fig. S15B). These results indicate that neurodevelopmental disorder risk genes might function across the progression of dorsal and ventral telencephalic NSCs from early to late states, likely affecting a myriad of derived neural cell types.

Different brain areas might show variable vulnerability to a disease, raising the need to define the spatial and temporal patterns of risk-conferring gene expression across telencephalic regions. Thus, we intersected disease risk genes with top regionally-enriched cell subtype markers (Fig. 7E and fig. S16A). The results show that the ASD gene *CDON* was preferentially expressed by frontal neuroepithelial stem cells, *SATB1, SATB2* and *ARPP21* were enriched in frontal deep layer excitatory neurons and *MYO16, CDH8* and *GRM7* were enriched in frontal upper layer excitatory neurons. Similarly, the glioblastoma genes *HEY1 and HES1* were preferentially expressed by frontal and occipital NSCs, respectively, reflecting the stem cell-like features of glioblastoma (Fig. 7E and fig. S16). Thus, this analysis determined spatial- and cell type-expression bias of the disease-associated genes across primate telencephalic development.

In conclusion, these data point to telencephalic regional patterning and NSC progression as risk events for the origins of neurodevelopmental disorders. Moreover, the molecular programs underlying these neurodevelopmental events might be dysfunctionally recapitulated in brain cancers (*77*).

## DISCUSSION

This work reveals the transcriptomic programs and cellular events underlying the establishment of the telencephalic regional identity across macaque fetal brain development.

The protomap model postulates that NSCs determine the layout of the cerebral cortex and the partition of the adult areas at the earliest stages of fetal development, by a coordinated expression of genes and enhancers’ activity operating in discrete domains (*5, 37*). However, studying these mechanisms in such early fetal post-mortem human samples is challenging. Thus, this work with fetal monkey brain may contribute to better define the early events underlying the specification of the NSCs and the diversification of the telencephalic regions in primates.

We show that the molecular mechanisms orchestrating regional identity are dynamic across telencephalic development. We found that the brain organizer states are transient (*4, 22*), although it is well-established that regional patterning continues across the developing cortex, later also regulating neuronal lamination and activity (*34, 35*). We observed that the early regionalized gradients of morphogens signal to competent NSCs, likely propagating region-specific gene cascades during the progression of the telencephalic RG cells. Thus, an initial molecular program might instruct the subsequent state transitions of the NSCs and their response to the developmental cues. Moreover, we found that certain key players are expressed from the early cortical NSCs throughout the differentiating neurons, suggesting a putative function for these genes to maintain the regional identity across the time. We found extensive regional variation in the early NSCs, however, these divergences are reduced at later phases of the RG progression, becoming again high in neurons and astrocytes. Importantly, the high regional variation that we reveal in the oRG cells underscores a further mechanism of area diversification in the OSVZ. Thus, our data highlight two waves of spatial diversification of the NSCs in the germinal zones, one for the early apical and one for the late basal RG cells.

Neurons and astrocytes show high regional divergence in their terminal differentiation phase, likely due to subcortical afferents, and cell interactions including synaptic inputs and myelination (*11, 46*). Region-specific gene cascades re-emerge from the IPC state throughout the neurogenic trajectory, suggesting that cell-autonomous programs in the proliferating precursors also might contribute to the diversification of the post-mitotic neurons. Interestingly, astrocytes show different competence to respond to endothelial cells’ signals, suggesting that their regional identity might also in part be intrinsic.

The origin of the neuron and astrocyte heterogeneity across cortical regions remains unresolved in this work. However, these data support a model where cell-autonomous programs characterize the early specification of the NSCs and their spontaneous progression, and then an interplay with extrinsic cues might further shape the identity of neurons and astrocytes during late corticogenesis. These cell intrinsic events, driving arealization and the further persistence of regional-specific features across NSC maturation and differentiation, also emerge here with the organoids. We show that brain organoids maintain the induced regional identity and differentiation program weeks after that the exposure to patterning ligands is terminated, and in more prolonged cultures, even in the absence of non-neural cells and extrinsic neuronal inputs (*39*).

This work also highlights that human and mouse telencephalons are governed by similar mechanisms of regional patterning (*4*). However, we found that the neuropeptides *GALP* and *GAL* are expressed in the antero-ventral domain of the early developing monkey telencephalon, but were not detected in the mouse at equivalent fetal stages. The data here highlight an interesting novel role for these neuropetides in early fetal regionalized NSCs, as they have been reported to be associated with more mature cortico-hypothalamic circuitries (*78*). The gain of *GALP* and *GAL* expression in developing primates could have enhanced the proliferation of the NSCs and contributed to the expansion of the frontal telencephalon across evolution, as suggested by our experiments with the brain organoids. Thus, we have identified novel mechanistic insights into the regional specification of the neural stem cells suggesting new candidate mechanisms for the evolutionary expansion of the telencephalon in primates. Many works point to neuronal dysfunctions occurring during human midfetal or late corticogenesis as the origins of neuropsychiatric disorders (*24, 68-70*). We show that many risk genes associated with mental diseases, including SCZ, ASD, ADHD and BD, map early in development, as they result to be expressed in brain organizers and in dorsal and ventral NSCs. Thus, we speculate that these disorders might also have an earlier neurodevelopmental origin, implicating dysfunctional patterning of the telencephalon and altered spatiotemporal identity of the RG cells. In general, these data indicate multiple phases of vulnerability to gene lesions across telencephalic development and corticogenesis, involving regional patterning of the NSCs and their transition towards neurogenic and gliogenic trajectories. These programs underlying fetal NSC states might also lead to brain cancers when dysfunctionally recapitulated in adults (*73, 77*). Interestingly, we show that glioblastoma and neurodevelopmental disorders, such as ASD and ADHD, share genes implicated in brain organizer and NSC functions, suggesting that gene lesions causing both disease types might converge to alter, in two different contests (fetal versus adult), similar NSC programs (*79-81*). This expression analysis of the brain cancer genes across monkey telencephalic development will also contribute to more comprehensively map the heterogeneity of the tumor cells into a primate neurodevelopmental landscape including early NSC states, and apply more targeted interventions.

Finally, this resource may guide the exploration of relevant development-related and disease-associated genes across the progressing cell types characterizing telencephalic development and corticogenesis. Integrating these data with other experimental platforms and brain multi-omics datasets will help better understanding the mechanisms governing human brain formation, evolution and diseases and even conceive more efficient experimental systems for modeling neurogenesis and its disorders *in vitro*.

### Summary of Materials and Methods

A full description of the materials and methods is available in the supplementary materials. Briefly, all procedures involving animals were carried out according to guidelines described in the Guide for the Care and Use of Laboratory Animals and were approved by the Yale University Institutional Animal Care and Use Committee (IACUC).

Single cell RNA-sequencing was performed using the 10X Genomics Single Cell 30 RNA-Seq V3 libraries and sequencing protocol. The alignment of scRNA-seq reads and the following barcode and UMI quantification were performed using CellRaner with the rhesus macaque genome assembly Mmul10 and NCBI RefSeq annotation (release 103). Following quality control removing low-quality cells and doublets, we performed normalization, batch correction, dimension reduction, clustering, and annotating cell types based on marker gene expression and transcriptomic comparison with published datasets. We conducted lineage inference to delineate the shared and divergent gene expression dynamics across brain regions, and predicted ligand-receptor mediated cell-cell interactions based on the expression of ligand genes in one cell type and the corresponding receptor genes on another cell type. Finally, we curated the gene lists associated with major brain disorders and assessed the expression patterns of these gene lists across cell types and intersected with the region-specific molecular signatures.

Telencephalic organoids from human and macaque iPSCs were generated by the directed differentiation protocol as previously described (*34*).

## Acknowledgements

The authors are grateful to Yale center for genome analysis (YCGA) for their help in building the scRNA-seq libraries and performing sequencing, Yale animal resource center (YARC) for macaque breeding, Yale veterinary clinical service for the surgery of the pregnant monkeys.

## Author contributions

N.M., S.M., M.L., S-K.K., N.S., P.R. conceived the paper and interpreted the results. N.M., J.A. and N.S. dissected the monkey fetal brains and collected the samples. N.M. and G.T. isolated cells for scRNA-seq. N.M. processed brain tissue for immuno-histrochemistry and RNAscope. S.M. and M.L. performed analysis of the single-cell dataset. S.M., X.M-B. and G.S. performed the analysis for Fig. 7 and related figures. S.M. coordinated bio-informatics and created the website. N.M. and S-K.K. generated human and monkey brain organoids and performed immunocytochemistry. N.M. and S.S. collected mouse embryos and performed immuno-histochemistry. N.M., S-K.K., S.S. performed imaging acquisition. N.M. and S-K.K performed image analysis. A.D. managed the rhesus monkey colony. N.M. designed the research and coordinate the work. N.S. and P.R. directed the research. N.M., S.M., M.L., G.S., N.S., P.R. wrote the manuscript. All authors edited the manuscript.

## Data and materials availability

The scRNA-seq data were deposited in the NEMO Archive (RRID: SCR_002001) under identifier dat-fjx1jbr accessible at https://assets.nemoarchive.org/dat-fjx1jbr. The data can also be interactively visualized at http://resources.sestanlab.org/devmacaquebrain.

