## Supplementary material for "Molecular programs of regional specification and neural stem cell fate progression in developing macaque telencephalon": Suppl.Figures & Methods

### SUPPLEMENTARY FIGURES

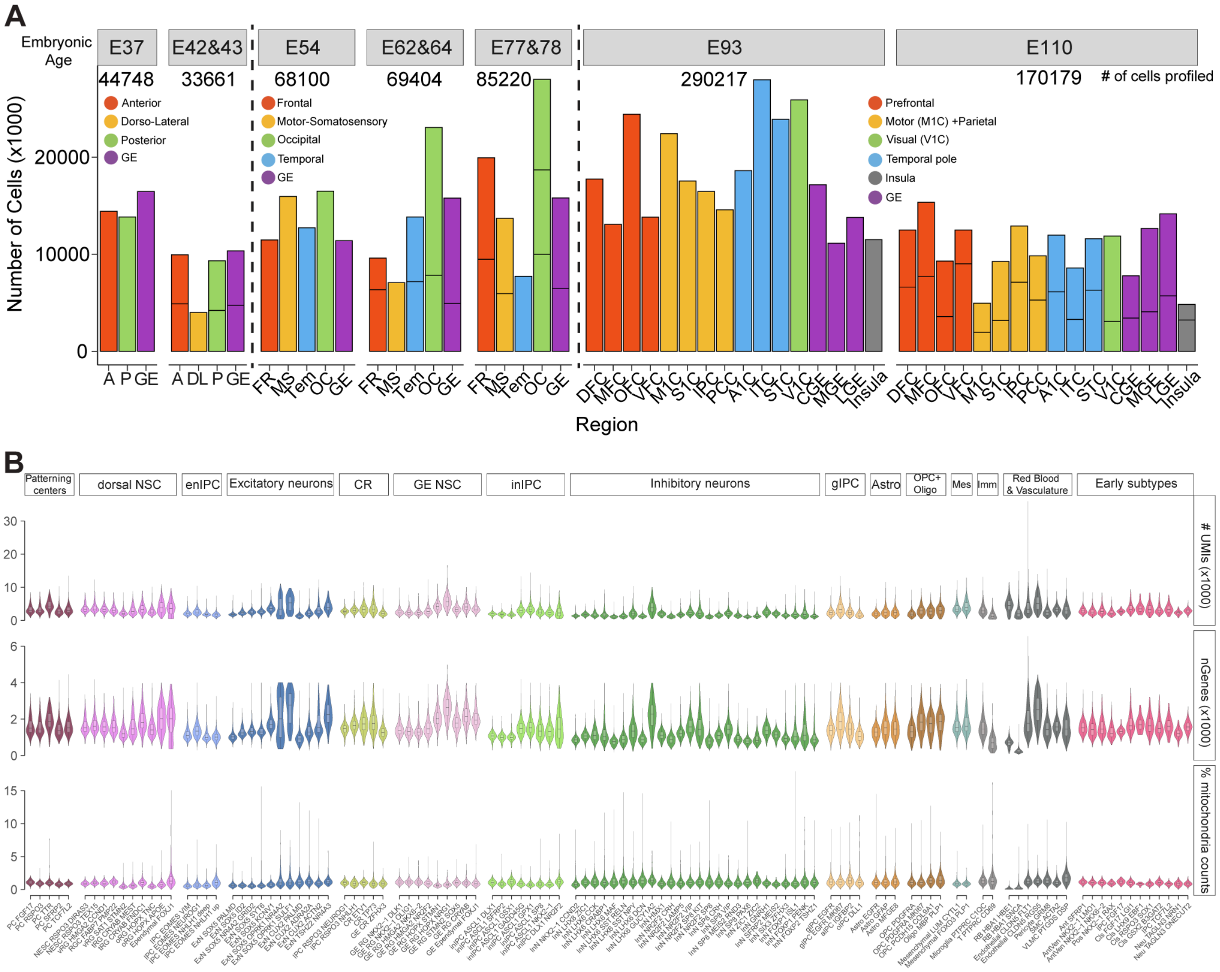


**Fig. S1. Quality overview of the macaque scRNA-seq data** (**A**) Bar plot displaying the sample size. Each segment represents a biological replicate. Number of cells at each age passing quality control is indicated. Cells are categorized in regions following the nomenclature of the regions across the time shown in Fig. 1. (**C**) Violin plot showing quality metrics of the identified cell subtypes.

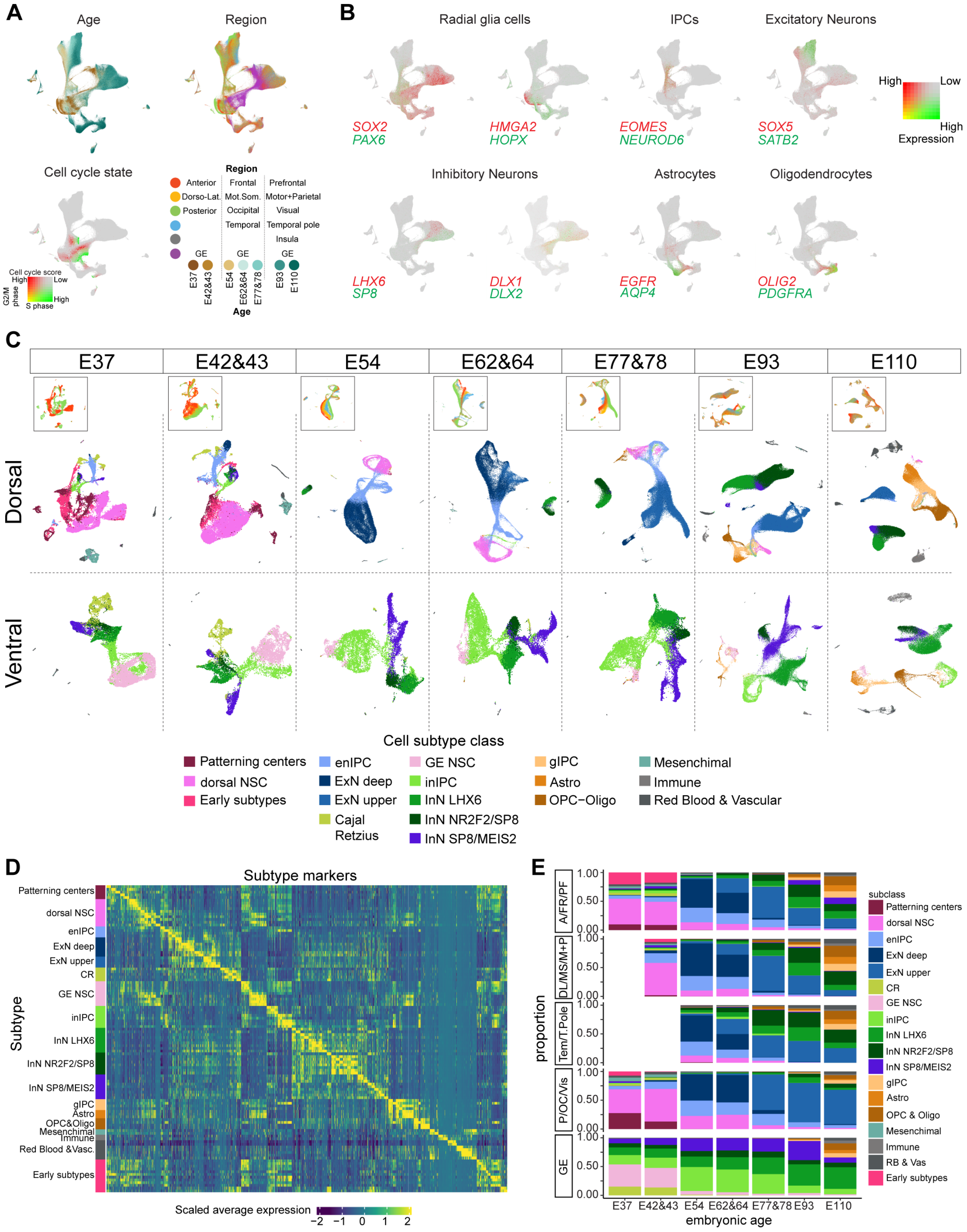


**Fig. S2. Spatiotemporal and transcriptomic overview of cell subtypes** (**A**) Age, region and cell cycle score distribution on the UMAP layout. The nomenclature of the regions across developmental time is indicated. (**B**) Expression of the major class markers on the UMAP. (**C**) Spatiotemporal dynamics of different major dorsal and ventral cell subtypes. Regions (insets) as indicated in the legend in A. (**D**) Top marker genes distinguishing different cell subtypes. **(E)** Proportion of the main cell subtypes across regions and age. Regions (left) across the embryonic ages follow the scheme in A.

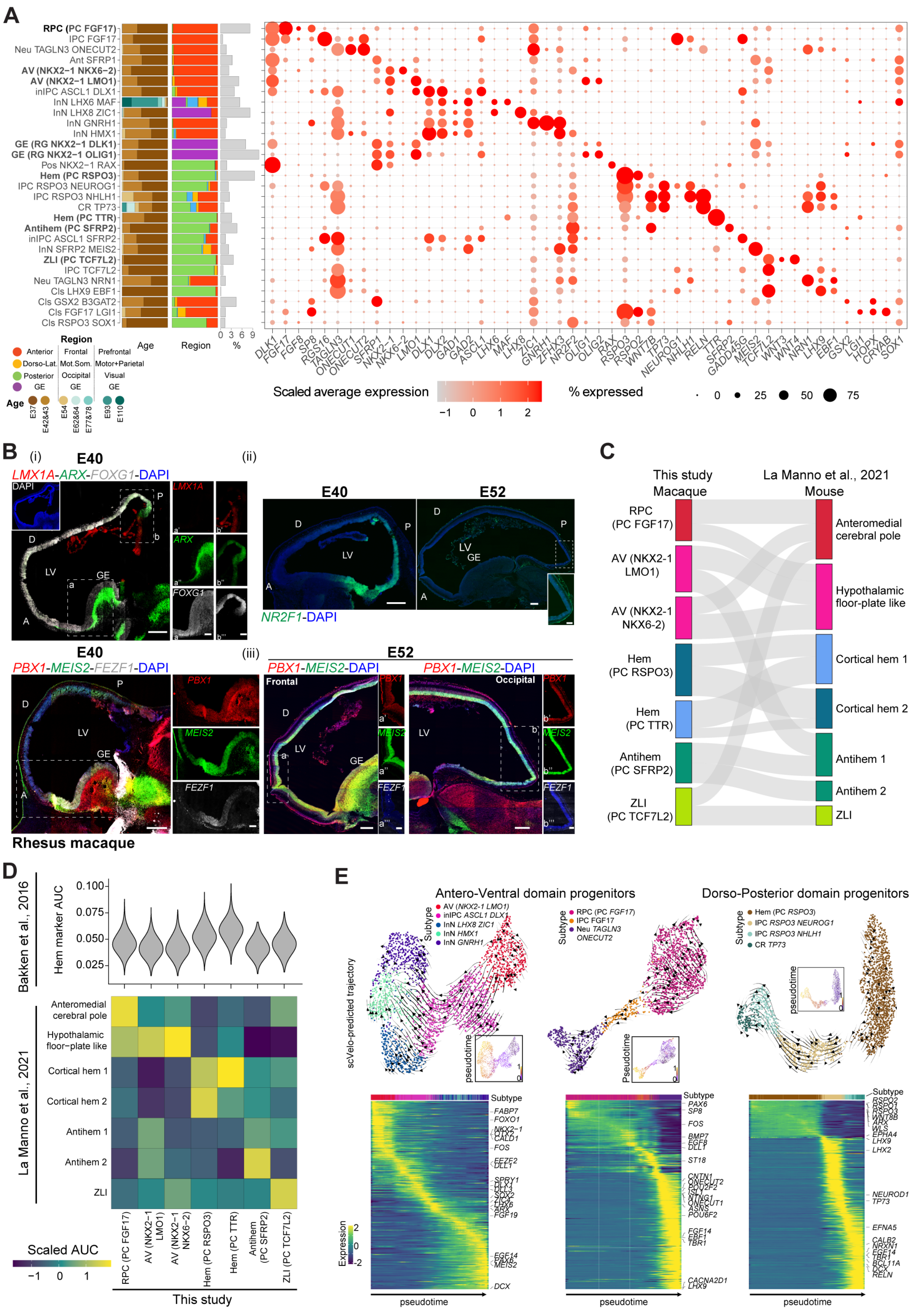
A
RPC (PC FGF17)
IPC FGF17
Neu TAGLN3 ONECUT2
Ant SFRP1
AV (NKX2−1 NKX6−2)
AV (NKX2−1 LMO1)
inIPC ASCL1 DLX1
InN LHX6 MAF
InN LHX8 ZIC1
InN GNRH1
InN HMX1
GE (RG NKX2−1 DLK1)
GE (RG NKX2−1 OLIG1)
Pos NKX2−1 RAX
Hem (PC RSPO3)
IPC RSPO3 NEUROG1
IPC RSPO3 NHLH1
CR TP73
Hem (PC TTR)
Antihem (PC SFRP2)
inIPC ASCL1 SFRP2
InN SFRP2 MEIS2
ZLI (PC TCF7L2)
IPC TCF7L2
Neu TAGLN3 NRN1
Cls LHX9 EBF1
Cls GSX2 B3GAT2
Cls FGF17 LGI1
Cls RSPO3 SOX1
Region
Anterior
Dorso-Lat.
Posterior
GE
Frontal
Mot.Som.
Occipital
GE
Prefrontal
Motor+Parietal
Visual
GE
Age
E37
E42&43
E54
E62&64
E77&78
E93
E110
Age
Region
%
0 3 6 9
Scaled average expression
% expressed
B
(i)
E40
LMX1A-ARX-FOXG1-DAPI
DAPI
(ii)
E40
E52
NR2F1-DAPI
E40
PBX1-MEIS2-FEZF1-DAPI
(iii)
E52
PBX1-MEIS2-DAPI
Frontal
Occipital
Rhesus macaque
C
This study
Macaque
La Manno et al., 2021
Mouse
RPC (PC FGF17)
AV (NKX2-1 LMO1)
AV (NKX2-1 NKX6-2)
Hem (PC RSPO3)
Hem (PC TTR)
Antihem (PC SFRP2)
ZLI (PC TCF7L2)
Anteromedial cerebral pole
Hypothalamic floor-plate like
Cortical hem 1
Cortical hem 2
Antihem 1
Antihem 2
ZLI
D
Bakken et al., 2016
Hem marker AUC
La Manno et al., 2021
Scaled AUC
This study
E
Antero-Ventral domain progenitors
Dorso-Posterior domain progenitors
scVelo-predicted trajectory
pseudotime
Subtype
Expression

**Fig. S3. Molecular characterization of the monkey putative telencephalic organizer domains** (**A**) Left bar plots: age, region composition and cluster proportions of the early cell subtypes. Region identity across the developmental age is indicated on bottom left. Right dot plot: expression level (color gradient) of and percentage of cells (dot size) expressing subtype markers. (**B**) Expression of *LMX1A*, *ARX*, *FOXG1* (top), *PBX1*, *MEIS2*, FEZF1 (bottom), and NR2F1 (right) detected by RNAscope in monkey E40 and E52 sagittal brain sections. Panoramic scans are shown. Higher magnification from the dashed areas in the antero-ventral (A-V) and posterior-caudal (D-C) domains are shown. Scale bars: 500 μm (panoramic view) and 200 μm (zoom-in view) (**C**) Sankey plot illustrating the transcriptomic similarity (Pearson correlation coefficients) between macaque (this dataset) and mouse (*22*) brain organizer subtypes. A cutoff was set at the 0.8 quantile of all similarity scores to remove putative background signals. (**D**) Top: AUC score assessing the enrichment of cortical hem markers generated in (*12*). Bottom: Average AUC scores measuring the enrichment of mouse patterning center subtype markers (*22*) in macaques. (**E**) Top: RNA velocity and pseudotime predicting the lineage progression of the antero ventral progenitors *FGF17*$^{+}$ and *NKX2-1*$^{+}$ (left) and the *RSPO3*$^{+}$ progenitors (right). Bottom: gene expression cascades along the lineages. FR: frontal; OC: occipital; D: dorsal; V: ventral; GE: ganglionic eminence; VZ: ventricular zone; LV: lateral ventricle.

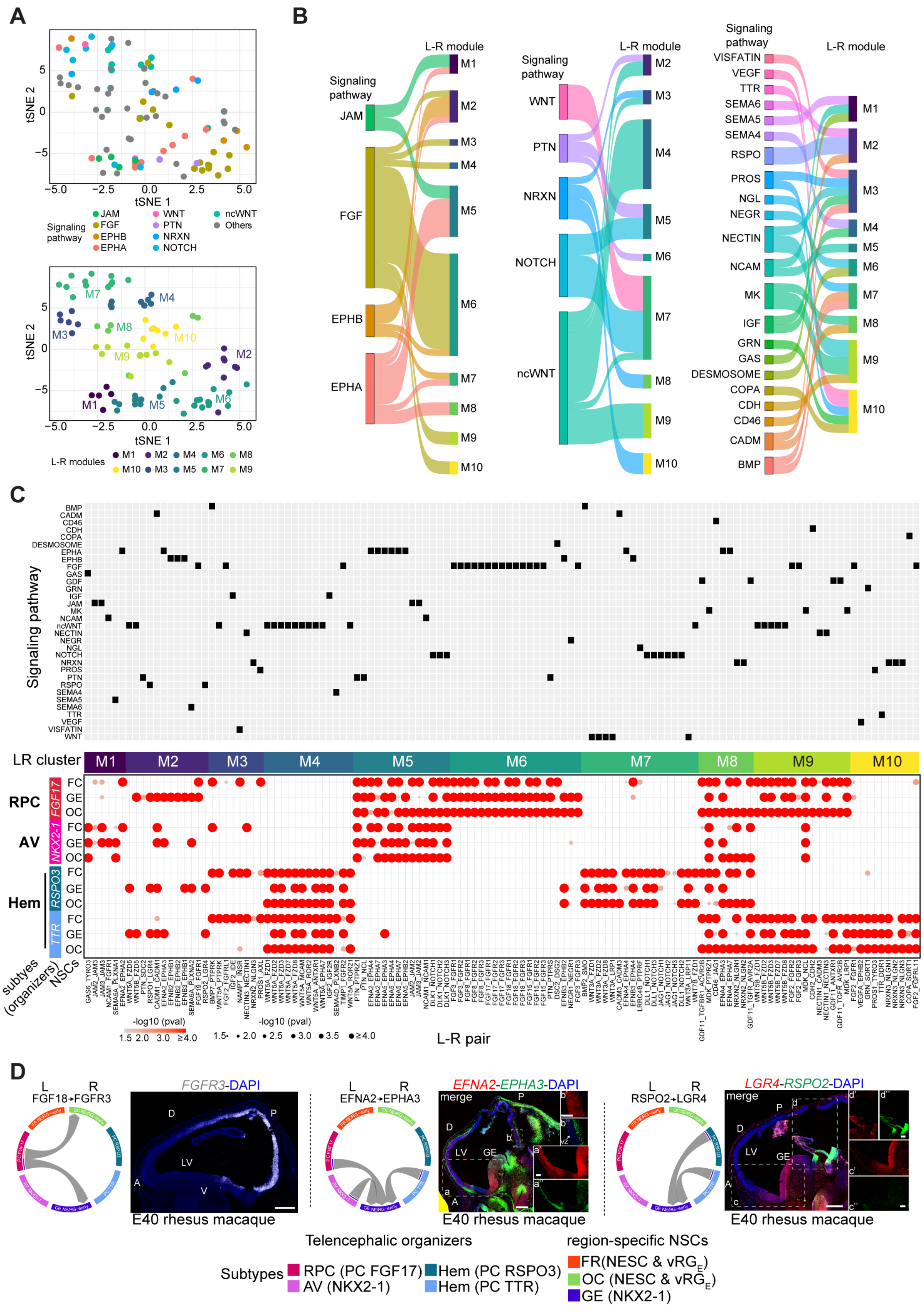
A
B
C
D
tSNE 1
tSNE 2
Signaling pathway
L-R modules
L-R module
Signaling pathway
LR cluster
RPC
AV
Hem
NSCs
subtypes (organizers)
L-R pair
-log10 (pval)
FGF18→FGFR3
EFNA2→EPHA3
RSPO2→LGR4
FGFR3-DAPI
EFNA2-EPHA3-DAPI
LGR4-RSPO2-DAPI
merge
E40 rhesus macaque
Telencephalic organizers
region-specific NSCs
Subtypes
RPC (PC FGF17)
AV (NKX2-1)
Hem (PC RSPO3)
Hem (PC TTR)
FR(NESC & vRGE)
OC (NESC & vRGE)
GE (NKX2-1)

**Fig. S4. Predicted cell-cell communications between putative telencephalic organizer domains and early radial glia cells** (**A**) t-distributed stochastic neighbor embedding (tSNE) plots visualizing the signaling pathways and the co-expression modules (M1-M10) of all the ligand (L)-receptor (R) pairs (dots). Signaling pathways with low numbers of L-R pairs were annotated as "others". (**B**) Sankey plot showing the L-R pair-mediated cell-cell interaction patterns (represented by modules, M1-M10) of all the signaling pathways. (**C**) Top: signaling pathway annotation for each L-R pair. Bottom: interaction patterns of all the L-R pairs between organizer subtypes (antero-ventral including RPC and Hem) and E37-43 NSC subtypes (NESCs and $vRG_E$) of frontal (FR), ganglionic eminence (GE) and occipital (OC) telencehalic regions organized by modules. (**D**) Circular plots showing the interaction pattern of the selected L-R pairs and RNAscope of E40 rhesus macaque brain tissue sagittal sections. Panoramic scans are shown. Higher magnifications from the dashed area are shown. The arrows show the interaction direction from the ligand (L)- to the receptor (R)-expressing cells (organizer domains and FR, OC or GE RG cells, respectively). Expression of the receptor *FGFR3* (left), the Ephrin signaling component *EFNA2* (ligand) and *EPHA3* (receptor) (middle), the WNT signaling members *RSPO2* (ligand) and *LGR4* (receptor) is shown (right).

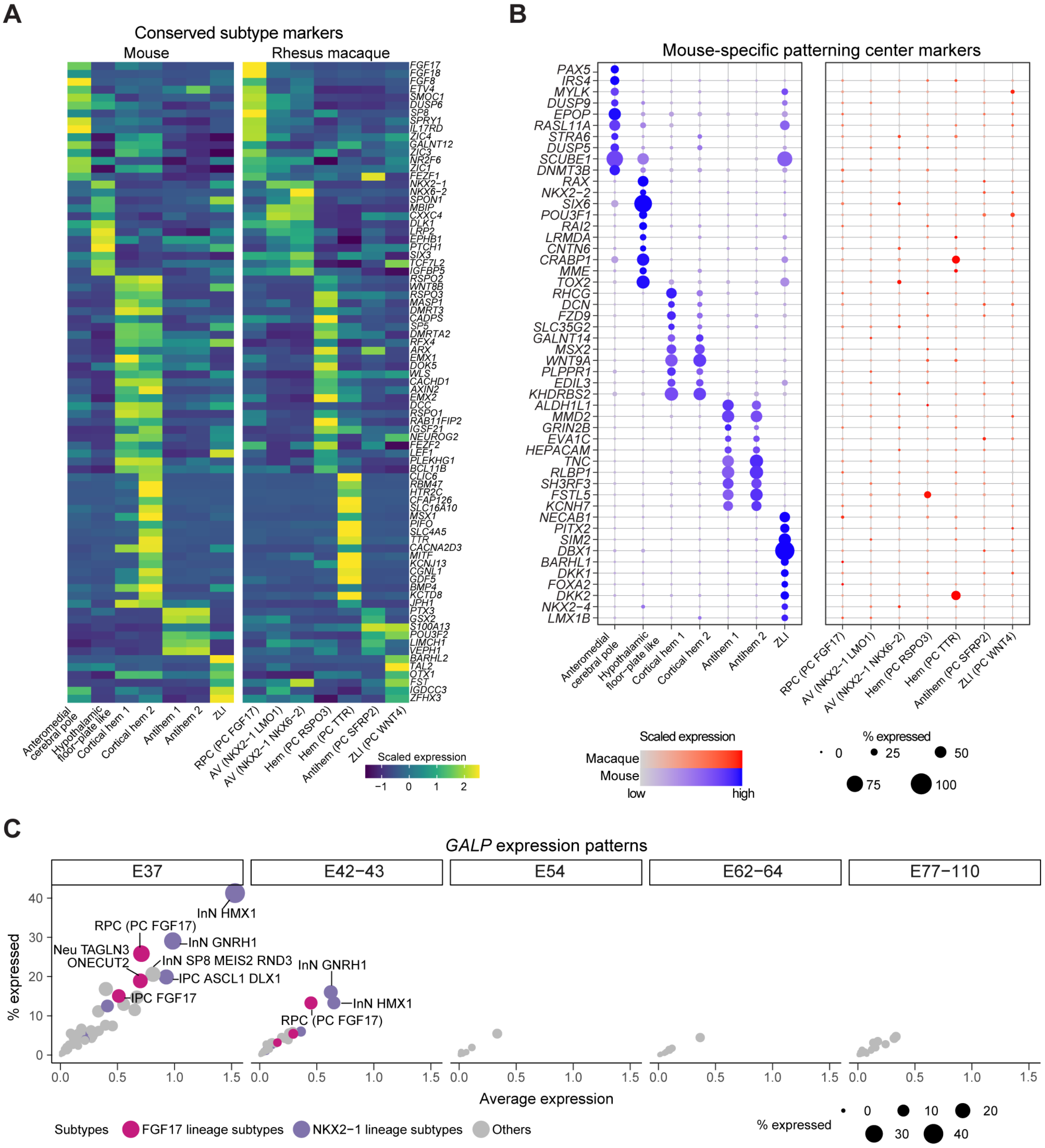


**Fig. S5. Conserved and divergent transcriptomic features between mouse and macaque brain organizer domain subtypes (A)** Genes displaying conserved expression between mouse and macaque brain organizer domain subtypes. (**B**) Expression of the top 10 genes showing mouse-enriched expression in each subtype. Dot color indicates scaled expression for each specie; dot size indicates the percentage of cells expressing a gene. (**C**) *GALP* expression across cell subtypes and time. The dot size represents percentage of cells expressing *GALP*. Top 7 cell subtypes in each age group showing a percentage no smaller than 10% and an average log-transformed expression value no less than 0.4 of *GALP* expression are colored and labeled.

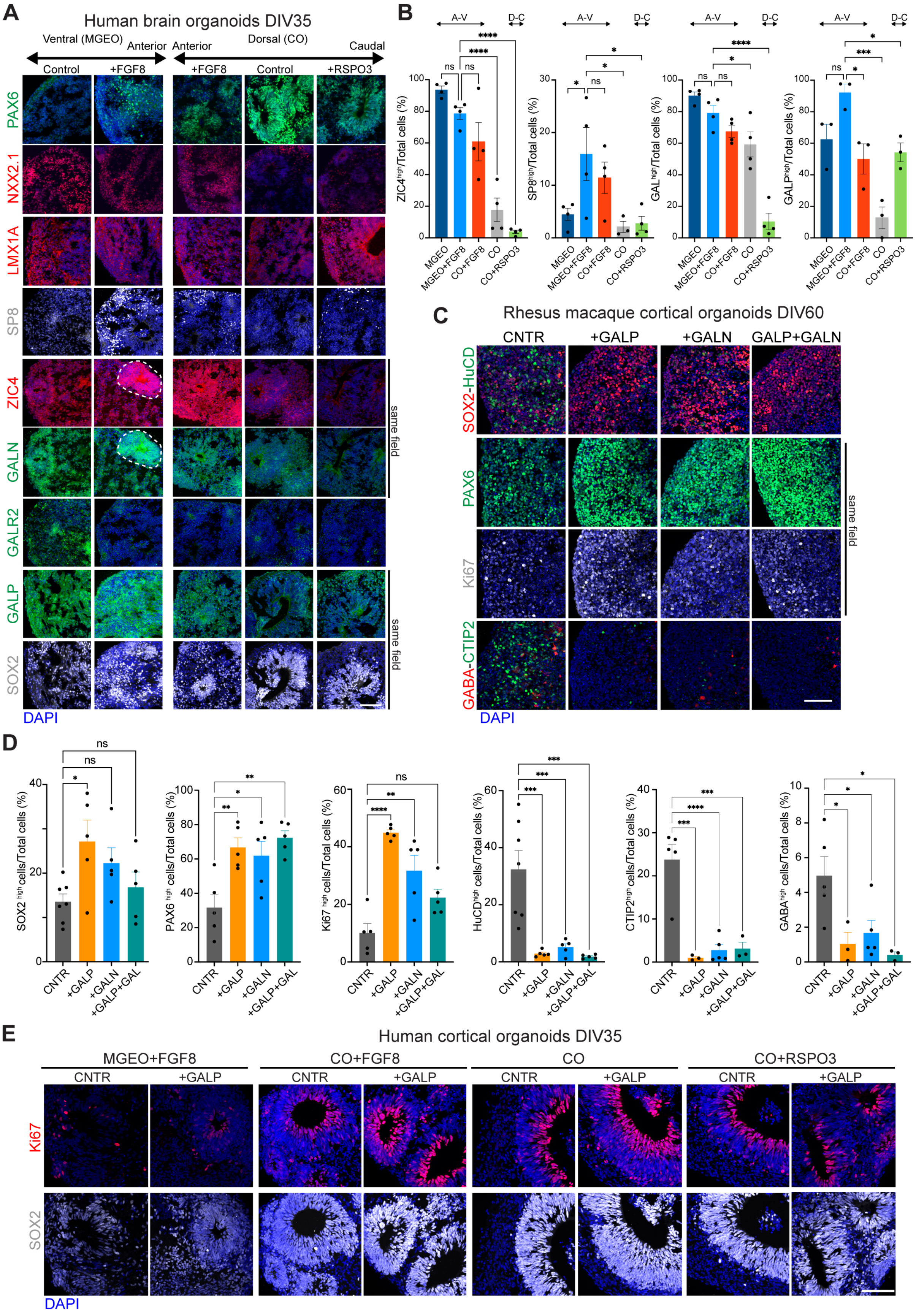
A
Human brain organoids DIV35
Ventral (MGEO)
Anterior
Anterior
Dorsal (CO)
Caudal
Control
+FGF8
+FGF8
Control
+RSPO3
PAX6
NXX2.1
LMX1A
SP8
ZIC4
GALN
GALR2
GALP
SOX2
same field
same field
DAPI
B
A-V
D-C
MGEO
MGEO+FGF8
CO+FGF8
CO
CO+RSPO3
C
Rhesus macaque cortical organoids DIV60
CNTR
+GALP
+GALN
GALP+GALN
SOX2-HuCD
PAX6
Ki67
GABA-CTIP2
same field
DAPI
D
CNTR
+GALP
+GALN
+GALP+GAL
E
Human cortical organoids DIV35
MGEO+FGF8
CO+FGF8
CO
CO+RSPO3
CNTR
+GALP
Ki67
SOX2
DAPI

**Fig. S6. Region-specific features of the monkey developing telencephalon recapitulated *in vitro* by brain organoids (A)** Cortical (hCO) and medial ganglionic eminence (hMGEO) organoids generated from human iPSCs and differentiated up to day *in vitro* (DIV) 35. hCOs were exposed to dorso-caudal (RSPO3) or antero-ventral (FGF8) patterning signals or to control conditions. Human MGEO were exposed to FGF8 or control conditions. Immunofluorescence images of the expression of the indicated markers. Scale bar 100 µm. **(B)** High-throughput image analysis showing the percentage of cells expressing the indicated proteins in the different culture conditions ± SD, One-way ANOVA, Dunnett's multiple comparisons. Significance is indicated. A-V: antero-ventral; D-C: dorso-caudal (**C**) Immunohistochemistry of the indicated genes in macaque iPSC-derived cortical organoids, control or exposed to GALP, GAL or both. Scale bar 100 µm. **(D)** Image analysis of at least 5 different fields showing percentage of cells expressing the indicated markers in the different conditions ± SD, One-way ANOVA, Dunnett's multiple comparison (*: $p < 0.1$; **: $p < 0.01$; ***: $p < 0.001$; ****: $p < 0.0001$; ns: not significant). (**E**) Immunohistochemistry of SOX2 and Ki67 from sections of human CO or MGEO exposed or not to patterning ligands (FGF8 or RSPO3) plus/minus GALP. Scale bar 100µm. CO+FGF8 panel and quantification shown in Fig. 3F. CNTR: control.

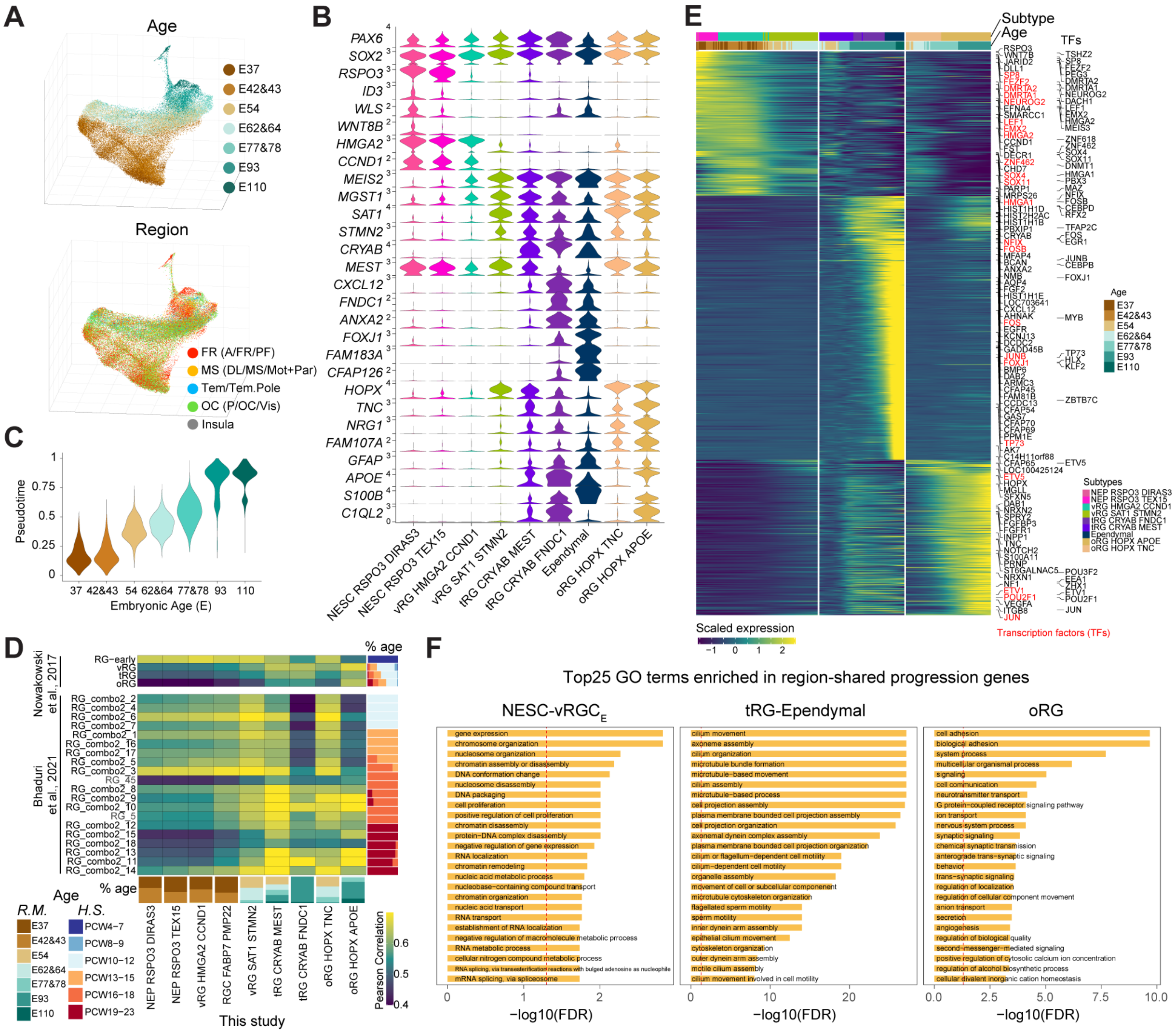

**Fig. S7. Macaque cortical neural stem cell subtype characterization (A)** Three-dimensional UMAP showing the age and region distribution of the NSC subtypes. The regions from each domain follow the nomenclature across the time shown in Fig. 1. (**B**) Violin plot showing expression patterns of NSC markers. (**C**) Pseudotime representing cortical NSC progression highly correlated with developmental age. (**D**) Pearson correlation measuring transcriptomic similarity between macaque and human (*14, 42*) RG subtypes, indicating higher similarity between the RG cells defined in (*14*) and those identified after E54 in this study. (**E**) Shared gene expression cascades across cortical areas along the NSC lineage progression. Transcriptin factors and chromatin remodeling genes are highligthed (**F**) Gene Ontology enrichment of the regional-shared genes in panel E. Top 25 significant terms are shown.

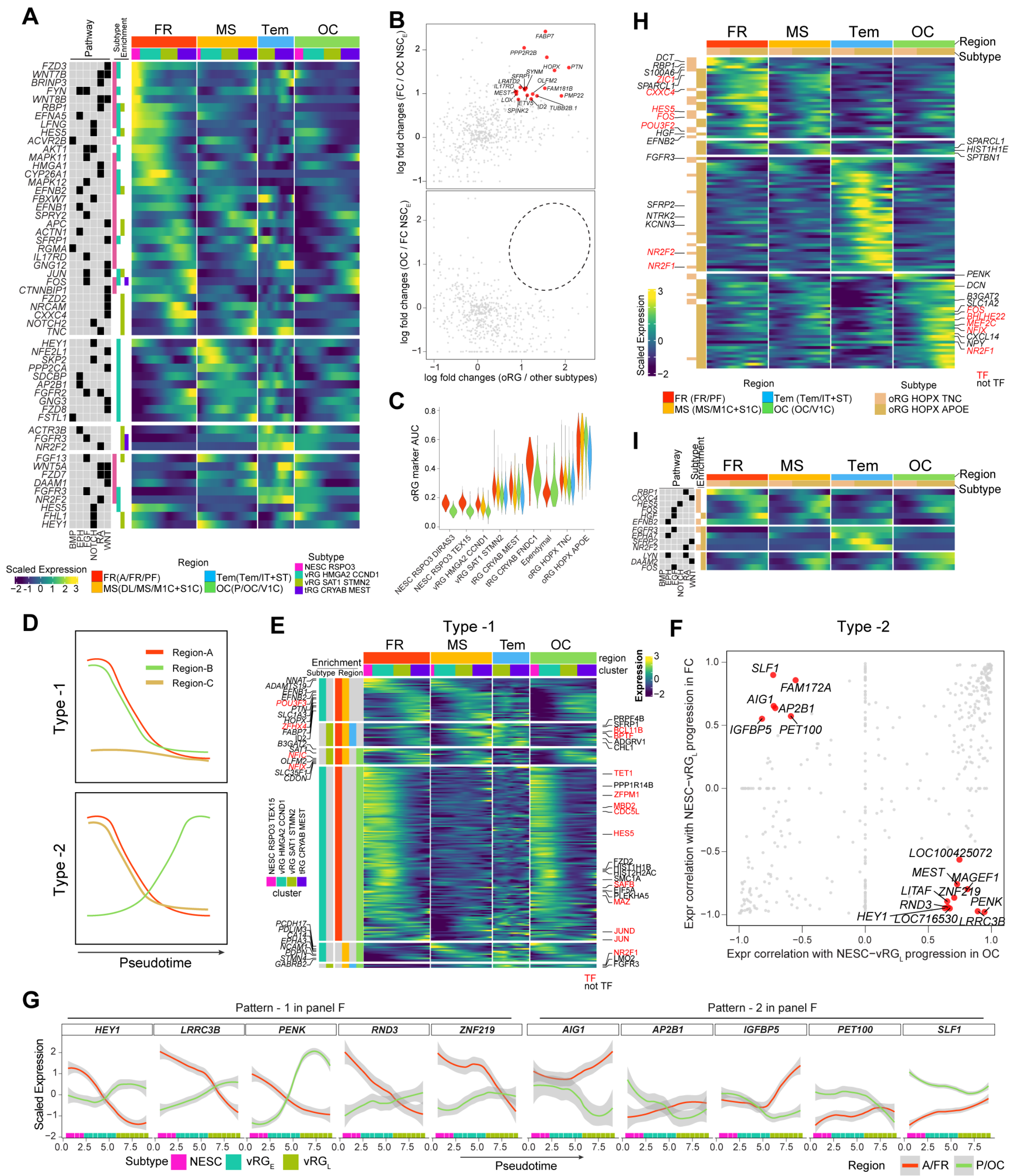

**Fig. S8. Divergent transcriptomic signatures underlying neural stem cell progression across cortical regions** (**A**) Expression of genes involved in the selected signaling pathways and displaying region-specificity along the progression of ventricular RG cells. Enrichment in a subtype is shown on the left side. The legend indicates the regions considered across the time from each domain, as indicated in Fig. 1. (**B**) Top: expression enrichment of regionally divergent genes in early frontal vs occipital NSCs, and in occipital oRG vs other occipital NSCs. Genes enriched in both NSCs and oRGCs simultaneously are shown as red dots and labeled. Bottom: Same as in the above panel, except the y axis shows the enrichment in early occipital versus frontal NSCs. Unlike frontal early NSC and oRG cells, no genes were found to be enriched in early occipital NSC- and oRG-enrichment simultaneously. (**C**) AUC scores assessing the enrichment of the occipitcal oRG subtype markers across all telencephalic regions collected and NSC subtypes. (**D**) Diagrams illustrating two models of region-specific genes employed by more than one region. (**E**) Expression cascades of genes enriched in multiple, but not all, regions displaying similar expression dynamics in the enriched regions. Subtype and regions employing same genes are indicated on the top of the heatmap. The regions considered are the same as in A. Transcription factor genes are labeled as red text. (**F**) Gene expression correlation of NSC progression in the frontal versus occipital region. Genes showing expression patterns positively correlated with occipital VZ NSC progression but negatively correlated with frontal VZ NSC progression, or viceversa, were highlighted and labeled. (**G**) Expression patterns of some representative genes highlighted in panel F. (**H**) Region-specific gene expression cascades along the progression of oRG cells. Cell subtype and region information are shown on the top and in the legend. The regions follow the nomenclature across developmental time shown in Fig. 1. (**I**) Same as in H, genes involved in the selected signaling pathways are shown.

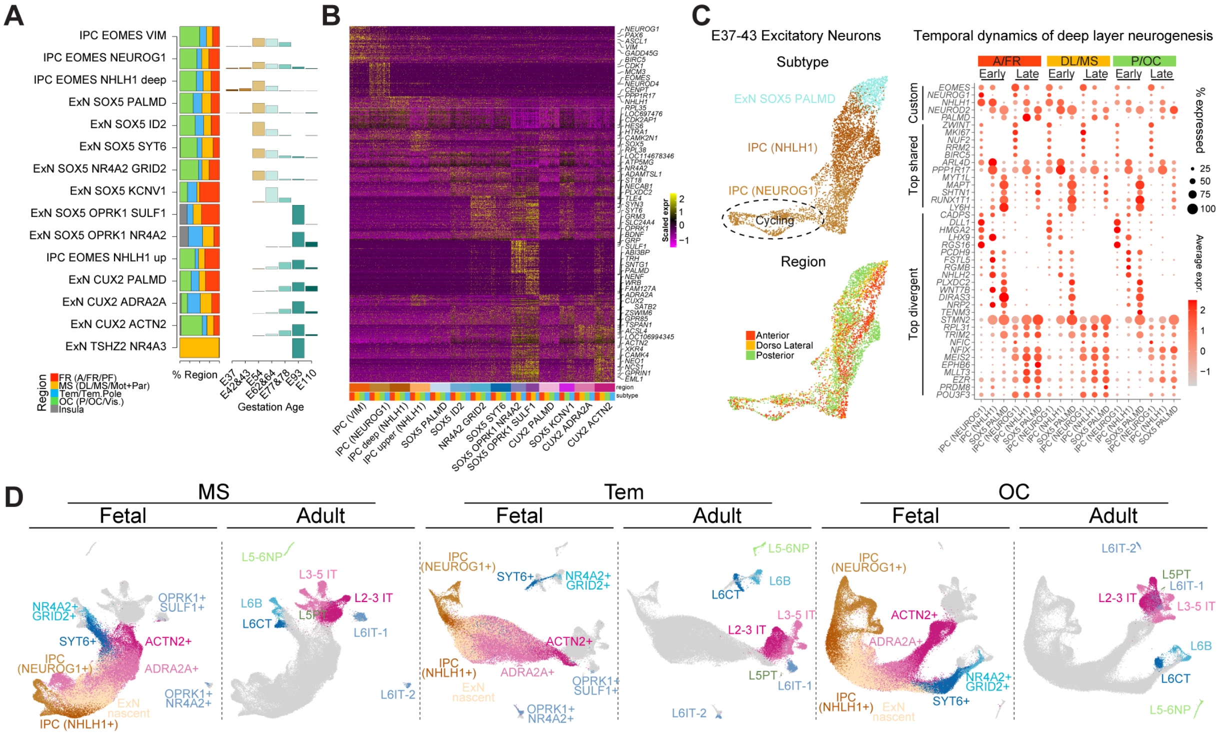

**Fig. S9. Characterization of excitatory neuron differentiation and maturation (A)** Age and region proportion of each cell subtype in the excitatory neuron lineage. Notice multiple subtypes of IPCs, deep and upper layer cortical neurons with different appearance across the time. The legend indicates the regions across the time from each domain, as indicated in Fig. 1. (**B**) Expression of regionally-shared markers of IPC and excitatory neuron subtypes. Region information are indicated with colors on the bottom of the heatmap. Colors of the regions as indicated in A. (**C**) Left: Early emergence (E37-43) of deep layer excitatory neurons. Right: These early-wave of excitatory neurons share signatures with the deep layer excitatory neurons generated at E54-64, although they also exhibit many divergent transcriptomic changes. The regions are indicated based on the developmental time, as indicated in Fig. 1. (**D**) Pairwise transcriptomic integration between the fetal excitatory neuron lineage of a given region analyzed in this study and the excitatory neurons from the adult macaque dorsolateral prefrontal cortex (*48*).

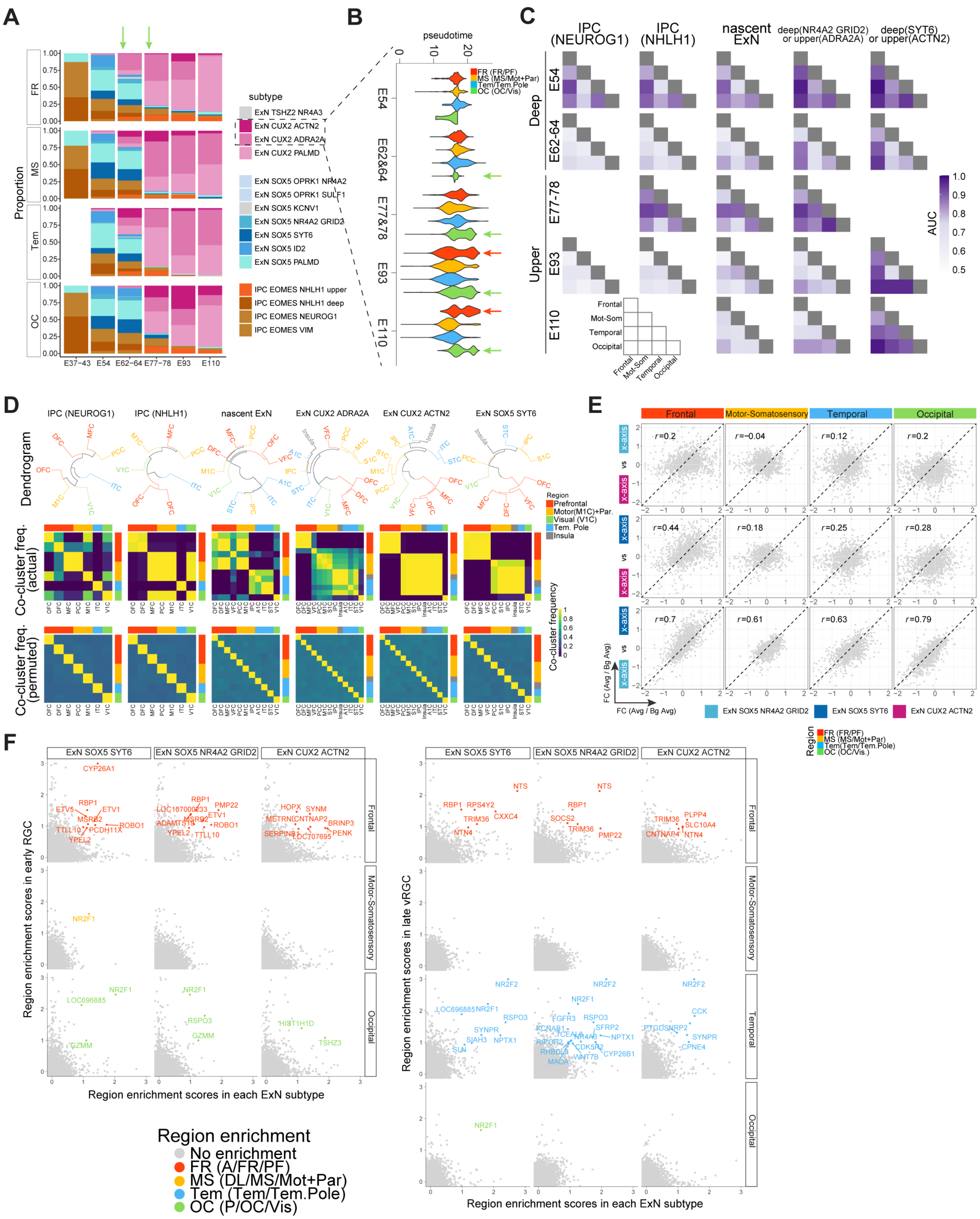


**Fig. S10. Molecular programs underlying the diversifications of neurogenesis across cortical regions (A-B)** Composition of IPC and excitatory neuron subtypes in each cortical region across the analyzed developing timepoints (A). Upper layer excitatory neurons emerge around E62-64 in prefrontal, dorsomedial and temporal, but not in the occipital regions. However, these neurons mature faster in the occipital cortex at E77-78, as depicted by the pseudotime scores (B). Their maturation state remain higher in prefrontal and occipital regions at E93-E110 (B). (**C**) Increasing transcriptomic divergence, represented by AUC scores calculated by Augur (material and methods), across cortical regions in the subtypes along the differentiation and maturation lineage of the excitatory neurons. Subtypes without sufficient cells in all cortical regions were not included. (**D**) Top: Hierarchial clustering of cortical regions for each cell subtype using highly variable genes. Middle: Co-cluster frequencies of pairwise regions for each subtype using bootstrap replicates of the highly variable genes. Bottom: Co-cluster frequencies of pairwise regions using permuted data. Although regional divergene emerge at IPC cells, these differences were not sufficient to distinguish regions with a pattern resembling their anatomical organizations, which does not occur until the generation of more mature excitatory neuron subtypes (ExN *CUX2 ACTN2* and ExN *SOX5 SYT6*). When for certain cell types there were not sufficient cells in certain regions, they were not included in the analysis. (**E**) Correlation of regional enrichment between deep and upper excitatory neuron subtypes for genes divergently expressed across regions using the peak stages of deep (E54-64) and upper layer (E93-110) neurogenesis. Here the regional enrichment is represented by the log fold changes of average expression in a given region versus the background regions. The Pearson correlation coefficients were listed on the top left of each panel. Notice high Pearson correlation coefficients between the late deep layer excitatory neuron subtypes indicating overlapping of the region-specific signatures (bottom panels). The same analysis was not performed for the upper layer neurons because only one late-phase differentiated subtype was identified. (**F**) Dotplot showing the overlap of region-specific signatures between early or late NSC subtypes and excitatory neuron subtypes. Shared region-specific genes are colored by regions and labeled. In each panel, the regions across the time from each domain follow the nomenclature indicated in Fig. 1.

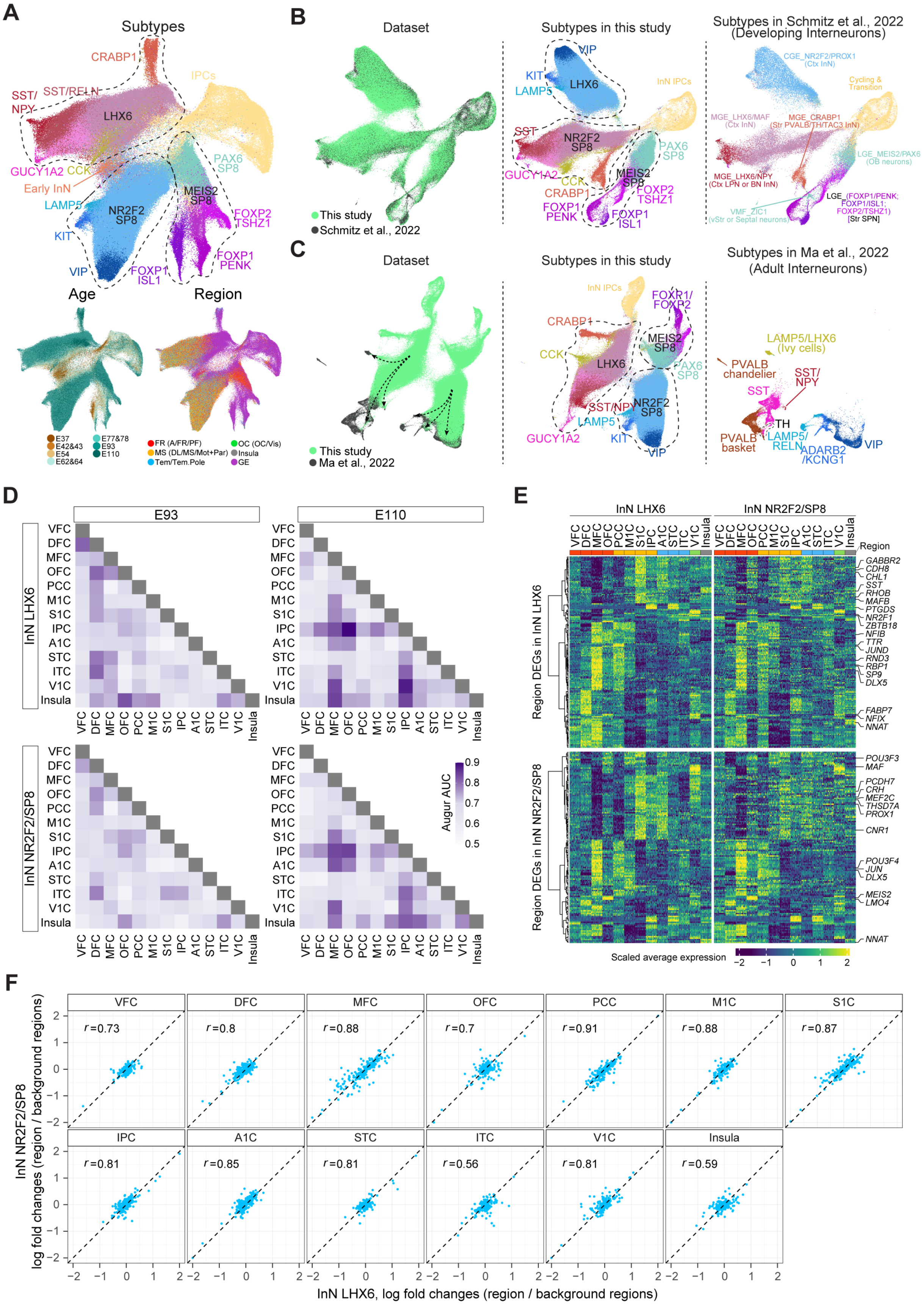
A
Subtypes
Age
Region
B
Dataset
Subtypes in this study
Subtypes in Schmitz et al., 2022 (Developing Interneurons)
This study
Schmitz et al., 2022
C
Dataset
Subtypes in this study
Subtypes in Ma et al., 2022 (Adult Interneurons)
This study
Ma et al., 2022
D
E93
E110
InN LHX6
InN NR2F2/SP8
Augur AUC
E
InN LHX6
InN NR2F2/SP8
Region
Region DEGs in InN LHX6
Region DEGs in InN NR2F2/SP8
Scaled average expression
F
InN NR2F2/SP8 log fold changes (region / background regions)
InN LHX6, log fold changes (region / background regions)

**Fig. S11. Limited transcriptomic divergences of the cortical inhibitory GABAergic interneurons across development (A)** UMAP plots showing cortical inhibitory neurons subtypes, and their region and age distribution. For conciseness, only the key markers from subtype names were labeled. The legend indicates the regions across the time from each domain, as indicated in Fig. 1. **(B)** Transcriptomic integration between this data and an independent macaque developing inhibitory neuron data (*52*)**.** The two datasets align well, however more heterogeneity emerges in this study. (**C**) Transcriptomic integration between fetal (this study) and adult (*48*) macaque inhibitory neurons. Adult cell subtypes locate at the tips of each lineages (indicated by arrows). The majority of the adult inhibitory neuron types matched to fetal inhibitory neuron lineages. (**D**) AUC scores calculated by Augur representing transcriptomic separability of inhibitory neurons between each pair of regions. The AUC scores are largely close to 0.5, suggesting the same inhibitory neuron types are quite transcriptomically homogenous across cortical regions. However, certain cortical regions resulted more transcriptomically unique. This uniqueness displayed similar patterns in *LHX6* and *NR2F2/SP8* inhibitory neurons lineages, but distinct dynamics between E93 and E110 samples. (**E**) Hierachical clustering of differentially expressed genes (DEGs) across cortical regions in *LHX6* (top) and *NR2F2/SP8* (bottom) cortical inhibitory neurons. For each set of DEGs, their expression were visualized in both *LHX6* (left) and *NR2F2/SP8* (right) inhibitory neurons, which exihibited almost identical expression patterns. (**F**) Correlation of regional enrichment between *LHX6* (x axis) and *NR2F2/SP8* (y axis) cortical inhibitory neurons. Here the regional enrichment is represented by the log fold changes of average expression in a given region versus the background regions. The Pearson correlation coefficients were listed on the top left of each panel.

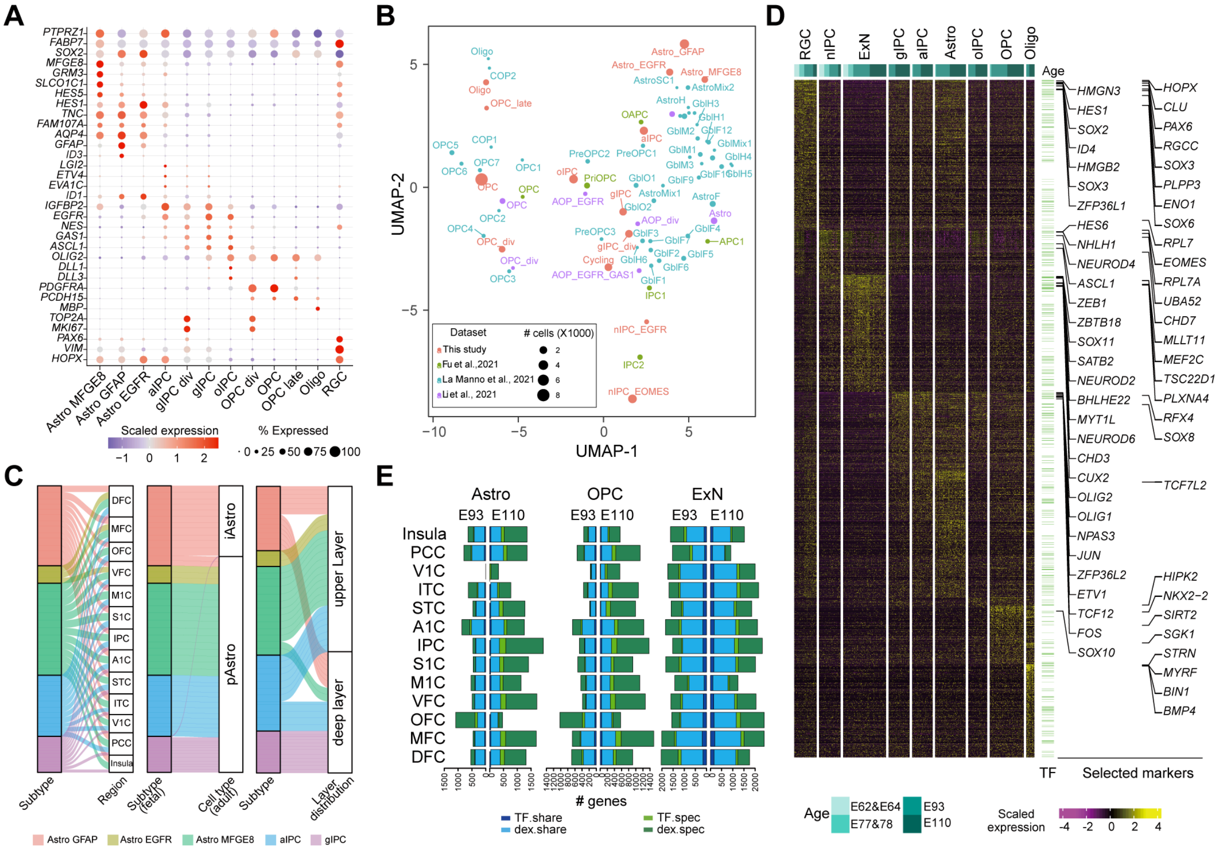

**Fig. S12. Cortical arealization during gliogenesis** (**A**) Expression of curated genes specific for RG cells and glia cell subtypes. The color strength and circle size scale the normalized gene expression and the percentages of expressed cells. (**B**) UMAP plot showing data integration of the current study with three other studies (*22, 57, 58*). The circle sizes scale the number of cells identified in each cell type, and the positions represent the genomic centers of each cell type in accord with the embedded UMAP plots. (**C**) Alluvial plots showing regional distribution of the astrocyte subtypes defined in this study compared with the astrocytes reported in two other studies: (*48*) and (*59*)). **(D)** Expression of detected genes specified for RG cells, nIPC, ExN, gIPC, aIPC, Astro, oIPC, OPC and Oligo. Transcriptional factors are highlighted beside the heatmap (green horizontal lines), and the selected markers are labeled on the right. (**E**) Bar plots showing the numbers of genes and transcription factors (TFs) identified as differentially expressed across regions in astrocytes (Astro), oligodendrocyte precursors (OPC), and excitatory neurons (ExN) at E93 and E110. Comparisons between E93 and E110 are conducted to reveal the temporal specification of the cells.

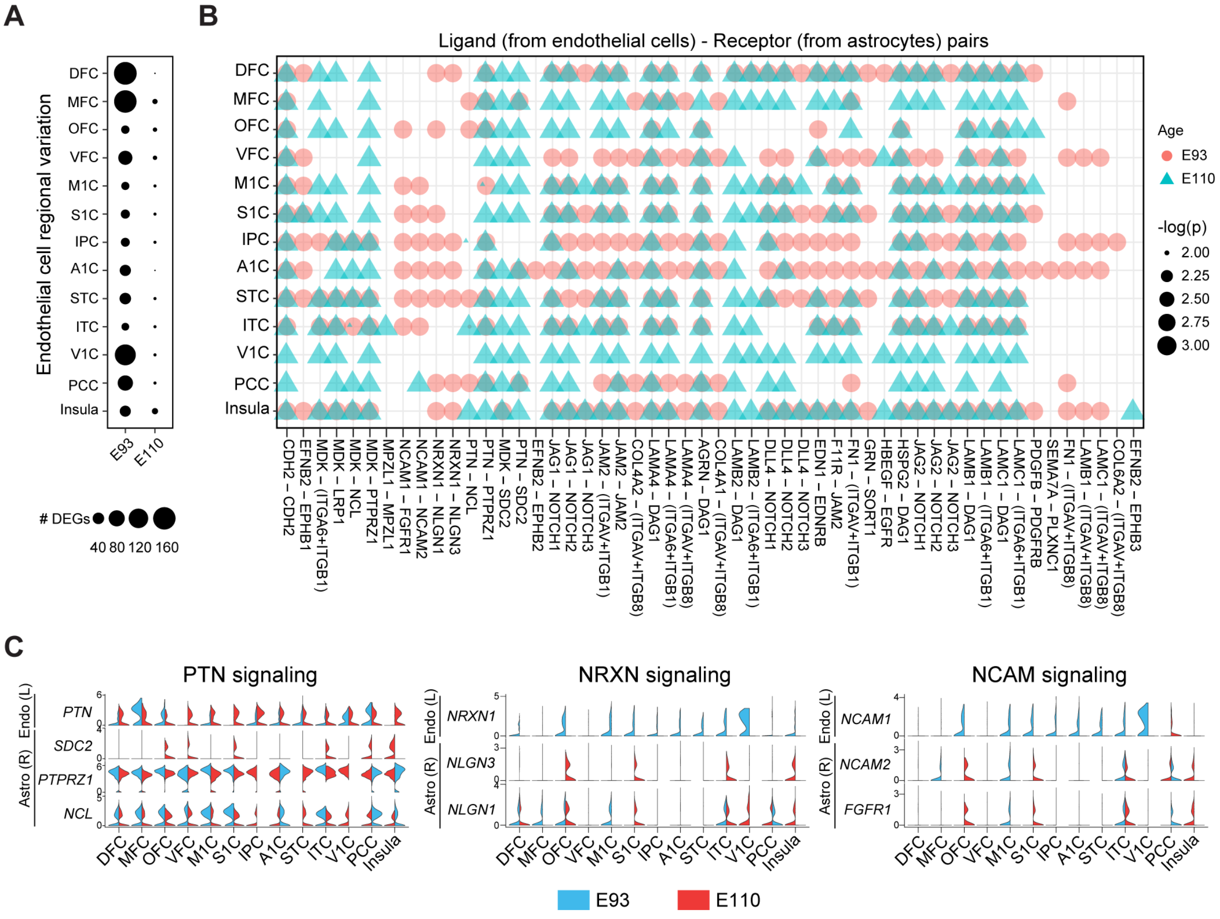


**Fig. S13. Cell-cell interactions between endothelial cells and astrocytes** (**A**) Dot plots representing the number of genes identified as regional differential expression in endothelial cells. (**B**) Interactions between ligand (L) from endothelial cells and receptor (R) genes from astrocytes at E93 (red circle) and E110 (blue triangle). The circle size scales the negative log2 transformed p-values. (**C**) Expression of ligand-receptor paired genes, where the ligand (L) is expressed by the by endothelial cells (Endo) and the receptor (R) by the astrocytes (Astro), involved in PTN, NRXN and NCAM signaling pathways across the regions at E93 and E110.

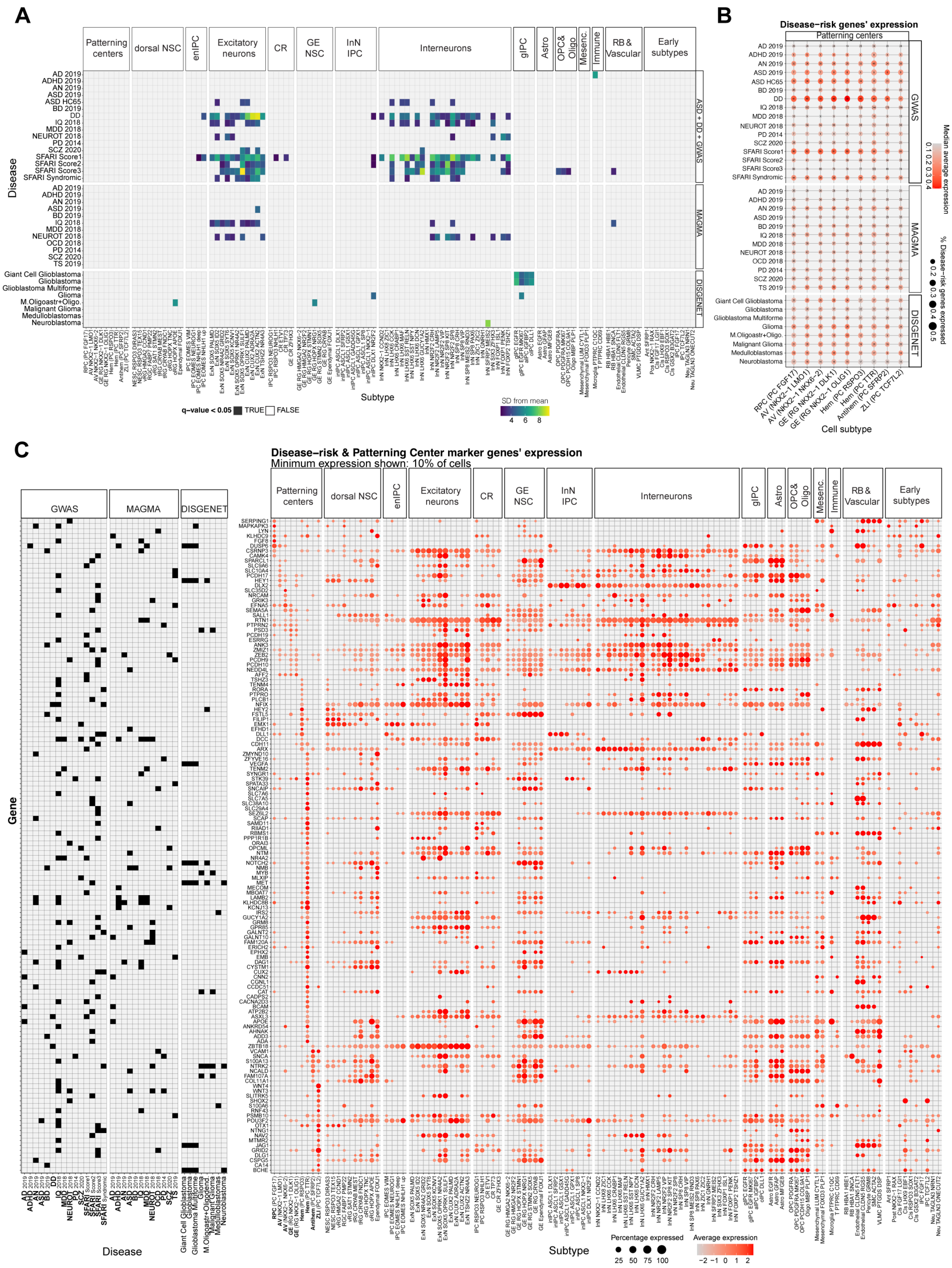
A
Disease
Subtype
q-value < 0.05 TRUE FALSE
SD from mean
4 6 8
Patterning centers
dorsal NSC
enIPC
Excitatory neurons
CR
GE NSC
InN IPC
Interneurons
gIPC
Astro
OPC& Oligo
Mesenc.
Immune
RB & Vascular
Early subtypes
ASD + DD + GWAS
MAGMA
DISGENET
B
Disease-risk genes' expression
Patterning centers
GWAS
MAGMA
DISGENET
Cell subtype
Median average expression
% Disease-risk genes expressed
C
Disease-risk & Patterning Center marker genes' expression
Minimum expression shown: 10% of cells
GWAS
MAGMA
DISGENET
Gene
Disease
Subtype
Percentage expressed
25 50 75 100
Average expression
−2 −1 0 1 2

**Fig. S14. Expression of brain disease-risk genes in in early development** (**A**) Heatmap representation of Expression Weighted Celltype Enrichment (EWCE) results. Only significant enrichment results (q-value < 0.05) are shown. The color scale represents the number of standard deviations over the expected mean of the corresponding enrichment result. (**B**) Dot-plot representation of gene expression for disease-associated genes in the organizer domain subtypes. Dot size represents the proportion of disease risk genes expressed in the given subtype. Dot color indicates the median gene average expression of the disease genes. The number inside each dot represents the number of genes from the given disease gene list that are expressed in the subtype. (**C**) Dot-plot representation of scaled gene expression for the organizer domain cells' marker genes linked to diseases across all subtypes. The subtype marker genes shown are expressed in more than 10% of cells of their respective organizer subtype and they are expressed in less than 10% of other clusters cells on average. The associations between genes and diseases are shown by the panel on the left, indicated by a black square.

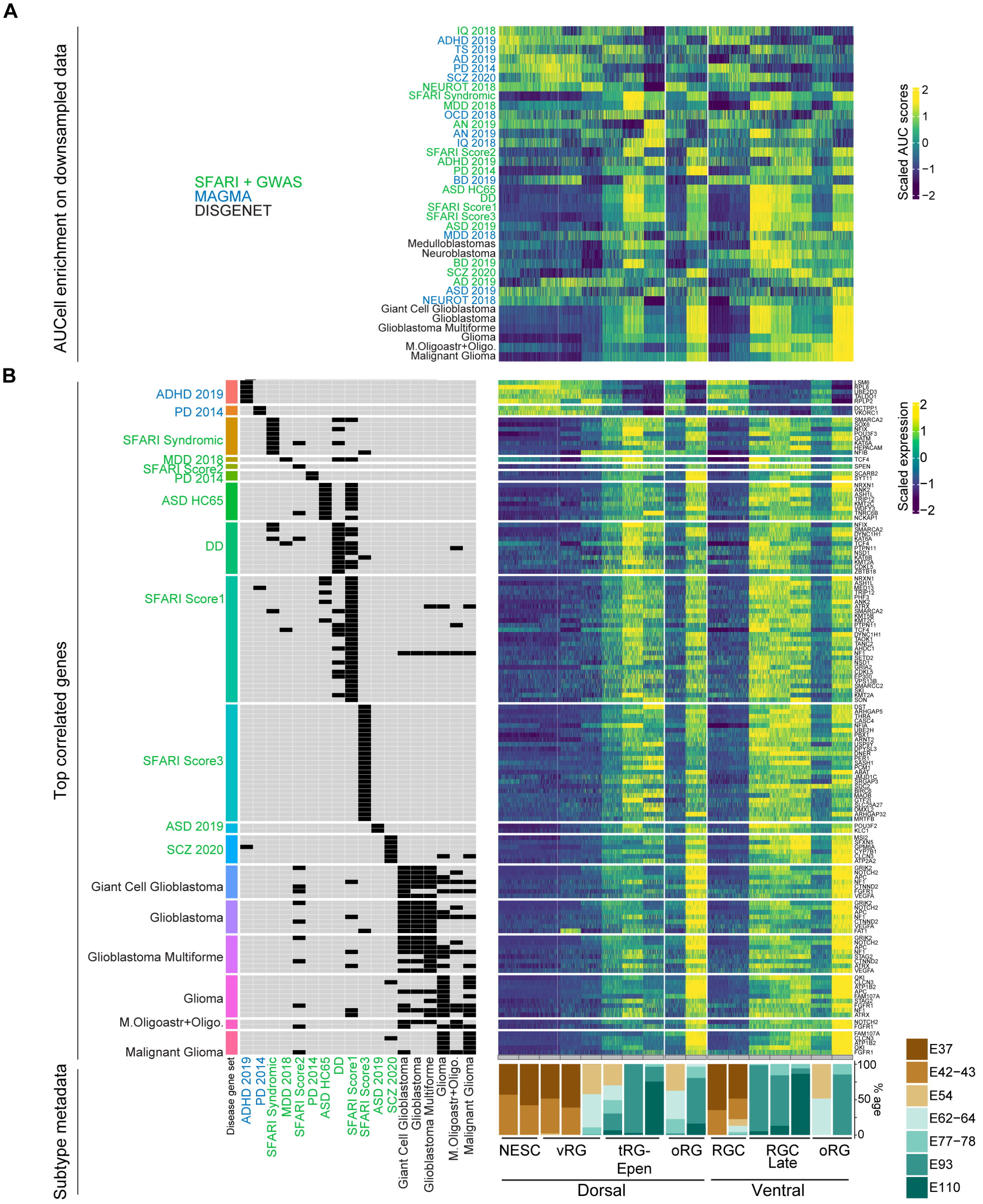


**Fig. S15. Enrichment of the disease risk genes along the progression of dorsal and ventral NSCs**

(**A**) Scaled AUC scores recapitulating the expression enrichment patterns of the disease genesets across NSC subtypes in the downsampled data. Genesets are arranged in the same order as that in Fig. 7D. (**B**) For each disease geneset, genes highly correlated (Pearson correlation coefficients > 0.7) with the geneset enrichment pattern were selected and visualized for expression.

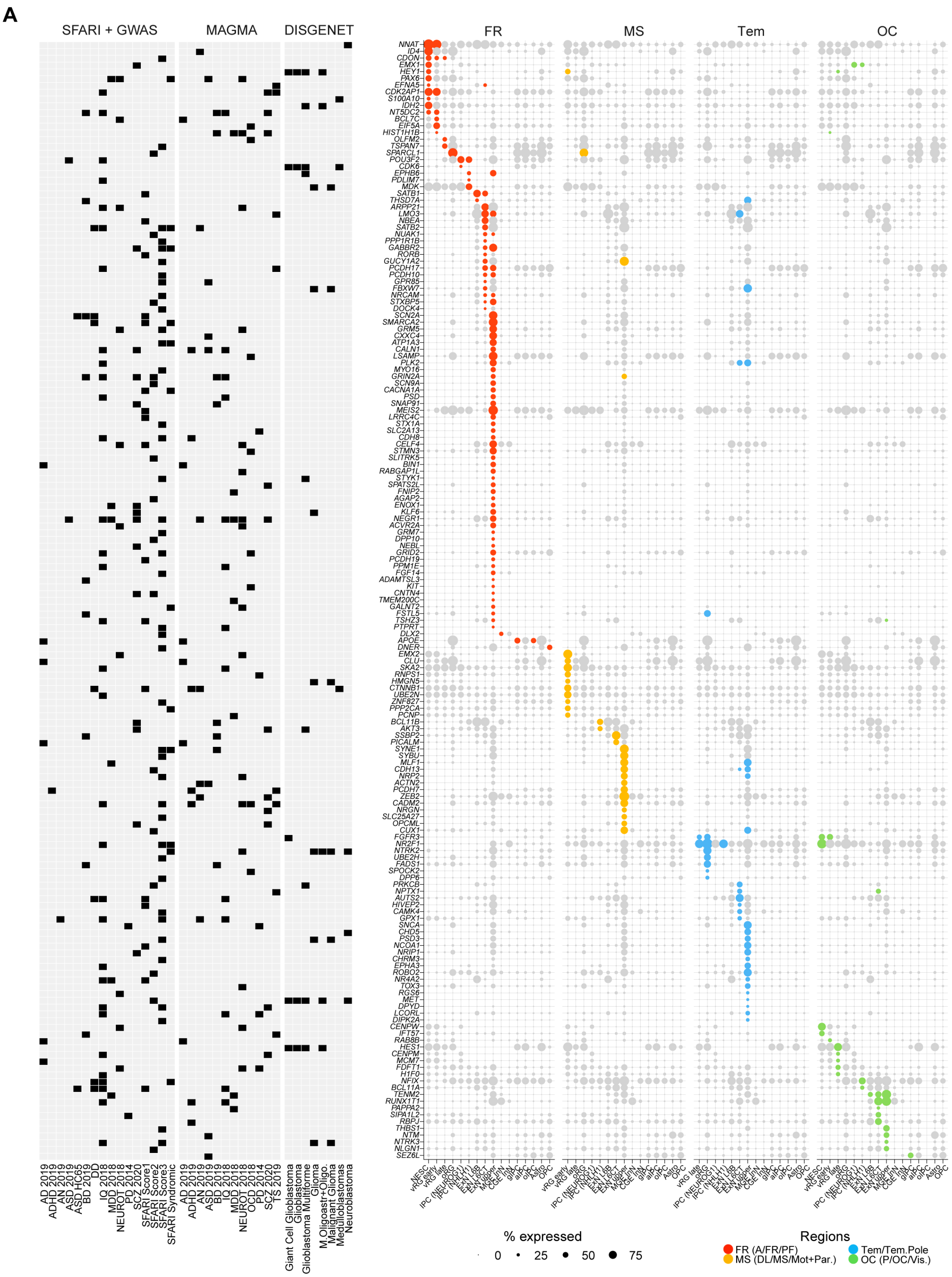
A
SFARI + GWAS
MAGMA
DISGENET
FR
MS
Tem
OC
% expressed
0 25 50 75
Regions
FR (A/FR/PF)
MS (DL/MS/Mot+Par.)
Tem/Tem.Pole
OC (P/OC/Vis.)

**Fig. S16. Regional expression preference of brain disease-risk genes across telencephalic development** (**A**) For each region, all the disease risk genes showing significant cell subtype enrichment and regional enrichment were selected and visualized. In the dot plot, lightgrey colors refer to no regional enrichment while region colors denote regional enrichment.

### Materials and Methods

All procedures involving animals, including monkeys and mice, were carried out according to guidelines described in the Guide for the Care and Use of Laboratory Animals, and were approved by the Yale University Institutional Animal Care and Use Committee (IACUC).

#### Caesarean sections of the pregnant monkeys and collection of the fetal brains

Rhesus macaque monkeys were bred in Rakic and Sestan primate breeding colony at Yale. Timed-pregnant monkeys were subjected to caesarian section at the required gestational age, performed by Yale's Veterinary Clinical Services (VCS). Monkeys were first sedated with ketamine (3 mg/kg) and atropine sulfate (Lily, 0.2 mg/kg). A butterfly catheter was introduced into the saphenous vein for continuous administration of fluids to prevent dehydration. Intravenous leads were secured subcutaneously for monitoring heart rate and respiration throughout the surgical procedure performed under isoflurane anesthesia and strict sterile procedures. A midline incision was made in the abdominal wall and the uterus gently exposed through the opening. The uterus was then incised between the primary and secondary lobes of the discoid placenta, the chorioallantoic membrane was punctured and the embryo delivered, decapitated still under the effects of anesthesia from maternal blood. The embryo head was transported to an adjacent room housing a BSL2 hood, brain tissue was dissected for single cell transcriptomics or fixed in paraformaldehyde (PFA). After delivery, the uterine and abdominal walls of the mother monkey was sutured in layers. Post-operatively, the animals were monitored several times a day until full recovery.

#### Fetal brain single cell dissociation

Fetal macaque brains were isolated from E37 to E110, put on a dish containing PBS and sectioned. Telencephalic regions were identified and dissected using a blade. For E37-43 brains, the entire anterior and medio-posterior parts of the hemispheres were cut. Each tissue was incubated with HBSS-Papain (2 mg/ml, BrainBits) from 15 (for the early ages) to 30 (for more advanced ages) minutes at 37$^{o}$C. The solution was removed and the tissue gently triturated in HBSS-DNase I 0.1mg/ml solution (StemCells) using a 2 ml pipette. Then, samples were filtered through 30 μm strain filters and the cells counted with an automatic cell counter (ThermoFisher Scientific). Samples were diluted to 1000 cells/microliter and processed for single cell RNA-seq analysis within 20 min at Yale Center for Genome Analysis (YCGA) core facility.

**Fixation and sectioning of the brain tissue**

Macaque and mouse fetal brains were dissected and immerse in 4% PFA overnight (ON) at 4$^{o}$C. Fixed brain blocks were immersed in step-gradients of sucrose/PBS up to 30% for 2-3 days at 4$^{o}$C, then embedded in OCT and frozen at -80$^{o}$C. Sagittal sections were prepared at 25 $\mu$m or 16 $\mu$m for the monkey and mouse brain tissue, respectively. Sections were prepared using a Leica CM3050S cryostat and stored at -80$^{o}$C until use.

**Construction of 10X Genomic Single Cell 3' RNA-Seq libraries and sequencing**

Sample Preparation. The first step of scRNA-Seq involved preparation of the single cell suspension.

GEM Generation and Barcoding. Single cell suspension in RT Master Mix is loaded on the Single Cell A Chip and partition with a pool of about 750,000 barcoded gel beads to form nanoliter-scale Gel Beads-In-Emulsions (GEMs). Each gel bead has primers containing (i) an Illumina® R1 sequence (read 1 sequencing primer), (ii) a 16 nt 10x Barcode, (iii) a 12 nt Unique Molecular Identifier (UMI), and (iv) a poly-dT primer sequence (30nt). Upon dissolution of the Gel Beads in a GEM, the primers are released and mixed with cell lysate and Master Mix. Incubation of the GEMs then produces barcoded, full-length cDNA from poly-adenylated mRNA.

Post GEM-RT Cleanup, cDNA Amplification and library construction. Silane magnetic beads are used to remove leftover biochemical reagents and primers from the post GEM reaction mixture. Full-length, barcoded cDNA is then amplified by PCR to generate sufficient mass for library construction. Enzymatic Fragmentation and Size Selection are used to optimize the cDNA amplicon size prior to library construction. R1 (read 1 primer sequence) are added to the molecules during GEM incubation. P5, P7, a sample index, and R2 (read 2 primer sequence) are added during library construction via End Repair, A-tailing, Adaptor Ligation, and PCR. The final libraries contain the P5 and P7 primers used in Illumina bridge amplification.

Sequencing libraries. The Single Cell 3' Protocol produces Illumina-ready sequencing libraries. A Single Cell 3' Library comprises standard Illumina paired-end constructs which begin and end with P5 and P7. The Single Cell 3' 16 bp 10x Barcode and 12 bp UMI are encoded in Read 1, while Read 2 is used to sequence the cDNA fragment (91bp). Sequencing a Single Cell 3' Library produces a standard Illumina BCL data output folder. The BCL data will include the paired-end Read 1 (containing the 16 bp 10x™ Barcode and 12 bp UMI) and Read 2 and the sample index in the i7 index read. Minimum sequencing depth is 20,000 read pairs per cell.

Single Cell 3' library analysis. The Cell Ranger™ analysis pipelines perform secondary analysis and visualization. In addition to performing standard analysis steps such as demultiplexing, alignment, and gene counting, Cell Ranger™ leverages the 10x Barcodes to generate expression data with single-cell resolution. This data type enables applications including cell clustering, cell type classification, and differential gene expression at a scale of hundreds to millions of cells.

**Single-cell RNA-seq data processing and filtering**

Cellranger was applied to align the scRNA-seq reads to rhesus macaque genome assembly Mmul10 together with the gene annotation file from NCBI RefSeq (release 103), followed by barcode counting and unique molecular identifier (UMI) quantification. The resulted filtered gene by cell UMI count matrices were used for additional quality control and filtering.

An initial clustering was performed for each sample using Seurat (*82*) to spot potential low-quality cell clusters, which include cells with low number of UMIs and/or high percentage of mitochondria UMIs. The resulted count matrices were used in scrublet package (*83*) to predict the doublet score of each cell. Cell clusters with high doublet scores and exhibiting combinatory expression of two different cell type markers were considered as doublet clusters and removed. Because samples might behave differently, a sample-wise doublet score threshold were selected for each sample. To further remove such outlier cells, cells belonging to the same cell class across different batches were clustered together for additional rounds of quality control, which increased the power in detection of outliers. The filtered gene by cell UMI count matrices were utilized in the downstream analysis.

**Normalization, clustering and dimension reduction of the scRNA-seq data**

Filtered UMI counts in each cell was first log-normalized using the *NormalizeData* function in Seurat (*82*), with the scaling factor set as 10,000. To embed all cells across different development ages and brain areas in the same reduced dimension space, we applied fastMNN (*84*) and Harmony (*85*) to integrate the data. Here, both methods perform batch correction in the reduced dimensions and they overall exhibited very similar results. However, Harmony showed slightly better performance in preserving inter-cell heterogeneity and fastMNN were marginally better in recapitulation of cell differentiation lineages. And accordingly, Harmony was largely utilized for cell cluster identification and fastMNN for lineage visualization (e.g., Fig. 1). Prior to batch correction, for each batch we identified the highly variable genes using the variance-stabilizing transformation implemented in the Seurat package (*86*). Because direct the default selction of the highly variable genes with the highest frequencies across all samples might lose some key signals in certain developmental ages with smaller sample sizes, we parcellated the developmental stages to three windows based on their transcriptomic similarity (E37-E43, E54-E78, E93-E110) and integrated the highly variable genes using the *SelectIntegrationFeatures* function in Seurat. The union of the genes across the three developmental periods were chosen for downstream analysis. Then, the normalized data were scaled separately in each batch (for Harmony), or together across all batches (for fastMNN), and used for principal components analysis (PCA), with elbow plots to select the significant principal components. The first 30 integrated reduced dimensions were used for UMAP visualization with the “umap-learn” method and “correlation” metric (*87*).

Cell clustering was performed via “Louvain” algorithm based on the first 30 integrated reduced dimensions with the k-nearest neighbor set to 25. Dividing cells from different cell types (e.g., early and late RGCs) might cluster together by cell cycle phases rather than their identities. Therefore, we categorized cell clusters into different cycling phases based on the expression of key cycling genes (e.g., S phase – *PCNA* and *MCM5*; G2M phase – *MKI67* and *TOP2A*) and gene set enrichment of cell cycling genes calculated via the Seruat *CellCycleScoring* function, and further sub-clustered cells within each phase to maximize the variance contribution from cell type heterogeneity rather than cell cycling differences. This method showed better performance than the traditional method that clusters cells using the data with cell cycle scores regressed.

**Cell subtype annotation**

Patterning center subtypes were identified based on several criteria: 1) temporal enrichment at E37 and E42-43; 2) high expression of neuroepithelial markers such as *SOX2* and *NES*, low expression IPC makers (e.g., *ASCL1*, *NEUROG1* and *EOMES*); 3) combinatory expression of known patterning center markers: rostral patterning center (FGF8$^+$, *FGF17*$^+$) (*26, 28*), cortical hem (*RSPO3*$^+$, *TTR*$^+$) (*30*) (*31*), antihem (*SFRP2*$^+$, *PAX6*$^+$) (*33*) and zona limitans (*IRX*, *WNT3*$^+$, *WNT4*$^+$) (*32*); 4) the spatial locations are consistent with the predicted identities. Furthermore, we spotted several early subtypes enriched in E37-43 and co-clustered with patterning center subtypes, which likely represent regional specialized domains. These included two NKX2-1+ subtypes (*LMO1*$^+$ and *NKX6-2*$^+$, respectively) detected in anterior samples representing anterior-ventral (AntVen) domain cells. Both subtypes were transcriptomically similar to the two *NKX2-1*$^+$ early GE subtypes (GE RG *NKX2−1 OLIG1* and GE RG *NKX2−1 OLIG1*) and all the four subtypes co-clustered with the rostral patterning center (*FGF17*$^+$; Fig. 2). We also identified subtypes forming continuous manifold and connecting with the patterning center subtypes and those domain-specific subtypes, resembling cell differentiation lineages. These contained the anterior-ventral domain *NKX2-1*$^+$/*LMO1*$^+$ subtypes give rise to the *DLX1*$^+$ IPC subtype (inIPC *ASCL1 DLX1*) which produces *GNRH1*$^+$ interneurons and *LHX8*$^+$/*ZIC1*$^+$ interneurons. The rostral patterning subtypes seemed to form lineages with IPC *FGF17* subtype and neuron subtype Neu *TAGLN3 ONECUT2*. The zona limitans subtype also led to a lineage with IPC *TCF7L2* subtype. In addition, one *NKX2-1*$^+$ subtype (*RAX*$^+$) in posterior domain (Pos), and one *SFRP1*$^+$ subtype with limited markers present in anterior domain (Ant) were also dectected. A few other early subtypes, including Cls *FGF17 LGI1*, Cls *LHX9 EBF1*, Cls *RSPO3 SOX1*, Cls *GSX2 B3GAT2*, were all small in sizes and their identities were left as unknown.

Neural stem cells in dorsal regions clustered together and were defined based on the expression profiles of *PAX6*$^+$/*SOX2*$^+$/*NES*$^+$/*EOMES*$^-$/*EGFR*$^-$. These cells formed a circular shape on the UMAP layout representing cell cycling states and they were also organized along the temporal axis resembling the progression of their identities. This started with putative neuroepithelial stems cells, which were identified based on the high expression of *RSPO3* and relatively lower expression of *PAX6*. Two early

RGC subtypes were defined at E37-43, one expressing *FABP7* and *PMP22*, enriched at anterior regions, and the other one labeled by *HMGA2* and *CCND1* ubiquitously present in all regions analyzed. Late RGCs included two oRG (*HOPX*$^{+}$/*NRG1*$^{+}$) subtypes, two tRG (*CRYAB*$^{+}$) subtypes and one vRGC subtype directly connecting with early RGCs and showing low expression of oRG and tRG markers. There is also an ependymal subtype marked by *FOXJ1* expression as well as well several CFAP genes (e.g., *CFAP45* and *CFAP54*). Neural stem cells in ventral regions also express *SOX2* and *NES*, but only the late RGCs express *PAX6*. We also categorized them into different subtypes based on the similar genes we used for dorsal neural stem cells including *HMGA2*, *HOPX* and *CRYAB* as well as ventral-specific signatures such as *NKX2-1*, *NKX6-2* and *OLIG1*.

Cajal Retzius cell lineages were extracted based on their evident *RELN* expression. Anterior-enriched Cajal Retzius cells showed higher expression of *ETV1*$^{+}$ while the posterior-enriched ones had high *TP73* expression (*88, 89*). We also found two putative *RSPO3*$^{+}$ IPC subtypes enriched in the posterior regions and they are marked by *NEUROG1* and *NHLH1* expression respectively.

Excitatory neuron lineages included IPCs marked by *EOMES* expression and post-mitotic neurons expressing *NEUROD2*. The IPCs consisted of three subtypes: *VIM*$^{+}$ subtype transcriptomically more similar to radial glial cells, *NEUROG1*$^{+}$ subtype dominating the cycling IPCs which also has non-cycling cells, and *NHLH1*$^{+}$ postmitotic subtype coming from *NEUROG1*$^{+}$ subtype. Subtypes of excitatory neurons were broadly categorized into deep and upper layer neurons based on their expression of *SOX5* and *CUX2*, respectively (*90, 91*). Deep layer neurons included two nascent subtypes (*PALMD*$^{+}$ and *ID2*$^{+}$), a corticothalamic subtype (*SYT6*$^{+}$), two intratelencephalic subtypes (*OPRK1*$^{+}$/*SULF1*$^{+}$, *OPRK1*$^{+}$/*NR4A2*$^{+}$) and a L6B subtype (*NR4A2*$^{+}$/*GRID2*$^{+}$) (*17, 48, 92*). Upper layer neurons contained two nascent subtypes (*PALMD*$^{+}$ and ADRA2A$^{+}$) and one intratelencephalic type (*ACTN2*$^{+}$) (*17, 48, 92*). We identified also a putative deep layer subtype (*SOX5*$^{+}$/*KCNV1*$^{+}$) which is transiently present at E62-64 and one excitatory neuron subtype enriched at posterior cingulate cortex (*TSHZ2*$^{+}$/*NR4A3*$^{+}$).

Interneurons were classified based on *DLX1*, *DLX2, GAD1 and GAD2* expression. IPCs in the interneuron lineages were categorized based on their *ASCL1* expression and their co-clustering on the UMAP connecting with post-mitotic interneurons. Putative identities of the postmitotic interneurons were classified by the expression of markers highly correlated with their developmental origins (MGE: *LHX6*;

C/LGE: *NR2F2*, *SP8*, *MEIS2*) and transcriptomic integration with existing datasets with established identities including an independent developing macaque interneuron scRNA-seq data (*52*) and an adult macaque snRNA-seq data (*48*). The *LHX6*$^+$ interneurons consisted of three major branches: *CRABP1*$^+$ interneurons recently reported to be enriched in primates (*16, 52*), *LHX8*$^+$ branch enriched at E37-43 likely contributing to cholinergic neurons (*54*), and *LHX6*$^+$/*CRABP1*$^-$ interneurons dominating the MGE-derived interneurons and giving rise to majority of the *SST* and *PVALB* interneurons. Within the *LHX6*$^+$/*CRABP1*$^-$ group, we spotted three potential sub-branches: *SST*$^+$/*NPY*$^+$ branch giving rise to long projecting inhibitory neurons (*48, 53*), *GUCY1A2*$^+$/*RELN*$^+$/*DCN*$^+$ branch producing major *PVALB* and *SST* interneurons; *CCK*+ branch generating *LAMP5 LHX6* interneurons that mapped to mouse hippocampus Ivy cells and recently reported to show abundance enrichment in primate neocortex (*16, 17, 93*). The CGE and LGE-derived interneurons encompassed two major branches marked by *NR2F2*/*SP8* and *MEIS2*/*SP8* expression, respectively. Within the *MEIS2*/*SP8* lineage, cells were largely parcellated by the expression of *PAX6* (olfactory bulb neurons) and *FOXP1*/*FOXP2* (striatum spiny projection neurons) (*52*). The *NR2F2*/*SP8* lineage consist of cell subtypes giving rise to different adult interneuron subclasses (*48*): *LAMP5*$^+$ interneurons becoming *LAMP5 RELN* subclass, *KIT*$^+$ interneurons becoming *ADARB2 KCNG1* subclass; *VIP*$^+$ interneurons becoming *VIP* subclass.

The glia cells were classified into three major groups based on expression of *OLIG2* (oligodendrocyte lineage-related cells), *AQP4* (astrocyte lineage-related cells), and *EGFR* (glia precursor cells). The oligodendrocyte lineage-related cells were further divided by the expression of *PDGFRA* (oligodendrocyte precursor cells, OPCs), *PDGFRA* and *MKI67* (oligodendrocyte precursor cells in proliferation stage, OPC *PDGFRA MKI67*), *PCDH15* (late oligodendrocyte precursor cells), and *MBP* (oligodendrocytes). The astrocyte lineage-related cells were divided into three subtypes by the expression of *GFAP*, *EGFR*, and *MFGE8*, respectively. Regarding glia precursor cells, we employed combinational expression of genes to sort them into astrocyte intermediate precursor cells (aIPCs, *EGFR*$^+$/*AQP4*$^+$/*IGFBP2*$^+$), oligodendrocyte intermediate precursor cells (oIPCs, *EGFR*$^+$/*PDGFRA*$^+$/*DLL1*$^+$), glia intermediate precursor cells (gIPCs, *EGFR*$^+$/*AQP4*$^-$/*PDGFRA*$^-$), glia intermediate precursor cells in proliferation stage (*EGFR*$^+$/*MKI67*$^+$).

The rest of the immune cells and vascular-related cells were categorized using the following strategies. All immune cells were identified as *PTPRC*$^+$, with microglia subtype further identified as *C1QC*$^+$ and T cells identified as *CD69*$^+$ (*48*). Two red blood lineage cell subtypes were classified based on the expression of *HBA1* and their unique expression of *HBE1* and *SNCA*, respectively. Vascular cells were characterized by their *FN1* expression and specific expression of other subtype markers: endothelial cells (*CLDN5*$^+$), pericytes (*GRM8*$^+$), smooth muscle cells (*ACTA2*$^+$) and vascular lepotomeningeal cell (*CEMIP*$^+$) (*48*). We also spotted two putative mesenchymal cell subtypes: one expressing *LUM*, consistent with the recently reported human mesenchymal cells in the human early developing telencephalon (*94*); the other is marked by *FOXD3* and *PLP1* expression, likely representing neural crest cells (*95, 96*).

**Identification of cell subtype markers and genes with region-divergent expression**

To identify genes differentially expressed between cell subtypes or brain areas, Wilcoxon Rank Sum test was utilized, and a minimum expression ratio of 0.1 and a Bonferroni-adjusted p value threshold of 0.01 were adopted. In certain analyses as detailed in the following sections, we incorporated additional requirements to filter the differentially expressed genes. For example, we inspected expression ratio fold changes, with a pseudo value of 0.01 adding to the gene expression ratio in the examined cell subtype (numerator) and background cells (denominator). Other criteria such as log fold changes of average expression and background expression ratios were also taken into consideration for gene filtering.

**Inference of regulatory networks in telencephalic organizer cells and regional specific NSCs**

The inference of transcription factor regulatory networks was suggested by the SCENIC workflow (*97*), with motif enrichment and gene co-expression information integrated. In this analysis, for simplicity, we merged the two anterior-ventral *NKX2-1*$^+$ subtypes (AV *NKX2-1 LMO1* and AV *NK2-1 NKX6-2*), as well as the two GE *NKX2-1*$^+$ subtypes (GE RG *NKX2-1 DLK1* and GE RG *NKX2-1 OLIG1*). The putative promoter region (upstream 800 bases and downstream 100 bases of the transcription start site) of each gene, as well as the motifs of transcription factors, were prepared (*98*). R package PWMEnrich (*99*) was used to perform the motif enrichment analysis, with p value threshold set at 0.05 to only retain transcription factor genes with significant motif enrichment at the promoter of the given gene. This analysis generated raw regulons, which refers to a module of genes including a transcription factor gene and a list of putative targets. Then, within each relevant cell subtype, we calculated the subtype markers using the *FindMarkers* function in Seurat (*82*) and intersected the markers with the raw regulons to filter transcription factors and targets. To gain a more robust expression association between the transcription factor and targets, we correlated their expression across pseudobulk samples of the subtype. Specifically, we ordered the cells along the UMAP-1 axis calculated via the Seurat *RunUMAP* function with a setting of dimension equal to one, and parcellated cells into 30 equal-width bins along the axis and removed bins with less than 15 cells. Average expression was calculated in each bin, which was used to assess the expression correlation between transcription factors and the predicted targets. To further evaluate the robustness of the expression correlation, we also permutated the gene expression in each subtype, maintaining the average expression levels and variance of each gene but disrupting the gene-gene correlations. The permutation was repeated for 1000 times and the p value for the correlation of a given gene pair is defined as (n + 1) / (1000 + 1), where n represents the number of permuted correlation coefficients exceeding the actual correlation coefficient. Only transcription factor-target pairs with significant correlations (p value < 0.1 and coefficient > 0.1) were retained. The regulons in each cell subtype were then merged and visualized in a network with arrows indicating regulatory directions from transcription factor to targets and nodes colored by the cell subtypes showing the significant regulation.

**Transcriptomic comparison between mouse and macaque organizer domain subtypes**

To assess the transcriptomic similarity between macaque and mouse telencephalic organizer domains (*22*), we derived the subtype markers in each dataset using the *FindMarkers* function in Seurat and extracted the shared subtype markers between the two species. These included many key genes labeling homologous cell subtypes. Average expression of the shared subtype markers were calculated across subtypes followed by Pearson correlation coefficient measurement for each pair of subtypes between the two datasets. To avoid noise from background transcriptomic similarities, any correlation coefficients below the 80% quantile of all values were removed. The filtered subtype similarity was visualized in a Sankey plot, which illustrate subtype matching between the two species.

Alternatively, we calculated the enrichment of mouse subtype markers in this dataset through the AUCell algorithm (*97*). Because this method uses rank-based expression values to assess expression enrichment, it is robust to potential quality differences between the mouse and macaque datasets. For a given set mouse subtype markers, we averaged its enrichment scores in each macaque subtype and visualized the results in a heat map, which recapitulated the subtype similarity patterns shown by the above correlation-based analysis.

In order to find species-specific expression patterns in homologous subtypes and avoid potential batch effects, we set a high threshold for the differential expression test. To identify macaque-enriched genes, we first extracted the top 100 markers for each macaque subtype and removed the genes that were also markers of the homologous mouse subtypes. Next, a minimum expression ratio of 0.2 was required in macaque subtypes whereas a maximum expression ratio of 0.05 was set in the mouse homologous subtypes. For each macaque subtype, the top 10 genes ranked by their expression fold changes between the given subtype and the background macaque cells were selected and visualized. The same approach was used to find mouse-enriched genes in homologous subtypes.

**Ligand-receptor mediated cell-cell communication between pattern centers and early neural stem cells**

We applied two complementary expression-based approaches, CellChat and CellphoneDB (*100, 101*), to infer putative cell-cell communications between organizer domain cell subtypes and early neural stem cells. In these analyses, we only included subtypes potentially secreting patterning ligands (RPC, anterior ventral domain subtypes, and cortical hem) and neural stem cell subtypes responding to these molecules (i.e., NESC and early vRG cells from anterior, posterior regions and ganglionic eminence). Each subtype was down-sampled to have equal number of cells (1000 cells), otherwise subtype size differences could affect marker detection in CellChat and permutation in CellphoneDB. In both analyses, we set a minimum expression ratio of 0.05 and p value threshold at 0.05. For the resulted ligand-receptor interactions, we only considered the directions with ligands expressed in organizer domain subtypes and receptors in regional neural stem cells. Because the output results from the two analyses are largely shared, we set the CellChat-based results as the reference and incorporated additional interactions reported by CellphoneDB-based analysis.

To get a broad view of the interaction patterns between ligand-receptor pairs, we performed t-SNE analysis using the interaction matrix, with rows as ligand-receptor pair names and columns as cell subtype pairs. In addition, we clustered the ligand-receptor pairs based on their orchestrated cell-cell interaction patterns using robust sparse K-Means clustering algorithm (*102*). The resulted 10 clusters are well separated on the t-SNE layout and further confirmed the distinct cell-cell interaction patterns mediated by ligand receptor pairs.

**Lineage construction from organizer domain progenitors to offspring cells**

To define the lineage progression from organizer domain progenitors to their progeny cells and also delineate the gene cascades along the lineage, RNA velocity analysis using scVelo package (*103*) was applied. In each lineage, UMAP layout was first obtained via the *RunUMAP* function in Seurat and the resulted Seurat object was converted to anndata for scVelo anslysis. For simplicity, cycling cells were not included in the analysis. After data filtering, normalization, identification of highly variable genes and computing moments for velocity estimation, dynamic model was applied to compute the RNA velocity vectors and pseudotime. The top 300 genes showing transcriptional variations along the lineages were visualized on heat maps.

**Transcriptomic comparisons with published data**

We applied the following two methods to evaluate cell subtype matching between datasets. In the first method, cross-dataset cell subtype similarity was measured by Pearson correlation coefficients. Specifically, the intersection of the highly variable genes between this study and a given published dataset were selected, and the average expression of these genes across each cell subtypes were computed via the *AverageExpression* function in Seurat (*82*) followed by log-transformation. Pearson correlation coefficients were calculated on the log-transformed average expression for each pair of subtypes between this study and the given published dataset. In the other approach, cells from this study and a given published dataset were integrated and visualized on UMAP to evaluate the subtype alignment. Because there were prominent batch effects between this study and publish datasets, largely attributed to differences of species, developmental stages and technical approaches, we applied Seurat integration algorithm (*82*), as a stringent method to remove batch effects. Here, the intersection of the top 2,000 highly variable genes from each study were used for canonical correlation analysis, followed by anchor finding ad hierarchical integration of normalized data using the *IntegrateData* function. The integrated data were then scaled, used for principal components analysis and UMAP visualization.

**Region-specific gene expression cascades**

We used the following approaches to construct the region-specific expression cascades. For each subtype in a given region, we calculated the genes showing expression enrichment in this region compared to all other regions using Wilcoxon Rank Sum test. To avoid the influence of cell number differences, for each subtype we downsampled each region to have the same number of cells. The differential expression analysis results were further filtered based on expression ratios, fold changes of average expression, fold changes of expression ratio and Bonferroni-adjusted p values, to get genes with most salient regional enrichment. By leveraging the defined pseudotime that organize cells from different regions on the same scale, we parcellated cells from the four regions into bins with equal pseudotime width. We then calculated the average gene expression along the pseudotime bins for each region and ordered regionally enriched genes based on their sequential waves of expression. The gene ordering was obtained by first fitting into impulse models (linear, single sigmoid or double sigmoid) implemented in the URD package (*104*), which returned the positions representing where gene expression arises and diminishes. To gain a better visualization of the expression patterns, average expression along the pseudotime bins were smoothed using *loess* function in R.

**Gene Ontology enrichment analysis**

Gene ontology (GO) enrichment analysis was performed by the Bioconductor package 'topGO' (http://bioconductor.org/packages/release/bioc/html/topGO.html) using the Fisher's exact test followed by FDR adjusting p values. Only GO terms under biological processes were included in the analyses and a threshold of FDR < 0.1 was selected to pick the significant terms.

**Transcriptomic divergence between the shared subtypes from different regions**

In order to evaluate the magnitude of regional differences between different cell types, which might come from the same lineage (e.g., IPCs, nascent and mature excitatory neurons) or belong to the same cell class (e.g., *LHX6*$^+$ versus *NR2F2*$^+$/*SP8*$^+$ interneurons), we applied the Augur algorithms to measure the transcriptomic separability of regions for each relevant subtype (*105*). To recapitulate regional variations as much as possible, for each batch we extracted the highly variable genes across the cells from all the analyzed brain areas and used the *SelectIntegrationFeatures* in Seurat to identify the top regionally variable features. In running Augur algorithms, these highly variable genes were directly used, with the mode set to “velocity” to avoid additional detection of highly variable genes. The Augur analysis was performed for each pair of regions, which provided a detailed view of transcriptomic divergence of cell types across brain areas.

In addition, we used the number of differentially expressed genes to evaluate regional difference changes along RGC progression and excitatory neuron differentiation and maturation (Figs. 4C and 5C). We parcellated cells into different equal-width bins along the pseudotime and downsample the cells from each region to have a balanced number of cells across bins and regions. Then we applied Wilcoxon Rank Sum test and calculated the number of differentially expressed genes between regions along the pseudotime bins followed by visualizing in log scale and smoothed via loess function.

**Hierarchical clustering of excitatory neuron lineage subtypes across cortical regions**

To check whether the regional differences of the excitatory neurons are correlated with the anatomical proximity of the brain regions they populate, we leveraged the refined regions sampled at E93 and E110. We first calculated the highly variables genes at each batch (here is individual) and used the Seurat *SelectIntegrationFeatures* function to capture the top 2000 genes with highest expression variability across analyzed brain regions. For a given excitatory neuron subtype, average expression was calculated in each region and cells from different regions were down-sampled to have the consistent number of cells (100 cells) prior to average expression calculation. The resulted average expression of the 2000 highly variable genes from the shared subtype in different regions were used for hierarchical clustering. Here, the distance matrices were defined as Pearson correlation coefficients subtracted from 1 and were subsequently used by the *hclust* function with “ward.D2” algorithm for clustering. Dendrogram visualization was achieved via the *circlize* R package (*106*).

To obtain robust estimate of region co-clustering, we generate 1000 bootstrap replicates from the above 2000 highly variable genes, resulting 80% of genes for each replicate. For each replicate, the same subtype across regions were clustered using the above strategy followed by cluster separation via splitting the hierarchical tree to k=3 clusters. The frequencies of region co-clustering were measured and visualized by heat maps. In addition, we permutated the gene expression for 1000 replicates, keeping the gene-wise characteristics (e.g., mean expression, variance) but destroying the gene-gene relationship. The average expression from the 1000 replicates of permuted data were used following above bootstrap clustering strategy to see how often subtypes from different regions cluster together. By comparing the co-clustering frequency in the actual data versus the permuted data, we confirmed the region co-clustering in IPCs reflect true regional differences

**Correlation of regional gene expression specificity between cell types**

To assess the effect of region-specific environment on cell type identities, we correlated the region-specificity of gene expression across multiple cell types. Specifically, we identified all the genes divergently expressed across brain areas and assessed their expression enrichment in each region. The region enrichment scores were derived through dividing the average expression in the given region by the average expression in other background regions, with a pseudo-value of 0.1 added to both the numerators and denominators. Within each region, the pairwise comparisons of such enrichment scores were visualized in dot plots and Pearson correlation coefficients were also calculated.

**Hierarchical clustering of regional genes expression patterns for inhibitory neurons**

Although Augur algorithm detected lower regional transcriptomic differences in inhibitory neurons, we could still find certain genes differentially expressed across cortical regions in the shared inhibitory neuron types. In order to obtain an overview of their global expression patterns across regions, we applied a hierarchical strategy. Here, we only considered two major cortical inhibitory neuron groups: *LHX6*$^{+}$/*CRABP1*$^{-}$ cells and *NR2F2*$^{+}$/*SP8*$^{+}$ cells. Regionally divergent genes were obtained through Wilcoxon Rank Sum test with the Bonferroni-corrected P value threshold set at 0.01. In case cell numbers affect the number of differentially expressed genes, we downsampled the cells in each region to the same level. For each region, we generated 20 pseudobulk samples each containing 200 random cells and calculated the average expression of the regionally-divergent genes across all pseudobulk replicates. The resulted expression matrix was used for hierarchical clustering, with the distance matrix defined as one minus Pearson correlation coefficients between gene pairs and clustering algorithm set as “ward.D2”.

**Lineage inference of the switch between neurogenesis and gliogenesis**

To explore the developmental transition from neurogenesis to gliogenesis, we employed two different approaches: unsupervised transcriptomic clustering by Seurat (*82*) analysis pipeline and cell lineage tracing by Monocle (version 2) analysis pipeline (*107*), to analyze the transcriptional association in-between radial glial cells, excitatory neurons and glia cells and define their lineage relationship.

In the Monocle analysis pipeline, we firstly recruited Seurat *FindMarkers* function to perform differential expression analysis for any pair of cell subtypes in each developmental stage. Subsequently, the differential expressed genes were accumulated and used to infer cell trajectory. Following up the recommended workflow, the suggested default parameters in Monocle were preferred, except that the 'DDRTree' method was use as reduction model and the batch correction was introduced. To compute the pseudotime for each cell along the cell trajectory tree, the tree root was manually selected through analyzing the tree structure and the distribution of cellular nature ages. Lastly, we used Seurat *FindMarkers* function to perform differential expression analysis for any pair of branches to identify genes specific to each tree branch, which led to a proxy for detecting the transcriptional program that dominates the segmentation and emergency of each tree branch.

**Global cross-dataset comparison for glia cells**

The glial cell types in the current study were compared to the external datasets, including datasets from prenatal humans, adult monkeys, and lifespan mice, to confirm the quality of cells and the precision of cell type annotation. For astrocytes, we defined three subtypes, which together with astrocyte precursor and glia precursor were compared among multiple brain regions to reveal their regional distribution. One external astrocyte dataset collected from different cortical layers in P14 mice (*59*)  was compared with our astrocytes to show their laminar distribution, and another external astrocyte dataset collected from dorsolateral prefrontal cortex in adult macaque (*48*) were compared with our astrocytes to investigate the correspondence between immature and mature stages. We used specScore script to compute specificity score for each gene in each subtype and correlated subtypes across datasets using the specificity scores, with the pairwise subtype similarity visualized by alluvial plots. On the other hand, the cross-dataset comparison was conducted by using abovementioned UMAP pipeline. Firstly, the different datasets were separately wrapped according to Seurat analysis workflow, which includes abovementioned a range of processes. Secondly, fastMNN (*84*) was used to correct unwanted batch variation by choosing the different datasets as the major source of systematic variation. Alternatively, other integration methods including Harmony (*85*) and LIGER (*108*) were considered to verify the results reported by fastMNN, whose analyses were not shown when an negligible difference of cell type correspondence was observed.

**Predication of ligand-receptor mediated cell-cell communication in mid-fetal stage**

In the monkey mid-fetal stage, E93 and E110 in current study, most of neuron and glia cell types have emerged, in particular for astrocytes that gradually increase and interplay with other cells. The communication in-between cells can be inferred by correlating the expression of ligand and the corresponding receptor genes. Rather than calculating cell-cell communication, we utilized CellChat (*101*) to compute the averaged communication in-between major cell types, including excitatory neurons, interneurons, astrocyte, enIPCs, OPC/oligodendrocyte, microglia and endothelial cells. The cell subtypes were aggregated into major cell types to achieve high accuracy with the reduction of single cell noise. Cell-cell communication were computed in each brain region separately, and subsequently the interaction strength between any pair of cell types were averaged among brain regions to provide a proxy of the general tendency of cell-cell communication.

**Compilation of brain disease risk gene lists**

We compiled disease-risk genes from DISGENET (*109*) and filtered-out those diseases having fewer than 30 risk genes associated. Genes from “Mixed oligoastrocytoma” and “oligodendroglioma” were combined in the “Mixed Oligoastrocytoma+Oligodendrogliomas” list (abbreviated as M. Oligoastr.+ Oligod). Also, genes associated with any medulloblastoma were combined in the “Medulloblastomas” list.

Multiple genome wide associated studies (GWAS) were used to collect gene lists for the study: Alzheimer’s disease (AD) (*110*), anorexia nervosa (AN) (*111*), autistic spectrum disease (ASD) (*112*), bipolar disorder (BD) (*113*), intelligence quotient (IQ) (*114*), major depressive disorder (MDD) (*115*), neuroticism (NEUROT) (*116*), Parkinson’s disease (PD) (*117*) and schizophrenia (SCZ) (*118*).

Genes identified by running Multi-marker Analysis of GenoMic Annotation (MAGMA) (*119*) were also included for the following conditions: attention-deficit/hyperactivity disorder (ADHD), AD, AN, ASD, BD, IQ, MDD, NEUROT, obsessive-compulsive disorder (OCD) (*120*), PD, SCZ and Tourette syndrome (TS) (*121*). Only genes with a nominal p-value of less than 0.05 were used and we selected the top 200 genes according to their p-value.

Genes implicated in ASD susceptibility were also obtained from the SFARI database (https://gene.sfari.org). Only genes of SFARI categories S (syndromic), 1 (high confidence), 2 (strong candidate) and 3 (suggestive evidence) were employed.

Additionally, high-confidence ASD genes from (*122*), and a set of genes involved in developmental delay from the Deciphering Developmental Disorders Study consortium (*123*).

**Cell type enrichment of disease gene expression**

scRNA-Seq data were preprocessed following the guidelines for EWCE analysis (*124*). In brief, normalization and variance stabilization of scRNA-seq data using regularized negative binomial regression was performed with the sctransform R library (*86*). SCT-corrected counts were computed for genes being marker genes of any subtype with an adjusted p-value < 0.05 and having a 1-to-1 matching human ortholog. Macaque-to-human orthologs were retrieved using the ortogene R library (https://github.com/neurogenomics/orthogene).

Then, we performed EWCE's bootstrap enrichment test of the disease-associated gene lists previously defined (20000 repetitions, geneSizeControl=FALSE, controlledCT=NULL and mtc_method='BH'). Only the risk genes present in the dataset after SCT normalization were tested.

**Identification of disease risk genes showing as organizer domain subtype markers**

Marker genes were computed using the Wilcoxon Rank Sum test on log2-normalized data by *FindAllMarkers* from the Seurat package (*82*). We computed marker genes of all the cell subtypes, all the cell types and the organizer domain subtypes independently.

In those subsets, expression data were scaled. Averages of gene scaled expression were obtained for every gene in the disease-associated gene lists and we computed the median of averages per list. We also retained the number of genes from each list that were expressed in at least 10% of cells for each cell subtype.

Regarding organizer domain subtype's marker genes, we were interested in those that were expressed exclusively in any of the subtypes. For that, we selected those marker genes expressed in at least 10% of cells in the respective subtype and in less than 10% of cells of other organizer domain subtypes. With those markers, we created 2x2 contingency tables for each disease-risk gene list and subtype with counts of genes being or not subtype markers, and genes being or not disease-associated. Using those tables, we estimated the log2 odd ratio of being disease-associated and subtype marker using the R function fisher.test.

**Expression enrichment of disease risk genes across radial glial cells**

To gain a detailed view of the expression enrichment patterns of disease risk genes, we applied AUCell (*97*) enrichment analysis on the disease gene list across neural stem cell subtypes. To have robust estimation of each subtype and further avoid the influence of certain low-quality cells, we generated 100 pseudobulk samples in each neural stem cells by randomly pooling 200 cells, followed by calculating average gene expression in each of the pseudobulk samples. AUCell analysis was directly performed on these pseudobulk samples together with the disease gene list, with the results visualized on a heat map. Diseases were ordered according to their peak enrichment along the x axis (i.e., subtypes). Because we see many disease risk genes show high enrichment in outer radial glia cells, we wondered if it was caused by quality bias. Although AUCell package is robust to data quality differences, as it utilizes rank-based method to assess the separability of query genes versus background gene, we further subset the data to have 2000 UMIs per cell (cells with less than 2000 UMIs were not included) and generated pseudobulks and performed enrichment following the same strategy as described above. This overall lead to a similar enrichment pattern.

We next decided to identify the genes underlying the dynamics disease enrichment patterns. For each disease, we extracted its risk genes and correlated (Pearson correlation) their expression patterns with the given enrichment pattern of the disease. We set a threshold of 0.7 to capture the strongest signals and resulted genes were visualized in a heat map.

**Intersection of disease risk genes and region-specific signatures**

We set relatively high thresholds to identify disease genes showing prominent regional and cell-type enrichment. For each region, we calculated cell type markers and regionally-enriched genes using Wilcoxon Rank Sum test followed by setting the same thresholds for both analyses: Bonferrroni-corrected p value < 0.01, log fold changes of average expression > 0.25, expression ratio fold changes > 1.25, and expression ratio > 0.2. The intersection of the cell subtype markers and regionally-enriched genes were further overlapped with disease risk genes and the results were visualized in fig. S15. The intersection of the top 25 cell subtype markers and top 25 regionally-enriched genes ranked by fold changes of expression ratios were overlapped with disease risk genes and visualized in Fig. 7E.

**Single-molecule RNA in situ hybridization**

RNA in situ hybridizations were performed by Advanced Cell Diagnostics, Newark, CA, using the RNAscopeTM technology previously described in (*125*). Paired double-Z oligonucleotide probes were designed against target RNA using custom software. The probes used for rhesus macaque and mouse brain tissue samples are shown in table S7. RNAscope LS Fluorescent Multiplex Kit (Advanced Cell Diagnostics, Newark, CA, 322800) was used with custom pretreatment conditions following the instruction manual. Fixed frozen monkey and mouse fetal brain tissue slides were manually post-fixed in 10% neutral buffered formalin (NBF) at room temperature for 90 minutes. Then the slides were dehydrated in a series of ethanols and loaded onto the Leica Bond RX automated stainer, performing the reagent changes, starting with the pretreatments (protease), followed by the probe incubation, amplification steps, fluorophores, and DAPI counterstain. RNAscope 2.5 LS Protease III was used for 15 minutes at 40°C. Pretreatment conditions were optimized for each sample and quality control for RNA integrity was completed using probes specific to the housekeeping genes *Polr2a*, *Ppib*, and *Ubc*, which are low, moderate, and high expressing genes, respectively. Negative control background staining was evaluated using a probe specific to the bacterial *dapB* gene. Coverslipping was done manually using ProLong Gold mounting media at the end of each run.

**Human and macaque telencephalic organoid culture**

The human iPSC line Y6 was provided by Yale Stem Cell Center. This cell line was generated from neonatal skin fibroblasts using CytoTune™-iPS Reprogramming Kit (Invitrogen, A13780). Chromosome analysis was performed on cultured cells. Of the five metaphases examined, no structural and numerical abnormality was noted and the karyotype was consistent with that of a female (46, XX) complement. The pluripotency of the cells was confirmed by teratoma assay. Macaque iPSC line was generated by reprogramming E40 macaque lung fibroblasts using CytoTune-iPS 2.0 Sendai Reprogramming kit (ThermoFisher Scientific, A16517) and authenticated by morphology and karyotyping. All human and macaque iPSC lines were tested negative for mycoplasma contamination, checked monthly using the MycoAlert Mycoplasma Detection Kit (Lonza). For maintenance of pluripotency, cells were dissociated to single cells with Accutase (STEMCELL Technologies, 07920) and plated at a density of $1 \times 10^5$ cells per $cm^2$ in Matrigel (BD, 354277)-coated 6-well plates (Corning, 3516) with mTeSR1 (STEMCELL Technologies, 85850) containing 5 μM Y27632, ROCK inhibitor (STEMCELL Technologies, 72302). ROCK inhibitor was removed 24 h after plating (*126*), and cells were cultured for another four days before the next passage. Telencephalic organoids were generated by the directed differentiation protocol as previously described (*34*). Human and macaque iPSCs were dissociated into single cells using Accutase. Neural induction was directed by dual SMAD and WNT inhibition (*127, 128*) using a neural induction medium composited with 50% (v/v) DMEM/F12 (ThermoFisher Scientific, 11330032), 50% (v/v) Neurobasal medium (Thermo Fisher Scientific, 21103049), 1% (v/v) N2 (Thermo Fisher Scientific, 17502048), 2% (v/v) B27 minus vitamin A (Thermo Fisher Scientific, 12587010), 1% (v/v) MEM non-essential amino acid (Thermo Fisher Scientific, 11140050), 1% (v/v) GlutaMAX (Thermo Fisher Scientific, 35050061), 1% (v/v) Penicillin/Streptomycin (Thermo Fisher Scientific 15140122), 0.1 mM 2-Mercaptoethanol (Sigma-Aldrich, M3701), 1 µg/ml heparin (STEMCELL Technologies, 07980). The dissociated cells were reconstituted with the neural induction medium and plated at 10,000 cells per well in a 96-well v-bottom ultra-low-attachment plate (Sumitomo Bakelite, MS-9096V). To increase the cell survival and aggregate formation, 10 µM Y-27632 was added for the first day. For cortical organoids (COs), cells were cultured with the neural induction medium supplemented with 100 nM LDN193189 (STEMCELL Technologies, 72147), 10 µM SB431542 (Sigma-Aldrich, S4317), and 2 µM XAV939 (TOCRIS, 3748) for the first 4 days and then with the medium supplemented with 100 nM LDN193189 and 10 µM SB431542 for the next 4 days. For medial ganglionic eminence organoids (MGEOs), cells were cultured with the neural induction medium supplemented with 100 nM LDN193189, 10 µM SB431542, and 2 µM XAV939 for 8 days. After 8 days, both Cos and MGEOs were transferred to a 6-well ultra-low-attachment plate (Corning, CLS3471) and cultured with organoid growth medium on an orbital shaker (Thermo Fisher Scientific, 88881101) rotating at a speed of 90 rpm to enhance the nutrient and gas exchanges. COs were grown in organoid growth medium composited with 50% (v/v) DMEM/F12, 50% (v/v) Neurobasal medium, 1% (v/v) N2, 2% (v/v) B27 minus vitamin A, 1% (v/v) MEM non-essential amino acid, 1% (v/v) GlutaMAX, 1% (v/v) Penicillin/Streptomycin, 0.1 mM 2-Mercaptoethanol, 2 µg/ml heparin, 2.5 µg/ml human insulin (Sigma-Aldrich, I9278), and 200 ng/ml laminin (Thermo Fisher Scientific, 23017015). For MGEOs, 1X B27 plus vitamin A (Thermo Fisher Scientific, 17504044) was used for the organoid growth medium, and human SHH (R&D System, 1845-SH/CF) and 1 µM purmorphamine (TOCRIS, 4551) were additionally added for ventral specification. From day in vitro (DIV) 21, organoids were cultured with a neuronal maturation medium composited with Neurobasal medium, 1% (v/v) N2, 2% (v/v) B27 plus vitamin A, 1% (v/v) MEM non-essential amino acid,

1% (v/v) GlutaMAX, 1% (v/v) Penicillin/Streptomycin, 0.1 mM 2-Mercaptoethanol, 2 μg/ml heparin, 1% (v/v) Chemically defined lipid concentrate (Thermo Fisher Scientific, 11905031), 10 ng/ml BDNF (R&D System, 248-BD-025; or Peprotech, 450-02), 10 ng/ml GDNF (Peprotech, 450-10), 10 ng/ml NT3 (R&D System, 267-N3-025; or Peprotech, 450-03), 200 μM cAMP (Sigma-Aldrich, D0627) and 200 μM ascorbic acid (Sigma-Aldrich, A92902). For the regionalization of human COs and MGEOs, 100 ng/ml RSPO3 (R&D System, 3500-RS/CF) or 100 ng/ml FGF8b (R&D System, 423-F8/CF) were added to the organoid growth medium from day 8 to day 21. For the GAL and GALP treatment, macaque COs were exposed to 30 ng/ml of human GAL (Phoenix Pharmaceuticals, 026-01), 30 ng/ml human GALP (Phoenix Pharmaceuticals, 026-51), or both from DIV 46 to 60 and collected on DIV 60. Human COs and MGEOs were exposed to 30 ng/ml GALP from DIV 8 to 21 and collected on DIV 35. Brain organoids were collected and fixed in 4% PFA for 24 hours at 4 $^{o}$C. Then, organoids were immersed in step-gradients of sucrose/PBS up to 30% for 2 days at 4 $^{o}$C, embedded in OCT and frozen at -80 $^{o}$C. Sections were prepared at 12 μm on a Leica CM3050S cryostat and stored at -80 $^{o}$C until use.

**Immuno-cytochemistry of the organoids**

Immuno-cytochemistry of the human and monkey telencephalic organoids was started first rehydrating frozen slides in PBS. Blocking was done 1 hour RT in PBS- 10% normal donkey serum (Sigma-Aldrich, D9663) plus 0.5% tween-20. Incubation with primary antibodies was performed in PBS- 10% normal donkey serum plus 0.5% tween-20 at 4 $^{o}$C, ON. The following primary antibodies were used at the concentration indicated: PAX6 (BioLegend, PRB-278P; 1:200); NKX2.1 (ABCAM, ab76013; 1:500); LMX1A (ABCAM, ab76013, 1:200); SP8 (ABCAM, ab73494; 1:200); ZIC4 (Lsbio, LS-B9905-50; 1:100); Galanin (Millipore sigma, AB2233; 1:100); GALR2 (ABCAM, ab188753; 1:100); GALP (Novus biological, NBP2-84950; or Thermo Fisher Scientific, BS-11526R; 1:100); SOX2 (R&D Systems, AF2018; or MAB2018; 1:200); HuC/D (Thermo Fisher Scientific, A-21271; 1:500); KI67 (ABCAM, ab15580; 1:200); GABA (Sigma-Aldrich, A2052; 1:100); CTIP2 (ABCAM, ab18465; 1:1000). Secondary antibody incubation was performed in PBS 1 hour RT. Secondary antibodies were Alexa Fluor 488-, 594-, or 647-conjugated AffiniPure Donkey anti-IgG (1: 200; Jackson ImmunoResearch). Nuclei were counterstained with DAPI (Sigma, D8417). Finally, the slides were mounted using Vector mounting medium.

**Microscopy and imaging**

Fluorescent brain tissue specimens and organoids sections were imaged using a Zeiss LSM800 confocal microscope, or a Zeiss 510 Meta confocal microscope. Z-stack confocal imaging of brain slices and organoids were assembled in Zeiss ZEN2009 and ImageJ (v.2.0.0-rc-69/1.52p) to create image stack projections. Large monkey brain sections were tile scanned using the Zeiss 800 or 510 microscope. Z-stack images of the organoids were analyzed using Volocity (v.6.3.1) and Spotfire (v.11.2.0) software.
